# HOROSCOPE: Decoding human centromere architecture from short reads using *k*-mer signatures

**DOI:** 10.64898/2026.06.10.731283

**Authors:** Carsten Hain, Tobias Rausch, Human Genome Structural Variation Consortium, Human Pangenome Reference Consortium, Jan O. Korbel

## Abstract

Centromeres safeguard genome integrity, yet their roles in human disease remain understudied because of their repetitive sequence. We present HOROSCOPE, a *k*-mer-based framework that infers centromere architecture and generates centromere length estimates directly from short-read data. Using a reference atlas of 11,836 centromeres from telomere-to-telomere haplotypes, we derive diagnostic *k*-mer signatures that classify chromosome-specific architectures with 98.2% precision and 99.1% recall. Applied to 4,029 genomes from 80 populations, HOROSCOPE uncovers continental centromeric structure and African-enriched rare architectures. Across 1,359 cancer genomes, HOROSCOPE reveals a dependency of chromosomal rearrangement locations on the kinetochore attachment site position.

## Background

Centromeres are essential chromosomal sites that direct kinetochore assembly and ensure faithful chromosome segregation during mitosis and meiosis. In humans, centromeres consist of megabase-scale arrays of α-satellite repeats composed of ∼171 bp monomers. Different configurations of monomer subtypes form chromosome-specific higher-order repeat (HOR) arrays, which exhibit extensive internal sequence similarity but differ substantially between chromosomes and among individuals in length, composition, and organization. These central α-satellite arrays are flanked by pericentromeric regions enriched for other repetitive sequences, including inactive HOR arrays, satellite repeats, segmental duplications, and mobile element insertions [1–5]. Functional centromere identity is primarily defined epigenetically by the histone H3 variant CENP-A, which nucleates kinetochore assembly and provides a heritable foundation for centromere activity [5–7]. Active HOR arrays are generally highly methylated, but contain a localized hypomethylated domain, termed the centromere dip region (CDR), that colocalizes with and stabilizes CENP-A-enriched chromatin [8–11]. Together, CENP-A occupancy and the CDR define the functional kinetochore assembly region, which typically represents only a fraction (∼10%) of the active HOR array [4]. Centromeric chromatin is further shaped by flanking pericentromeric heterochromatin enriched for H3K9me3, which is thought to act as a centromere boundary and stabilize CENP-A position [12,13].

Recent telomere-to-telomere (T2T) assemblies have demonstrated extensive population-level diversity in human centromeres, including variation in HOR array length, composition, and organization, as well as mobile element content, kinetochore position, and cases of multiple kinetochore-associated hypomethylated regions within a single centromere. Therefore, centromeres are among the most structurally diverse and rapidly evolving regions of the genome [3,4,14–16].

Despite these advances, the interplay between genetic and epigenetic factors in centromere identity remains largely unresolved. Kinetochores preferentially localize to highly homogeneous, recently expanded regions of HOR arrays [2,4], and α-satellite DNA may support long-term maintenance of CENP-A domains [17]. Yet centromere activity is not strictly dependent on canonical centromeric sequence, as fully functional neocentromeres can form and be maintained at loci lacking underlying α-satellite DNA [18–21].

Due to their central role in chromosome segregation, centromeres are directly relevant to human health and disease [22]. Defects in centromere function, as well as intrinsic fragility, can contribute to chromosomal abnormalities observed in genetic diseases and cancer [22–24]. Cancer genomes are characterized by widespread somatic copy-number alterations (CNA), which result from numerical and structural alterations of chromosomes, including arm-level rearrangements, interstitial losses and gains, and more complex alterations leading to structurally distinct derivative chromosomes, such as isochromosomes [25,26]. Centromeres and pericentromeres have been implicated in chromosome-arm rearrangements; however, the repetitive architecture of centromeric DNA has limited systematic analyses of these regions in larger, typically short-read sequenced, sample cohorts [27–31].

A major gap therefore remains between recent high-resolution centromere assemblies and the much larger public population and cancer cohort sequence datasets available as short-read sequencing data. To address this gap, we developed HOROSCOPE (**H**igher-**O**rder **R**epeat **o**rganization and **S**ize of **C**entromeres using **O**ligonucleotide **P**rofiles for **E**stimation), a readily scalable *k*-mer-based tool for inference of chromosome-specific centromere architecture and HOR array size from short-read data. Applying HOROSCOPE to two large human diversity panels reveals comprehensive insight into the population distribution of centromeric variants around the globe. Furthermore, application of HOROSCOPE to a pan-cancer genomic cohort identifies centromeric HOR truncation patterns in arm-level losses, implying how the CDR position constrains the extent of tolerated centromere breakage.

## Results

### A population-scale dataset of complete human centromeres from near-T2T assemblies

We assembled a comprehensive dataset of completely assembled human centromeres to enable the identification of distinct centromere architectures. Using near T2T assemblies from 562 haplotypes generated across multiple sequencing projects [3,14,32–34], we extracted a total of 11,836 complete human centromere sequences (**Table S1**). Centromeres were identified from 26 human populations with existing near-T2T assemblies by alignment of 200 kb flanking sequences from the annotated centromeres of the CHM13 reference genome to each assembly, and defining the framed region as the centromere plus flanking sequences. To avoid redundancy, we removed duplicate centromere haplotypes arising from overlapping samples across projects or from alternative assembly methods in the HGSVC dataset [14]. Our final dataset comprises a mean of 493 complete centromere haplotypes per chromosome (range: 135 for chromosome Y, which is present only in males and in haploid form, up to 562 for chromosomes 9 and 10) (**Fig. S1**). By incorporating samples representing 26 human populations from five continental ancestries, this dataset captures substantial global human genetic diversity.

### *k*-mer-based clustering of centromeres into distinct architectures

*k*-mer-based strategies have proven useful for annotating repetitive regions in genome assemblies and have shown utility for inference of centromeric haplogroups [35,36]. Building on this general principle, we reasoned that decomposing centromeric sequences into long *k*-mers would enable the identification of distinct centromere architectures for each chromosome and facilitate the extraction of architecture-specific *k*-mers for downstream detection of centromere architectures in short-read datasets (**Fig. 1A**, **Fig. 2A**). We selected *k*-mers of length 61 because they yield a substantially higher fraction of unique *k*-mers in CHM13 (90.34%) than observed for shorter *k*-mers (e.g., 82.37% for 31-mers).

**Figure 1:**
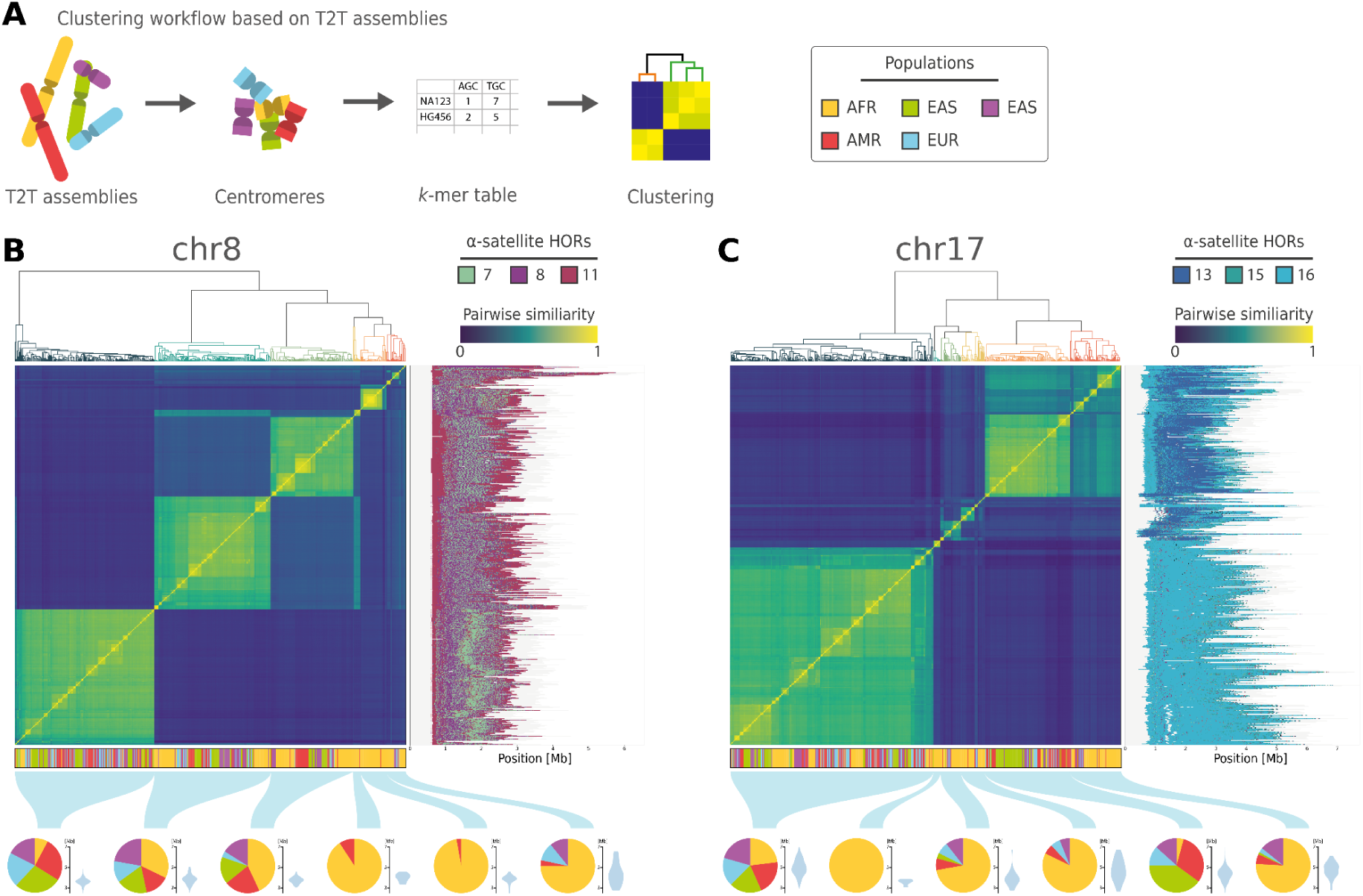
*k*-mer-based clustering of complete human centromeres reveals distinct architectures. (**A**) Workflow for *k*-mer-based centromere clustering. Starting from near T2T assemblies, centromeres with 100 kb flanking regions were extracted and decomposed into 61-mers. Pairwise similarity between haplotypes was calculated based on 61-mer dosages and used for hierarchical clustering. (**B**, **C**) 61-mer based clustering of chromosomes 8 and 17. For each chromosome, the top panel shows the hierarchical clustering, the middle panel displays a matrix of pairwise similarity between haplotypes, and the lower panels indicate the continental population of each haplotype and the distribution of continental population and centromere sizes across clusters. The HOR structure of each haplotype is shown to the right, with HORs colored according to their repeat unit length. Only the most abundant HORs are shown in the legend. Continental population colors follow the 1000 Genomes Project scheme: African (AFR, yellow), American (AMR, red), East Asian (EAS, green), European (EUR, blue), and South Asian (SAS, purple).

**Figure 2:**
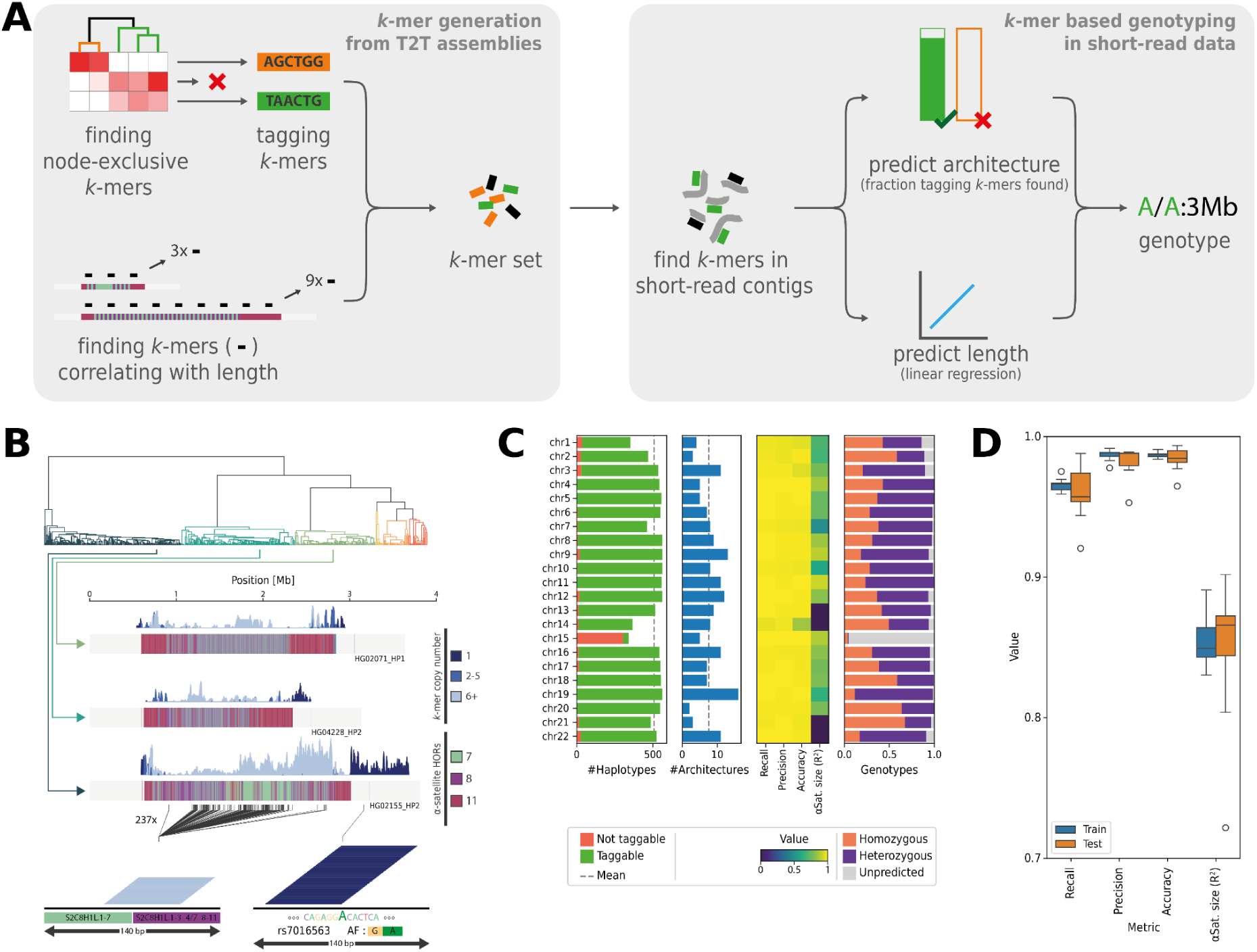
*k*-mer-based characterization of centromere architecture in short-read data. (**A**) Workflow for *k*-mer selection (left) and centromere characterization from short-read data (right). (**B**) Representative chromosome 8 haplotypes selected from major clusters in the hierarchical clustering. The locations of perfect matches of architecture-specific *k*-mers on each haplotype are shown as density plots, with different shades of blue indicating varying levels of *k*-mer repetitiveness. For each haplotype, the HOR structure is displayed, with major HORs indicated in the legend. For haplotype 2 of sample HG02155, two 140 bp excerpts with architecture-specific *k*-mer stacks are highlighted: one marking a junction between the HORs S2C8H1L.1-7 and S2C8H1L.1-3_4/7_8-11 within the α-satellite array, and another tagging the SNP rs7016563 close to the HOR array. (**C**) Summary statistics for centromere architecture characterization from short-read data. From left to right: the number of complete centromeres per chromosome captured by our architecture classification; the number of distinct architectures per chromosome; precision, recall, accuracy, and correlation between predicted and true α-satellite length; and the proportion of samples with homozygous or heterozygous centromere architectures. (**D**) Prediction performance from ten rounds of leave-out cross-validation, showing recall, precision, and accuracy for centromere architecture predictions, as well as correlation coefficients between predicted and true α-satellite length.

We fragmented all extracted centromeric sequences from a single chromosome into all possible 61-mers (**Fig. S2**). The resulting *k*-mer sets were filtered to remove non-repetitive, highly shared and rare 61-mers. Non-repetitive *k*-mers were removed to restrict the analysis to the repetitive portion of the centromere. *k*-mers shared between more than 90% of haplotypes were excluded, as they do not contribute to distinct centromere architectures. Similarly, 61-mers observed in five or fewer centromeres (approximately 1% allele frequency) were removed to focus on more common genetic variation. From the filtered *k*-mer sets, we computed pairwise similarities (see **Methods**) between haplotypes of the same chromosome and used these to cluster haplotypes. The resulting distance-based hierarchical clusterings reveal distinct centromere haplotype clusters, which vary in number, size, population distribution, and, in some cases, in the location of the CDR region, as well as the length of the HOR array (**Fig. 1B, 1C**, **Fig. S3**, **S4**).

Extensive comparison of these clusters shows that haplotypes within the same cluster share a median of 68% of their 61-mers (range: 49%-81%), compared to only 29% (range: 11%-47%) between haplotypes from different clusters. The number and size of clusters vary substantially across chromosomes. Some chromosomes (e.g., chr5, chr6, chr8) exhibit multiple large clusters, whereas others (e.g., chr4, chr17, chr21) are dominated by two major clusters. In addition, each chromosome consistently shows several rare clusters. A notable exception to this pattern is chr15 with no pronounced clustering observed.

We next examined the population distribution of centromere architectures. We find that large clusters are typically population-diverse and contain the majority of non-African haplotypes. In contrast, African haplotypes are significantly more dispersed across architectures when compared to haplotypes from other continental regions (χ² test, *p* < 9.4×10^-5^ for all chromosomes, except chromosome 15). We also find multiple small-to medium-sized clusters that are highly enriched for, or exclusive to, African haplotypes (**Fig. S5**). Haplotypes from other continental populations rarely form such distinct clusters and are instead predominantly found within large, shared architectures, occasionally forming subclusters (e.g., East Asian-enriched subclusters on chr17 or Finnish-enriched clusters on chr10).

In addition to population structure, the *k*-mer-based clustering captures key aspects of centromere organization and HOR structure. Distinct clusters show characteristic HOR composition, structure, and length, with consistent patterns within clusters and marked differences between them. Notable examples include a high-density stretch of HOR S2C8H1L.1-7 in the large, mixed-population cluster of chromosome 8 (**Fig. 1B**), variation in the length of the leading HOR S2C8H1L.1-11 across different chromosome 8 clusters, and the division of chromosome 17 into two distinct clusters – one enriched for S3C17H1L.1-10_15_15-16, and the other dominated by S3C17H1L.1-16 (**Fig. 1C**). In addition to HOR structure, we find that centromere architectures also differ in the positioning of the CDR. Some chromosomes display consistent CDR localization across all architectures (e.g., chr8 with predominantly q-distal CDRs), whereas others (e.g., chr17) show broader variability. In contrast, chromosomes such as chr5 and chr10 exhibit architecture-specific CDR positioning, with distinct architectures associated with either p- or q-distal CDRs (**Fig. S4**).

### Identification of diagnostic *k*-mers for centromere architecture and length

To enable the inference of centromere architecture and length from short-read sequencing data, we aimed to define “diagnostic” *k*-mers for the clustered centromere haplotypes (**Fig. 2A**). We reasoned that both architecture-specific *k*-mers and those *k*-mer whose dosage would reflect centromeric length would be of particular interest. To maximize the number of data points, we utilized all available haplotypes for the identification of such “diagnostic” *k*-mers. To ensure chromosome specificity, we prefiltered *k*-mers by removing *k*-mers present on other chromosomes or in non-centromeric regions of the CHM13 reference genome.

We then queried the hierarchical clusterings and, for each node containing five or more centromere haplotypes, selected all *k*-mers present in at least 98% of haplotypes within the node and in fewer than five haplotypes outside it. After identifying all nodes with architecture-specific *k*-mers, we operationally selected a subset of architectures to meet the following criteria: each haplotype is assigned to a single architecture, and as many haplotypes as possible are captured while a detailed resolution of different architectures is achieved (see **Methods**, **Fig. S6**).

Using this approach, a mean of 93.8% of haplotypes (range: 10.8-100%) could be assigned to a mean of 7.7 architectures per chromosome (range: 2-16) (**Fig. 2C, Table S2**). Consistent with the earlier haplotype clustering, the number and relative sizes of centromeric architectures vary substantially across chromosomes. A cumulative analysis of centromere architectures across the assembled haplotypes shows that the current set of completely assembled centromeres has not yet reached saturation, with rare architectures continuing to accumulate as additional haplotypes are added, particularly after inclusion of African haplotypes (**Fig. S7**).

The resulting architecture-specific *k*-mer sets span a wide range of copy numbers (**Fig. S8**), but are predominantly composed of low-copy *k*-mers, with most appearing only once. These *k*-mers primarily originate from small sequence variants located outside the HOR array (**Fig. 2B**). In contrast, a smaller subset of highly repetitive *k*-mers derives from specific monomers or monomer combinations within the HOR array and shows localized enrichment patterns characteristic of distinct centromere architectures (**Fig. 2B**).

In addition to architecture-specific *k*-mers, we set out to identify *k*-mers whose dosage correlates with the length of the α-satellite array (**Fig. 2A**). This analysis yields a mean of 2,646 (range: 414-5,312) length-informative *k*-mers per chromosome. Notably, this strategy identifies *k*-mers predictive of centromere length independently of the haplotype architecture, such that in diploid samples their aggregate dosage reflects the mean length of both centromere alleles.

Because length estimation relies on *k*-mer dosage, which scales with sequencing depth, we selected chromosome-arm-specific normalization *k*-mers. These *k*-mers occur exactly once per haplotype and are located as close as possible to the centromere array, allowing normalization across samples with varying sequencing coverage. This strategy reveals sufficient numbers of normalization *k*-mers for both chromosomal arms for 17 of 22 autosomes, whereas we find substantially lower numbers of normalization *k*-mers for the q arm of chromosomes 2 and 12 and the acrocentric chromosomes 13, 14 and 21 (**Fig. S9**).

### Performance of *k*-mer-based centromere characterization from short-read sequencing data

We evaluated the performance of our *k*-mer-based centromere analysis using a benchmark dataset consisting of 272 samples from the 1000 Genomes Project for which both short-read sequencing data and corresponding T2T genome assemblies are available [14,32,34,37,38]. Short-read sequencing data from these samples were represented as de Bruijn graphs (DBGs) (see **Methods**) using a *k*-mer size of 61. This representation reduces sequencing errors and preserves *k*-mer dosage information comparably to raw sequencing data (**Fig. S10**).

To assess centromere architecture inference, architecture-specific *k*-mers and their dosages were queried in the DBG representations. For each architecture, we calculated the fraction of architecture-specific *k*-mers detected and considered architectures with more than 50% of their architecture-specific *k*-mers present as detected (**Fig. 2A**). Comparison of short-read-based predictions with architecture assignments derived from T2T assemblies demonstrates high precision (mean 98.2%, range: 85.9%-100%) and recall (99.1%, range: 97.1%-100%) across chromosomes (**Fig. 2C**). Individual chromosomes display varying degrees of heterozygosity of centromere architectures, which correlate with the number of distinct architectures observed per chromosome (**Fig. S11A**). Samples of African ancestry exhibit higher heterozygosity than other continental populations, reflecting their greater contribution to rare centromere architectures (**Fig. S11B**).

We next evaluated centromere length prediction using length-informative *k*-mers. We normalized the *k*-mer dosage for sequencing coverage and corrected for collinearity using principal component analysis. The resulting data were fitted to α-satellite array length derived from T2T assemblies. This approach yields strong correlations between *k*-mer-dosage and α-satellite length (mean R^2^=0.75, range: 0.49-0.92) (**Fig. 2C**). We applied this approach to all centromeres except those on acrocentric chromosomes 13, 14, 21, and 22. In these cases, normalization *k*-mers lack unimodal dosage distributions (**Fig. S12**), indicating elevated copy numbers in variable regions for a subset of *k*-mers. This compromises coverage normalization, leading to their exclusion from length prediction analyses.

To evaluate the robustness of our method, we performed ten rounds of leave-out cross-validation on chromosome 8. In each round, 20% of samples with two complete haplotypes were randomly selected as the test set. To avoid circularity, we employed alignment-based clustering for this benchmarking rather than *k*-mer-based clustering. The two clustering approaches produced highly similar results (**Fig. S13**), with only 25 of 561 haplotypes assigned to different architectures. We pursued the complete *k*-mer-based centromere analysis procedure, including the identification of architecture- and length-informative *k*-mers and the linear model for α-satellite length, on the training set and subsequently applied it to the test set. The predictions generated, notably, show strong concordance with T2T-derived ground truth (**Fig. 2D**), yielding accuracy, precision, and recall comparable to those obtained using all haplotypes, and accurately recovering centromere length. While classification performance remains stable across cross-validation rounds, estimates of α-satellite length show moderate variability, reflecting the influence of structurally atypical centromeres on model training. To facilitate application of this approach, we implemented this framework into HOROSCOPE, a streamlined workflow for centromere characterization from short-read sequencing data (see **Methods**).

### Centromere architecture across 80 human populations mirrors continental genetic structure

To investigate population-level variation in centromere architecture, we applied HOROSCOPE to a collection of 4,029 short read-sequenced genomes, comprising samples sequenced in the 1000 Genomes Project [37,38] (N=3,201 samples) as well as the Human Genome Diversity Project [39] (N=828 samples). This combined set of genomes spans 80 populations (**Fig. S14**) with a median of 22 individuals per population (range 2–179; median for 1000 Genomes Project samples: 115; **Fig. S15**).

Using allele frequencies of centromere architectures across populations, multiple factor analysis (MFA) reveals a global population structure (**Fig. 3A**) that closely mirrors patterns observed for other forms of genetic variation such as SNPs [40] and structural variants [41]. We next quantified centromere architecture diversity across populations, observing marked differences between continental groups, with African populations showing the highest and Oceanian populations the lowest diversity (**Fig. 3B**).

**Figure 3:**
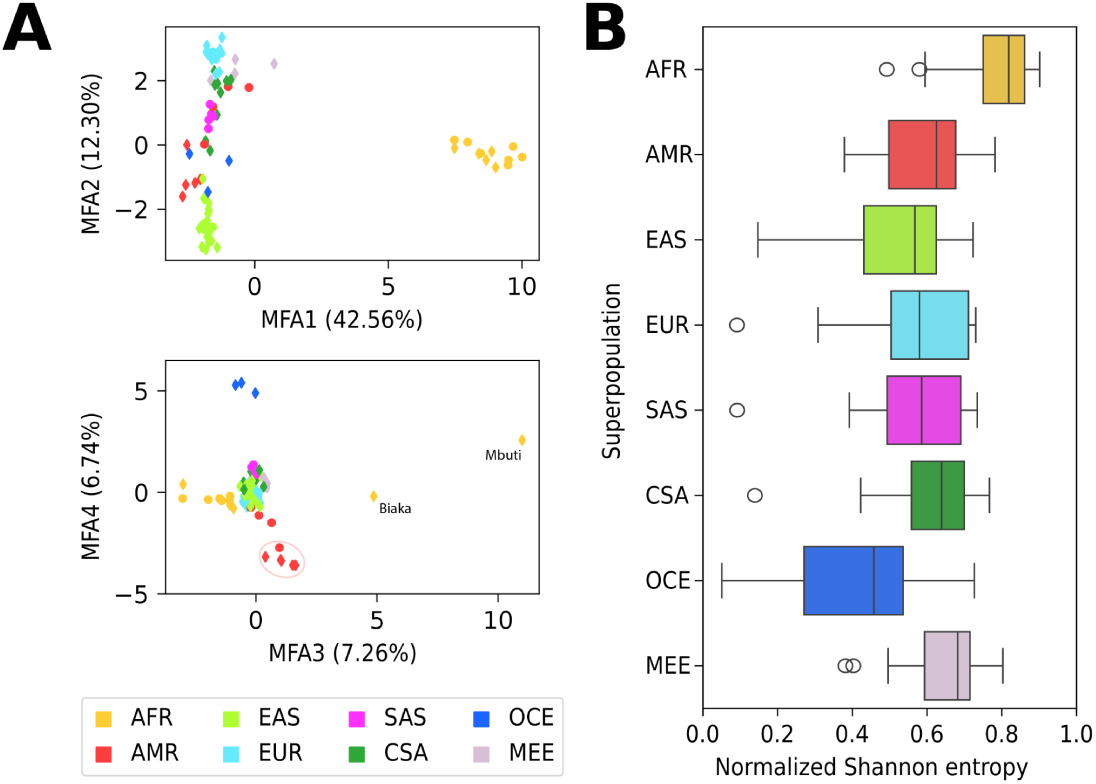
Population structure and diversity inferred from centromere architecture frequencies. (**A**) Multiple factor analysis (MFA) of centromere architecture allele frequencies across 80 populations from the 1000 Genomes Project and the Human Genome Diversity Project. The first and second, as well as the third and fourth, MFA dimensions are shown; each point represents one population. Circles denote populations from the 1000 Genomes Project, and diamonds denote populations from the Human Genome Diversity Project. In the lower panel, markers denoting the Biaka and Mbuti populations are labeled, and populations of American ancestry with a high proportion of Native American ancestry are circled. (**B**) Distribution of centromere architecture diversity across continental groups, measured using normalized Shannon entropy. For each continental population, the boxplot shows chromosome-wise diversity values, with one value per chromosome calculated from centromere architecture frequencies within that continental population. Populations are grouped into continental population: African (AFR), American (AMR), East Asian (EAS), European (EUR), South Asian (SAS), Central/South Asian (CSA), Oceanian (OCE), and Middle Eastern (MEE).

Within MFA space, Eurasian populations form a continuous gradient that roughly follows geographic relationships, extending from European (EUR) through Central and South Asian (CSA, SAS) to East Asian (EAS) populations. Middle Eastern samples occupy an intermediate position between EUR and CSA populations, with the Mozabite population showing increased diversity (**Fig. S16**) and a closer similarity to African (AFR) populations, consistent with previously reported admixture signals based on SNPs [42]. We further find that Oceanian (OCE) and most American (AMR) populations separate from Eurasian populations along higher MFA dimensions (**Fig. 3A**). OCE populations exhibit the lowest centromere diversity among all continental groups. AMR populations show broader dispersion across Native American (NAM) admixture levels. Populations with low-to-medium NAM (MXL, PUR and CLM) show a centromere diversity comparable to other continental groups, whereas high-NAM populations (PEL and HGDP populations) [43] exhibit reduced diversity. (**Fig. S16**).

Consistent with observations made from other classes of genetic variation, AFR populations exhibit the highest overall diversity of centromere architectures [37]. Notably, the Mbuti and Biaka are separated from other AFR populations in MFA space and appear to show reduced centromere diversity. However, this pattern is notably accompanied by a 14- to 24-fold higher frequency of missing architecture predictions across chromosomes compared with other AFR populations (**Fig. S17**). Missing predictions in the Mbuti are most frequent on chromosomes 3 (7 of 10 individuals), 7 (4 of 10), and 8 (3 of 10), whereas in the Biaka, these occur most often on chromosomes 1 (8 of 24) and 12 (5 of 24). These missing calls are likely to reflect centromere architectures underrepresented or absent in other global populations, suggesting that the Biaka and Mbuti harbour additional centromeric diversity not yet captured in current T2T assemblies.

### Centromere alterations and structural variation in cancer genomes

We next explored the centromeric landscape in cancer genomes sequenced with short reads, thereby focusing on data from the Pan-Cancer Analysis of Whole Genomes Consortium (PCAWG) [44] (N=2,642 samples).

To facilitate centromeric analysis, we excluded samples with evidence of whole-genome duplication as defined by the original studies, as well as samples exhibiting systematic shifts in *k*-mer dosage between tumor and matched normal in genomic regions expected to be copy-number neutral (see **Methods**). After filtering, 1,359 tumors across 35 cancer histologies were retained for downstream analysis, corresponding to 69.6% of all non-whole-genome duplicated samples in the combined cohorts. To these samples, we applied HOROSCOPE to predict centromere architecture and length for each tumor-normal pair across the cohort.

We first asked whether centromere features were associated with chromosomal instability [25]. To this end, we performed logistic regression incorporating germline centromere length and architecture together with relevant clinical and genomic variables, including cancer type, *TP53* mutation status, mutational burden, age, ancestry, and sex. While the model demonstrates good discriminatory performance (AUROC = 0.76), we find that centromere features show no significant association with chromosomal instability within the resolution and statistical power of our analysis. Instead, the strongest predictors are *TP53* mutation status, overall mutational burden, and tumor histology (**Fig. S18**).

We therefore focused on somatic chromosome-scale alterations with centromeric breakpoints to ask whether centromere organization influences the structure of individual CNA events rather than affecting overall chromosomal instability burden. Chromosome-scale alterations were then classified based on CNA profiles into whole-chromosome aneuploidies with modal copy numbers of 1 (n=775), 3 (n=664) and 4 (n=61) as well as arm losses (n=471), arm gains (n=111), and isochromosomes (n=96). Our assignment criteria included concordant CNA patterns, minimal additional broad CNA alterations, and copy number states near the centromere consistent with centromeric breakpoints.

Based on our centromere size predictions, we estimated, for each chromosome, the number of centromeres per clonal tumor cell by integrating centromere length predictions from the tumor and matched normal with tumor cell fraction (**Fig. 4A, 4B**). We find that whole-chromosome aneuploidies exhibit discrete, ladder-like distributions, with centromere number scaling proportionally with copy number. In contrast, arm-level alterations show broader and asymmetric distributions, spanning configurations from near-diploid states to cases with one to three centromeres per cell. We also observe intermediate, non-integer centromere counts, consistent with partial truncation of centromeric arrays. Isochromosomes show symmetric but wider distributions. Together, these patterns are consistent with substantial variability in breakpoint location within the centromere, leading to either truncated or extended centromeric arrays.

**Figure 4:**
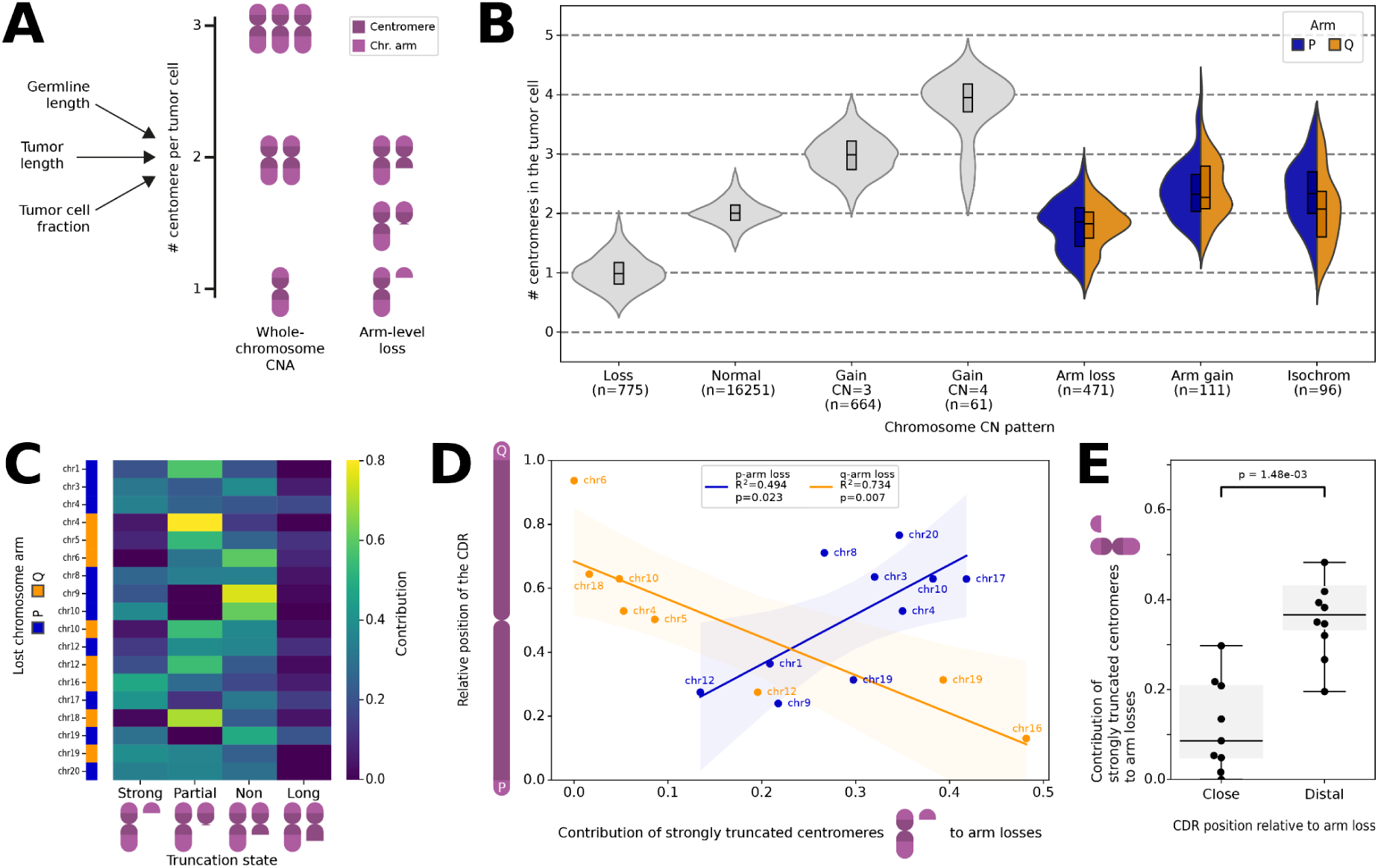
Landscape of centromeres in cancer. (**A**) Schematic overview of the approach used to estimate the number of centromeres per tumor cell, based on predicted HOR array lengths in matched germline and tumor samples and the tumor cell fraction (left). Representative centromere configurations in the genomes of tumor cells are depicted, illustrating whole-chromosome CNAs and arm-level losses with varying degrees of centromere truncation (right). Centromeres are shown in dark purple, chromosomal arms in light purple. (**B**) Number of centromeres per tumor cell across different chromosomal configurations. CNAs affecting the whole chromosome are shown in gray, while arm-level CNAs spanning from telomere to centromere are stratified into those that affect either the p arm (cyan) or q arm (gold). The number of events for each class is stated below. (**C**) The distribution of centromere lengths in arm-loss CNAs was deconvoluted into events resulting in strong or partial truncation, apparent elongation, or no change in centromere length. The contribution of each state to losses of different chromosomal arms is shown as a heatmap. The y-axis lists affected chromosomes, and the lost arm is indicated by color (cyan, p arm; gold, q arm). (**D**) Correlation between the fraction of strongly truncated centromeres in arm-loss events and the relative CDR position inside the centromere separately for p- and q-arm losses. Regression fits with confidence intervals, R², and p-values are shown. **(E)** Fraction of highly truncated centromeres stratified by the relative position of the CDR with respect to the lost chromosomal arm (two-sided Mann-Whitney U test). Events are grouped based on the mean CDR position within the centromere, distinguishing cases where the CDR lies in the centromeric half close to the lost arm versus the half distal to the lost arm.

We aimed to investigate the distribution of centromeric breakpoints and the resulting truncation patterns of centromeric arrays in arm-loss events in greater detail (**Fig. S19)**. To this end, we applied Gaussian mixture modeling to the inferred distribution of centromere number per tumor cell for each chromosomal arm separately. This approach enabled estimation of the fraction of arm-loss events showing strong truncation, partial truncation, no detectable truncation, or apparent elongation of the centromeric array (**Fig. 4C**). Across all arm losses, centromeric arrays show comparable fractions of strong truncation (30%), partial truncation (29%), or no detectable loss (31%), while a smaller subset (10%) exhibits apparent elongation. Interestingly, these patterns differ markedly between short (p) and long (q) chromosomal arms: losses on p arms more frequently lead to strong truncation (34%, compared to 23% for q arms), whereas losses on q arms more often result in partial truncation (46%, 21% for p arms). The proportion of events with no detectable centromeric truncation is lower in p- than q-arm losses (26% and 33%, respectively). We also note chromosome-specific differences, for example, p-arm losses on chromosome 17 more frequently result in strong (42%) instead of partial truncations (11%), whereas chromosome 12 shows the opposite pattern, with partial truncations (39%) occurring more frequently than strong truncations (13%).

These differences in truncation patterns prompted us to investigate whether breakpoint location within the centromere is influenced by its internal organization. Given that centromere function is localized to the CDR, which marks the kinetochore attachment site, we examined the relationship between CDR position and centromere truncation. The proportion of strongly truncated centromeres following arm loss correlates with the position of the CDR (**Fig. 4D, Fig. S4**), showing a positive relationship for p-arm losses (R² = 0.494, *p* = 0.023) and a negative relationship for q-arm losses (R² = 0.734, *p* = 0.007). When aggregating p- and q-arm losses, events in which the CDR is located in the centromeric half distal to the lost arm show a 3.0-fold higher proportion of highly truncated centromeres (*p* = 1.48×10^-3^, Mann-Whitney U test) (**Fig. 4E, Table S3**). This trend is retained after stratifying events by *TP53* mutation status, suggesting that the association between CDR position and centromere truncation is not primarily driven by *TP53*-associated genome instability (**Fig. S20**). Together, these results indicate that centromeres with a CDR positioned close to the lost arm tend to exhibit fewer strongly truncated α-satellite arrays than those in which the CDR is located at intermediate or distal positions.

## Discussion

Centromeres have long remained inaccessible to population-scale analysis due to their highly repetitive sequence. Recent advances in long-read genome assembly have begun to overcome this limitation, revealing extensive structural and epigenetic diversity across human centromeres [14,15,33,45,46]. However, extending centromere genomics to population-scale analyses and leveraging the vast collections of existing short-read sequencing data remains a major challenge.

By leveraging complete centromere assemblies to define diagnostic *k*-mers and quantifying their presence and dosage in short-read data, HOROSCOPE infers key features of centromere organization including length information at scale. *k*-mer-based approaches have recently been used both to annotate repetitive elements in complete genome assemblies and to infer centromeric haplogroups from short-read data using rare centromeric *k*-mers, highlighting the potential of alignment-free sequence representations for repetitive genomic regions [35,36]. In contrast, HOROSCOPE uses both presence and dosage of long *k*-mers to jointly infer centromere organization and size across large short-read cohorts and performs robustly in a subset of the 1000 Genomes Project with matched ground truth. Despite this performance, we emphasise that some limitations remain. Inference is less reliable for acrocentric chromosomes, and the approach is sensitive to sequencing protocols and data quality, necessitating stringent filtering of heterogeneous datasets. Furthermore, our approach does not resolve haplotype-specific centromere length and does not capture complex rearrangements, translocations, and epigenetic variation, which necessitate long-read sequencing.

We derived the *k*-mers central to HOROSCOPE primarily from complete T2T assemblies generated by the HGSVC and HPRC consortia. With continued improvements in long-read methodology [47,48], and with ongoing efforts by these and other groups, the number and diversity of complete human T2T genomes are expected to increase substantially. In this context, HOROSCOPE and the underlying principle of T2T assembly-derived *k*-mers represent an extensible framework for the genomics community.

At the same time, the number of short-read-sequenced human genomes already reaches into the millions [49–51], far exceeding the number of available long-read or T2T assemblies. HOROSCOPE therefore provides a scalable bridge between the resolution of complete genome assemblies and the scale of short-read sequencing cohorts. Applying HOROSCOPE to these large datasets could enable population- and disease-scale analyses of centromere variation, increasing statistical power to identify associations between repeat structure, ancestry, genome instability, and human disease.

Applying HOROSCOPE to globally diverse cohorts reveals that centromere architecture frequencies recapitulate continental population structure, closely mirroring patterns observed for genome-wide genetic variation [40]. This suggests that centromeres are shaped by the same demographic and evolutionary processes that govern variation across the rest of the genome. However, the continued accumulation of rare architectures in newly assembled haplotypes, together with elevated rates of missing predictions in Mbuti and Biaka samples, suggests that current telomere-to-telomere genome assemblies still incompletely capture global human centromere diversity.

Motivated by the observation that many CNAs in cancer originate near centromeric regions [30,31,52], we further explored the analysis of pan-cancer cohorts using HOROSCOPE. We did not find a significant association between centromere architecture or α-satellite array length and hallmarks of chromosomal instability. This suggests an inherent stability across the diverse centromere structures, at least given the limitations of our method and used dataset. Yet, we identify a correlation between the location of the CDR and the location of breakpoints. In arm-level losses, we find a general prevalence of centromeres with little to no truncation and we find that the position of the CDR correlates with the prevalence of strongly truncated centromeres. Centromeres whose CDR lie close to the lost arm exhibit fewer strongly truncated α-satellite arrays than those with more distal CDR positions.

While loss of α-satellite DNA itself may progressively compromise centromere function, for example by reducing levels of CENP-A or α-satellite-derived transcripts [53–55], the CDR-proximity-dependent truncation pattern may reflect two non-mutually exclusive mechanisms. First, chromosomes that lose the CDR itself would be predicted to be subject to negative selection and might be preferentially lost rather than maintained as truncated derivative chromosomes. This might explain why extensive α-satellite loss is more frequently observed when the CDR lies distal to the lost arm, allowing large truncations while preserving the kinetochore-associated region. Second, breakpoints may be enriched at or near the CDR. This possibility is consistent with elevated sequence variation within CDRs in population-scale centromere assemblies [4] and structural and genetic variation at the CDR in the genome of a melanoma cell line [56]. In addition, we note that the apparent dependence of cancer rearrangement breakpoint positions on CDR positions determined from lymphoblastoid cell lines suggests that CDR positioning within centromeric architecture is broadly conserved across cancer tissues. This interpretation extends recent findings, reported in a preprint [57], on the stability of CDR location across non-malignant cell-fate transitions.

Irrespective of the actual mechanism, centromere disruption could lead to a range of downstream configurations [22], depending on whether residual centromeric sequence remains functional, whether centromere activity is re-established as a neocentromere, or whether broken chromosomal fragments are stabilized through additional rearrangements. How these outcomes relate to the original centromere organization and the precise breakpoint location remains unknown. These structures cannot be resolved with short-read data alone, but recent long-read-based cancer genomes resolved centromeric breakpoints in cancer cell lines [56,58]. Finally, the observed elongation of α-satellite arrays following centromere breakage suggests the involvement of repair-associated mechanisms, consistent with experimental evidence that human centromeric arrays can expand in cells [59]. Resolving these structures at the basepair level will require high-resolution analyses using long-read based genomic assemblies in extended human sample cohorts.

## Conclusions

HOROSCOPE infers centromere architectures and generates length estimates from widely available short-read data, enabling centromere analysis across large existing disease and diversity cohorts. Applied to thousands of genomes, HOROSCOPE resolves centromeric structure across human populations, suggests that the diversity of human centromeres remains incompletely catalogued and implies that CDR position constrains centromere-associated rearrangement patterns in cancer. As an open-source Nextflow pipeline, HOROSCOPE enables population-scale study of these previously inaccessible genomic regions.

## Methods

### Extraction of centromeric regions from near T2T assemblies

Centromeres and their flanking regions were identified by aligning manually defined centromere-flanking regions to near T2T assemblies from multiple genome projects.

Flanking regions were selected on the CHM13 reference genome using the UCSC Genome Browser CenSat annotation track. Specifically, 200 kb regions up- and downstream of each centromeric HOR array were chosen to overlap predominantly unique, non-repetitive sequences. Coordinates for these regions are listed in **Table S3**. Flank sequences were extracted from CHM13 using bedtools v.2.31.1 getfasta [60]. To select centromeres from T2T assemblies, the flanks were aligned using minimap2 v.2.30 (parameter -x asm20 -secondary=no) [61] and regions were extracted where both flanks of a chromosome align to the same contig in the same orientation. The resulting extracted region included both flanks and the centromere in between. All extracted centromeres were standardized to the same orientation by aligning them to CHM13. If the primary alignment was in reverse orientation, the reverse complement of the sequence was used. For all extracted centromeres, additional information like contig length, associated continental population of the haplotype and genome project were gathered in one table (**Table S1**). To remove duplicate haplotypes, the following criteria were applied: when more than two complete centromeric haplotypes were extracted per sample, two haplotypes were retained preferentially from the same project and assembler. If this was not possible, the two haplotypes with the greatest length difference (at least ≥1 kb) were selected. The haplotypes retained for downstream analyses are indicated in Table S1.

### Higher-order repeat annotation

Higher-order repeats (HOR) in centromeres were annotated using humAS-HMMER (https://github.com/fedorrik/HumAS-HMMER_for_AnVIL) to detect individual α-satellite monomers. Monomer annotations were then grouped into HORs using StV (https://github.com/fedorrik/stv). For visualizations of centromeric HOR structure, each HOR unit was colored according to the number of α-satellite monomers it contains. From the humAS-HMMER annotations the combined size of all live α-satellite monomers (indicated by a L in their annotations) was calculated for each haplotype.

### Generation of 61-mer count tables from centromeres

Single centromere haplotypes were decomposed into 61-mer counts using dicey v.0.2.8 [62] (parameter chop --length 61 --se) in forward and reverse mode. Forward and reverse 61-mers were merged, sorted and counted to generate a complete set of all 61-mers and their associated counts for each haplotype. For each chromosome, haplotype-level 61-mer counts were merged by first aggregating all 61-mers present across haplotypes, and then querying the count of each 61-mer in each haplotype. This process resulted in one count matrix per chromosome, with unique 61-mers as rows and individual haplotypes as columns.

### Pairwise similarity of centromeres based on 61-mers

The similarity between two centromeres was assessed based on the produced *k*-mer count tables. To prepare the *k*-mer tables for this analysis, 61-mers that are shared, rare or non-repeating were removed. 61-mers were defined as shared if they occur in more than 90% of all haplotypes, they were defined as rare if they occur in 5 or fewer haplotypes and they were defined as non-repeating if their maximum copy-number in any haplotype is less than 3. Based on the filtered 61-mers, the pairwise similarity between haplotypes A and B is calculated as:

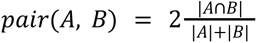

where |A∩B| is the number of 61-mers shared by both haplotypes, and |A| or |B| are the total number of 61-mers present in haplotype A or B, respectively.

### Clustering of haplotypes based on pairwise similarity

From a pairwise similarity matrix, clustering was performed using the linkage module of scipy v1.12.0 with the weighted method and a clustering cutoff of 3.15.

### Analysis of population distribution in centromere architectures

From the assignments of haplotypes to distinct centromere architectures in the hierarchical clustering the number of haplotypes from each continental population in each architecture was computed. A different distribution of African samples, compared to non-African samples, was analyzed using Chi-square test of independence of variables in a contingency table using the chi2_contingency module of scipy v1.12.0.

### Selection of chromosome and centromere specific *k*-mers

*k*-mer sets from all centromeres on a given chromosome may include *k*-mers that are either present elsewhere in the genome or shared with centromeres from other chromosomes. To select *k*-mers specific to the centromeric region of a single chromosome, we applied a two-step filtering process. First, all *k*-mers found in the *k*-mer sets of other chromosomes were removed. The remaining *k*-mers were then mapped to the CHM13 reference genome, in which the centromeric regions - defined by the flanking coordinates used for initial centromere extraction - had been masked. *k*-mers that aligned with an edit distance of zero (NM=0) were excluded. The remaining *k*-mers were considered centromere- and chromosome-specific.

### Selecting architecture-specific *k*-mers

Based on the centromere- and chromosome-specific *k*-mer sets and the hierarchical clustering performed for each chromosome, we selected architecture-specific *k*-mers. To avoid inflation of rare architectures driven by minimal differences, only nodes in the hierarchical clustering containing five or more haplotypes were considered. For all nodes meeting this criterion, node-specific *k*-mers were extracted. For this analysis all centromere-and chromosome-specific *k*-mers were used, including those with a repeat count of one.

A *k*-mer was considered specific to a node if it occurred in at least 98% of the haplotypes within that node and in five or fewer haplotypes outside the node. The cutoff of five haplotypes outside the node was chosen to represent approximately 1% of all haplotypes and was reduced to one for small nodes containing 10 or fewer haplotypes.

To manage very large *k*-mer sets from individual nodes and improve usability in downstream analyses, *k*-mers with the lowest repeat counts were randomly removed until each *k*-mer set contained at most 20,000 *k*-mers. Because individual nodes within the hierarchical clustering correspond to distinct centromere architectures, the *k*-mers selected through this process were designated architecture-specific *k*-mers.

### Selection of centromere architecture nodes for downstream inference

Selection of architecture-specific *k*-mers yielded specific *k*-mers for nodes at different levels of the hierarchical clustering tree. For downstream analyses, we sought to reduce this full set of nodes with specific *k*-mers to a non-overlapping and interpretable set of centromere architectures that still captures the structural diversity observed for each chromosome.

To define this subset, nodes were selected manually following three criteria.

1. In the subset, each haplotype is assigned only to a single architecture.
2. The nodes selected for the subset were chosen to maximize the number of haplotypes represented by architectures with architecture-specific *k*-mers.
3. The subset preserves detailed resolution of the distinct centromere structures observed for each chromosome.

The second and third criteria represent a trade-off between population coverage and architectural resolution. We therefore prioritized the largest possible number of distinct architectures while retaining more than 95% of haplotypes within groups with architecture-specific *k*-mers whenever possible.

### Selection of length-informative *k*-mers

To infer HOR array size from short-read data, we selected *k*-mers whose dosage correlates with α-satellite array size for each chromosome. Candidate *k*-mers were drawn from the centromere- and chromosome-specific *k*-mer sets and required to have a maximum copy number greater than 20 across haplotypes. For each candidate *k*-mer, we calculated the Pearson correlation coefficient between *k*-mer copy number and HOR size. HOR size was derived from the humAS annotations as the summed length of all active (L) HOR calls. *k*-mers with a correlation coefficient greater than 0.75 were retained as length-informative *k*-mers.

### Selection of normalization *k*-mers

Normalization *k*-mers were obtained from the centromere- and chromosome-specific *k*-mer set by selecting *k*-mers with a copy number of one across all centromere haplotypes for a given chromosome. These *k*-mers were mapped to the CHM13 centromere contig using BWA-MEM v.0.7.18 [63], retaining only exact matches (edit distance NM = 0). As the majority of these *k*-mers originated from the non-repetitive regions flanking the centromeric array on the p- and q-arm sides, positional clustering was performed using K-means (k = 2) from scikit-learn v1.5.1. The cluster with the lower mean genomic position was annotated as the p-arm flank, and the cluster with the higher mean position as the q-arm flank. If possible, 5,000 *k*-mers were randomly sampled from each side and formed the final normalization *k*-mer set.

### De Bruijn graph from short read data

Raw short read data were transformed into the format of a de Bruijn graph with *k*-mer size 61. To calculate such a graph, raw short reads were first error-corrected with Lighter v.1.1.3 [64] (parameter -trim -discard -k 23 3100000000 0.188) to reduce sequencing-error-derived *k*-mers. Corrected reads were assembled with BCALM2 v.2.2.3 [65] (parameter -kmer-size 61 -abundance-min 3). The resulting de Bruijn graph consists of unitig sequences representing paths of overlapping 61-mers, with *k*-mer abundance retained as the mean unitig coverage in the km field of each FASTA header. This km value was used as the dosage estimate for queried *k*-mers.

### Finding *k*-mer presence and dosage in BCALM2 de-Bruijn graphs

*k*-mer presence and dosage were extracted from short-read de Bruijn graphs. For each query *k*-mer, both the forward sequence and its reverse complement were searched against the graph unitigs using grep. The -B 1 option was used to retrieve both the matching unitig and its FASTA header, which contains the mean *k*-mer abundance for that unitig. *k*-mer dosage was then extracted from the km field of the unitig header. If neither the query *k*-mer nor its reverse complement was detected in the graph, its dosage was set to zero.

### Training centromere length prediction models

To enable prediction of centromere and α-satellite lengths from short-read sequencing data, we trained linear regression models based on *k*-mer dosage. These models utilize length-informative *k*-mers (selected as described above) that correlate with centromere length independently of specific centromere architectures, allowing estimation of the mean centromere length across both haplotypes. The models were trained using 272 samples from the 1000 Genomes Project for which both short-read data and fully assembled centromeres for both haplotypes of a given chromosome were available. For each of these samples, the mean HOR lengths across haplotypes were calculated. The dosage of each length *k*-mer was extracted from the sample’s de Bruijn graph and normalized for varying coverage by dividing by the median dosage of the chromosome-matched set of normalization *k*-mers. Normalized length-informative *k*-mers were correlated with HOR array size and *k*-mers with a Pearson correlation coefficient below 0.7 were excluded. The dosage of the remaining *k*-mers were reduced to two dimensions using principal component analysis (PCA). Linear regression models were then trained on the PCA-transformed data to predict HOR array size. All trained models and their corresponding *k*-mer sets were saved for downstream prediction tasks.

### Centromere prediction from short-read data with HOROSCOPE

We developed HOROSCOPE (https://github.com/carstenhain/HOROSCOPE/), a workflow to infer centromere architecture and HOR array length from short-read sequencing data. The method is based on quantifying the dosage of predefined *k*-mer sets, including architecture-specific *k*-mers, length-informative *k*-mers, and normalization *k*-mers (see above). These dosage profiles are then used to infer centromere architecture and estimate HOR array length.

As a first step, *k*-mer dosages are extracted from short-read data. Short-read data can be input as a de Bruijn graph or as raw sequencing data. For de Bruijn graph input, *k*-mer dosages are extracted as described above. For raw sequencing data, *k*-mers are counted using Jellyfish [66] with the parameters count -m 61 -s 2G -C, and the dosages of the predefined *k*-mer sets are extracted using the Jellyfish query command.

The resulting *k-*mer dosages are normalized to account for differences in sequencing coverage. Because normalization is a key parameter for length prediction, and because cancer genomes may contain copy-number alterations affecting centromeric regions, the workflow supports several normalization strategies. First, normalization can be performed using the mean dosage of all normalization *k*-mers across chromosomes. Second, normalization can be restricted to using only normalization *k*-mers from the chromosome being analyzed. Third, a predefined normalization value can be supplied, forcing the workflow to use this value for normalization. After selection of a normalization value, normalized *k*-mer dosages are obtained by dividing each *k*-mer dosage by the selected normalization value.

Centromere architecture is inferred from the normalized dosages of architecture-specific *k*-mers. For each architecture, we calculated the fraction of architecture-specific *k*-mers with a normalized dosage greater than 0.1. If more than 50% of the architecture-specific k-mers for a given architecture exceeded this threshold, the sample was inferred to contain at least one haplotype with that architecture.

HOR array length is predicted separately for each chromosome using length-informative *k*-mers. The normalized dosages of these *k-*mers are projected onto a pretrained principal component analysis model, followed by linear regression to estimate HOR array length.

The complete procedure was implemented as a single Nextflow workflow, enabling prediction of centromere architecture and HOR array length directly from raw or assembled short-read data. In addition to centromere feature predictions, the workflow reports quality-control metrics for each chromosome, including the mean dosage, standard deviation, and number of detected p- and q-arm normalization *k*-mers. These outputs provide direct metrics for assessing sequencing quality and the reliability of normalization. Because we observed the best performance when using de Bruijn graph representations as input, the workflow also includes an option to convert raw sequencing data into a de Bruijn graph prior to centromere inference.

### Evaluation of centromere architecture and length prediction

To assess the accuracy of centromere architecture and length prediction, we analyzed samples from the 1000 Genomes Project for which both short-read sequencing data and corresponding T2T assemblies were available. For each sample, the ground truth centromere characteristics - specifically the centromere architecture of each haplotype and the mean centromere length - were extracted from the centromere assemblies. Predicted architectures and lengths were then compared to these ground truth values. Prediction performance for centromere architecture was quantified using precision, recall, and accuracy. For centromere length prediction, we calculated the coefficient of determination (R^2^) between predicted and true values.

### Pairwise similarity of centromeres based on MUMmer4 alignment

Pairs of centromere sequences were aligned using the MUMmer4 v.4.0.0 suite [67]. The alignment was generated with nucmer, followed by processing of the delta file using delta-filter (parameter -1). The filtered alignment was analyzed with dnadiff. The mean percentage of AlignedBases was used as the pairwise similarity metric, representing the average fraction of bases that can be reciprocally and uniquely aligned between the two centromeres.

### Benchmarking by leave-out cross-validation

To evaluate the performance of our method, we conducted benchmarking using leave-out cross-validation. Only samples with two complete centromere haplotypes for chr8 were included in this analysis. In each round, the dataset was randomly split into training and test sets, containing 80% and 20% of the samples, respectively. Architecture-specific, length-predictive, and normalization *k*-mers were selected exclusively from the training set, and centromere length prediction models were trained accordingly, as described above. Predictions of centromere architecture and length were then performed on the test set using short-read data, and results were compared to the ground truth obtained from the corresponding complete centromere assemblies. This procedure was repeated 10 times.

### Detection and normalization of hypomethylated CDR region inside centromere using long-read data

Using ONT R9.4.1 data from 36 HGSVC3 samples available through IGSR, basecalled with Guppy v6.5.7, we identified hypomethylated CDRs. For each sample, the Verkko assemblies of both haplotypes were concatenated, and ONT reads containing CpG methylation information were mapped to the combined assembly using minimap2 with -ax map-ont -Y -y --secondary=no. Reads shorter than 30 kb were prefiltered using chopper with --minlength 30000 [68]. After mapping, alignments were removed if less than 80% of the read length was aligned or if the normalized edit distance, calculated as NM / aligned length, exceeded 0.1. CpG methylation was quantified using modbam2bed v0.10.0 with --cpg.

Methylation dips were detected using a custom script with an approach similar to CDR-Finder (https://github.com/EichlerLab/CDR-Finder_smk). The α-satellite region, as determined by humAS annotations, was divided into 5-kb bins, and the mean methylation frequency was calculated for each bin. Bins with methylation frequency more than two standard deviations below the mean were extracted, and adjacent bins were merged. For each detected CDR, we recorded its position, number, size, and mean methylation fraction.

To compare CDR positions across samples and centromere haplotypes, genomic CDR coordinates were converted into relative positions within the HOR array. Centromeres, HOR annotations, and CDR calls were first oriented correctly. When multiple HOR arrays were present within a centromere, arrays were ordered along the centromere sequence into one consecutive HOR array, and the relative CDR position was calculated within this HOR array. For centromeres containing multiple CDRs, the mean relative CDR position across arrays was used.

This HOR-based coordinate transformation was used because our short-read method predicts HOR content rather than total centromere length. These quantities are not always equivalent, particularly on chromosomes 3 and 4, where the HOR array is interrupted by other repeat sequences.

Median and quantile values per chromosome for relative CDR position obtained in such a way and used for analysis are given in Table S3.

### Shannon diversity of centromere architecture

For each combination of chromosome and (continental) population, we quantified the diversity of centromere architecture using a normalized Shannon entropy. For each sample, the inferred centromere architectures on both haplotypes were retrieved. Only samples with a complete assignment (both haplotypes non-null) were included. For each distinct architecture, the allele count was obtained by summing the occurrences of that architecture across both haplotypes of all samples in the (continental) population and, based on the allele count, the allele frequency was computed. Based on this information the normalized Shannon entropy was calculated. Chromosomes with fewer than 3 distinct architectures or (continental) populations with fewer than 5 samples were excluded from the analysis.

### Multiple factor analysis of centromere allele frequencies

Multiple factor analysis (MFA) of centromere architectures was performed using PRINCE v0.14.0. For each population, allele frequencies of all predicted centromere architectures were calculated across chromosomes. Architecture frequencies were grouped by chromosome so that MFA balanced the contribution of each chromosome regardless of the number of architectures identified for that chromosome. MFA was then performed with PRINCE using n_components = 20 and n_iter = 5. The first four MFA dimensions were used for visualization.

### Prediction of chromosomal instability using a logistic regression model

Chromosomal instability (CIN) was modelled as a binary outcome using logistic regression. CIN status was obtained separately for each chromosome in each cancer sample from the PCAWG cohort. A chromosome was classified as unstable if it showed evidence of chromothripsis, as defined by the PCAWG project, or if it harboured copy-number alterations affecting more than 25% of its bases. The predictor variables included *TP53* mutation status, the number of single-nucleotide variants on the chromosome, tumor histology, genetic ancestry, patient age, patient sex, centromere architecture, and centromere length.

Continuous variables were standardized within each training fold using the training-set mean and standard deviation, and the same transformation was then applied to the corresponding test fold. Binary variables were retained as encoded, while categorical variables were one-hot encoded with one reference category omitted. For centromere-allele features, one allele was specified as the reference and excluded from the model, with the remaining alleles included as dosage variables.

Model performance and coefficient stability were evaluated using repeated stratified k-fold cross-validation (scikit-learn v.1.8.0). Specifically, the data were split into stratified five-fold partitions, repeated five times with a fixed random seed, yielding 25 train-test evaluations while preserving the proportion of chromosomal-instability-positive samples in each fold. For each fold, a generalized linear model with binomial error distribution and logit link function was fitted using statsmodels v.0.14.6. Predictive performance was quantified by the area under the receiver operating characteristic curve (AUROC) in both training and held-out test samples.

For each predictor, regression coefficients, standard errors, Wald test P values, likelihood-ratio test P values, odds ratios, and 95% confidence intervals were estimated within each cross-validation fold. Feature-level estimates were summarized across folds by averaging coefficients. Uncertainty was reported as 95% confidence intervals based on the t distribution. Multiple-testing correction was performed using the Benjamini–Hochberg false discovery rate procedure.

### Cancer sequencing data

Sequencing data and associated derived data, including tumor purity estimates, CNA and SNV calls, driver mutation annotations, and metadata, were obtained from the respective data sources for PCAWG [44]. Germline and tumor sequencing data were converted into de Bruijn graphs as described above.

For inference of centromere length from cancer sequencing data, a normalization value was precomputed for each sample. This value was calculated by aggregating the *k*-mer dosages of normalization *k*-mers across all chromosomal arms with a neutral copy number of 2 and then taking the mean. This approach was used to set the predicted centromere length into direct relationship to the centromere length of the matched diploid germline sample.

### Quality filtering of cancer short-read sequencing data for centromere analysis

Cancer short-read sequencing data were filtered to retain samples suitable for *k*-mer-based centromere analysis.

As a first filtering step, all samples annotated as having undergone whole-genome duplication in the original studies were excluded.

Among the remaining non-whole-genome duplicated samples, we observed that some cancer projects showed systematic shifts in predicted HOR array length between tumor and matched normal samples. These shifts were evident for all chromosomes at similar ratios, including those without CNAs affecting the centromeric region, suggesting a technical rather than biological origin. For example, liver cancer samples from one PCAWG project showed systematically longer predicted centromeres, a pattern not observed in liver cancer samples from other centers. Conversely, prostate cancer samples from one center showed systematically shorter predicted centromeres, which was not observed in data from other prostate cancer projects. In addition, some samples showed fluctuations in predicted germline and tumor centromere length on chromosomes without centromere-affecting CNAs. These samples also showed increased fragmentation of their de Bruijn graphs, consistent with reduced sequencing quality or sequencing dropout.

Because such project-specific shifts and sample-quality-associated noise compromise the comparability of *k*-mer dosage between tumor and matched normal samples, affected samples were excluded from downstream analyses.

Filtering was based on the expectation that tumor and matched germline samples should have similar inferred HOR array lengths on chromosomes without CNAs affecting the centromeric region. All chromosomes classified as copy-number neutral and not showing centromere-affecting CNAs (see **CNA and coverage based classification into whole-chromosome or arm-level aneuploidies**) were evaluated. A chromosome was considered matched if the log-ratio of inferred tumor to germline centromere length was between −0.12 and 0.12.

Samples were retained if they contained at least four centromere-copy-neutral chromosomes and if at least 50% of these chromosomes showed matching inferred centromere lengths between tumor and germline predictions. These cutoffs were chosen empirically as a trade-off between retaining sample numbers and excluding low-quality samples, as assessed by de Bruijn graph structure and copy-number profiles.

### Estimation of the number of centromeres per tumor cell

The number of centromeres per tumor cell was calculated based on the HOR length of the tumor sample *T*, the HOR length of the normal sample *N* and the tumor cell fraction *ρ* as follows:

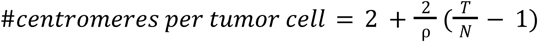

Using this approach leads to a purity adjusted estimate of the centromere content in a single clonal tumor cell.

### Gaussian mixture modeling of truncated centromeres

We estimated the number of centromeres per tumor cell across the cancer cohort and observed non-integer centromere counts, particularly in chromosomes with arm-level losses where the breakpoint was inferred to lie within the centromere. To consistently quantify distinct centromere truncation states, we fitted Gaussian mixture models to the distribution of centromere counts per tumor cell.

For this analysis, we included only events classified as arm-level losses and stratified them by chromosome and lost arm. Only groups with at least 10 events were analyzed.

For each group, we parameterized a mixture of 11 Gaussian distributions, each with a fixed standard deviation of 0.15 and means evenly spaced between 0.6 and 2.6 centromeres per tumor cell. The coefficients of the Gaussian components were fitted using SciPy v1.12.0, with bounds of [0, 1] and an equality constraint requiring the sum of all coefficients to equal 1. To reduce the risk of convergence to local optima, we performed 40 independent optimization runs with random coefficient initializations. For each group, the solution with the lowest error across all runs was retained.

To facilitate interpretation, the coefficients of the 11 Gaussian components were grouped into four categories representing distinct centromere states. Each component corresponds to a defined range of inferred centromere counts per tumor cell and, therefore, to a different degree of centromere truncation or elongation:

**Strong truncation:** components centered at 0.6, 0.8, 1.0, 1.2, and 1.4

**Partial truncation:** components centered at 1.6 and 1.8

**No detectable truncation:** components centered at 2.0 and 2.2

**Apparent centromere elongation:** components centered at 2.4 and 2.6

### CNA and coverage based classification into whole-chromosome or arm-level aneuploidies

All chromosomes from each cancer sample were classified into distinct copy number profiles consistent with whole-chromosome or arm-level alterations for downstream stratification. To this end, each chromosomal arm was rasterized into 100 equally spaced positions, and the copy number of each position was extracted from the CNA calling data for the tumor sample obtained from the respective sequencing project. Positions with more than 100 missing values across the cohort were excluded from the analysis.

For each chromosomal arm, we calculated: (a) the fraction of positions per arm falling within a given copy-number bin (CN ± 0.5), (b) the mean arm-level copy number, and (c) the mean copy number of the five positions closest to the centromere. Arms were then assigned to a copy number CN if all the following criteria were met simultaneously: (a) >50% of positions fell within CN ± 0.5, (b) the mean arm-level copy number was within CN ± 0.5, and (c) the mean copy number of centromere-adjacent positions was within CN ± 0.15. Based on the copy-number assignment of each arm, chromosomes were classified into whole-chromosome aneuploidies, arm-level losses or gains, or isochromosome-like events (**Table S5**).

In addition to CNA-based classification, we used copy-number information from normalization *k*-mers to validate copy-number states adjacent to the centromere. Because these *k*-mers were selected close to the centromere, their dosage provides an independent estimate of centromere-flanking copy number. For each chromosome, we defined the measured flanking coverage ratio as: R_measured_=P/Q, where P and Q denote the median dosage of p-arm and q-arm normalization *k*-mers, respectively.

Using the tumor purity ρ, we also define the expected flanking coverage ratio R_expected_ for each alteration class. For whole-chromosome aneuploidies, R_expected_ = 1. For arm-level losses R_expected_ = (2-ρ)/2 for p-arm losses and R_expected_ = 2/(2-ρ) for q-arm losses. For arm-level gains R_expected_ = (2+ρ)/2 for p-arm gains and R_expected_ = 2/(2+ρ) for q-arm gains. For isochromosome-like events, in which one chromosome arm has copy number 1 and the other has copy number 3, R_expected_ = (2-ρ)/(2+ρ) for iso(q)-like and R_expected_ = (2+ρ)/(2-ρ) for iso(p)-like events.

A chromosome was considered to satisfy the centromere-flanking coverage criterion for a given alteration class if R_measured_ falls between 0.6×R_expected_ and 1.4×R_expected_. Chromosomes were finally assigned to a whole-chromosome or arm-level aneuploidy class only when both the CNA-profile criteria and the centromere-flanking coverage criterion were satisfied.

During this process, we observed that the small number of normalization *k*-mers identified for the p arm of chromosome 2 (75 distinct *k*-mers after collapsing forward and reverse-complement representations) showed markedly elevated copy number in short-read data. This indicates that the normalization *k*-mers for chr2 p-arm are not reliably single-copy and therefore do not provide accurate estimates of p-arm coverage. Because this compromised coverage normalization, chromosome 2 length estimates were subsequently excluded from downstream analyses in the cancer cohort.

### Rescaling of centromere number per tumor cell

Based on the inferred HOR array length in tumor and matched normal samples, together with tumor purity estimates provided by the respective sequencing projects, we calculated the number of centromeres per tumor cell (see above). For whole-chromosome aneuploidies, these estimates showed a distinct ladder-like structure with approximately equal spacing between copy-number states. However, the distributions were not centered exactly on their expected copy numbers and were slightly shifted toward copy number 2, likely because the coverage of copy-number 2 regions was used for normalization prior to length estimation in all cancer samples.

To correct for this systematic shift, we fitted a linear regression between the inferred number of centromeres per tumor cell for whole-chromosome aneuploidies and the corresponding expected copy number. This model showed an excellent fit (R² = 0.99973) and was used to rescale centromere number estimates across all whole-chromosome and arm-level aneuploidy classes.

## Abbreviations

AFR: African
AMR: American (admixed American)
AUROC: Area under the receiver operating characteristic curve
CDR: Centromere dip region
CENP-A: Centromere protein A
CIN: Chromosomal instability
CNA: Copy-number alteration
CSA: Central/South Asian
DBG: De Bruijn graph
EAS: East Asian
EUR: European
HGDP: Human Genome Diversity Project
HGSVC: Human Genome Structural Variation Consortium
HOR: Higher-order repeat
HOROSCOPE: Higher-Order Repeat organization and Size of Centromeres using Oligonucleotide Profiles for Estimation
HPRC: Human Pangenome Reference Consortium
MEE: Middle Eastern
MFA: Multiple factor analysis
NAM: Native American
OCE: Oceanian
PCAWG: Pan-Cancer Analysis of Whole Genomes
SAS: South Asian
SNP: Single-nucleotide polymorphism
SNV: Single-nucleotide variant
T2T: Telomere-to-telomere

## Ethics approval and consent to participate

Not applicable.

## Consent for publication

Not applicable.

## Availability of data and materials

Complete human T2T assemblies used for centromere extraction were obtained from multiple publicly available sources [14,33,34,45]. All assembly download links are listed in **Table S2**.

For detection of centromere dip regions, we used basecalled ONT data from 65 samples for which CpG methylation calls were available at the time of analysis. The samples used are listed in Table S6. Data were downloaded from IGSR and the HGSVC3 ONT rebasecalled data repository (https://ftp.1000genomes.ebi.ac.uk/vol1/ftp/data_collections/HGSVC3/working/20240816_JAX_ONT_guppy6_Rebasecalled/). Only basecalled reads passing the filter with CpG methylation calls were used.

For population and cancer cohort analyses, we re-analyzed publicly available sequencing data from the 1000 Genomes Project [37,38], the Human Genome Diversity Project [39], and the Pan-Cancer Analysis of Whole Genomes Consortium [44].

HOROSCOPE, the workflow for centromere prediction from short-read data developed here, is available at https://github.com/carstenhain/HOROSCOPE/. The repository also contains the complete *k*-mer set and linear regression model required for centromere architecture and size inference. Version information for all software and tools used in this study is provided in the relevant Methods sections.

## Acknowledgments

The authors gratefully acknowledge the consortia and research groups that generated and made available human telomere-to-telomere genome assemblies used in this study. We thank the Human Pangenome Reference Consortium for providing access to assemblies prior to publication of the HPRC year 2 manuscript. In recognition of these contributions, we included the Human Pangenome Reference Consortium and the Human Genome Structural Variation Consortium therefore as banner authors. The authors are also deeply grateful to all individuals who donated biological material and contributed data to the datasets analyzed in this study. We would also like to acknowledge the National Genome Research Institute (NHGRI) for funding the following grants supporting the creation of the human pangenome reference: U41HG010972, U01HG010971, U01HG013760, U01HG013755, U01HG013748, U01HG013744, R01HG011274, and the Human Pangenome Reference Consortium (BioProject ID: PRJNA730823).

## Authors’ contributions

C.H., T.R., and J.K. conceived the project and designed the study. C.H. led the study. C.H. and T.R. performed the computational analyses. T.R. and J.K. supervised the project. C.H. wrote the original draft of the manuscript. T.R. and J.K. revised the manuscript. All authors read and approved the final manuscript.

## Human Pangenome Reference Consortium Version 2 Authors

Derek Albracht^1^, Ivan A. Alexandrov^2^, Jamie Allen^3^, Alawi A. Alsheikh-Ali^4^, Nicolas Altemose^5^, Casey Andrews^6^, Dmitry Antipov^7^, Lucinda Antonacci-Fulton^1^, Mobin Asri^8^, Marcelo Ayllon^9^, Jennifer R. Balacco^10^, Floris P. Barthel^11^, Edward A. Belter Jr^1^, Halle D. Bender^8^, Andrew P. Blair^8^, Davide Bolognini^12^, Katherine E. Bonini^13^, Christina Boucher^14^, Guillaume Bourque^15,16,17^, Silvia Buonaiuto^18^, Shuo Cao^18^, Andrew Carroll^19^, Ann M. McCartney^8^, Monika Cechova^8^, Mark J.P. Chaisson^20^, Pi-Chuan Chang^19^, Xian Chang^8^, Jitender Cheema^3^, Haoyu Cheng^21^, Claudio Ciofi^22^, Hiram Clawson^8^, Sarah Cody^1^, Vincenza Colonna^18^, Holland C. Conwell^23^, Robert Cook-Deegan^24^, Mark Diekhans^8^, Maria Angela Diroma^22^, Daniel Doerr^25,26,27^, Zheng Dong^6^, Danilo Dubocanin^5^, Richard Durbin^28,29^, Jana Ebler^25,30^, Evan E. Eichler^9,31^, Jordan M. Eizenga^8^, Parsa Eskandar^8^, Eddie Ferro^14^, Anna-Sophie Fiston-Lavier^32,33^, Sarah M. Ford^23^, Willard W. Ford^34^, Giulio Formenti^10^, Adam Frankish^3^, Mallory A. Freeberg^3^, Qichen Fu^6^, Stephanie M. Fullerton^35^, Robert S. Fulton^1^, Yan Gao^36^, Gage H. Garcia^9^, Obed A. Garcia^37^, Joshua M.V. Gardner^8^, Shilpa Garg^38^, Erik Garrison^18^, Nanibaa’ A. Garrison^39,40,41^, John E. Garza^1^, Margarita Geleta^42^, Mohammadmersad Ghorbani^43^, Tina A. Graves-Lindsay^1^, Richard E. Green^23^, Cristian Groza^44^, Bida Gu^20^, Andrea Guarracino^11,18^, Melissa Gymrek^45^, Maximilian Haeussler^8^, Leanne Haggerty^3^, Ira M. Hall^46,47^, Nancy F. Hansen^7^, Yue Hao^11^, Mohammad Amiruddin Hashmi^4^, David Haussler^8^, Prajna Hebbar^8^, Peter Heringer^25,26,27^, Glenn Hickey^8^, Todd L. Hillaker^8^, S. Nakib Hossain^3^, Neng Huang^36,48^, Sarah E. Hunt^3^, Toby Hunt^3^, Alexander G. Ioannidis^5,8^, Nafiseh Jafarzadeh^8^, Nivesh Jain^10^, Erich D. Jarvis^10,31^, Maryam Jehangir^11^, Juan Jiang^6^, Eimear E. Kenny^13^, Juhyun Kim^7^, Bonhwang Koo^10^, Sergey Koren^7^, Milinn Kremitzki^1,6^, Charles H. Langley^49^, Ben Langmead^50^, Heather A. Lawson^6^, Daofeng Li^6^, Heng Li^36,48^, Wen-Wei Liao^46,47^, Jiadong Lin^9^, Tianjie Liu^6^, Glennis A. Logsdon^51^, Ryan Lorig-Roach^8^, Jonathan LoTempio Jr^52^, Hailey Loucks^8^, Jane E. Loveland^3^, Jianguo Lu^53^, Shuangjia Lu^46,47^, Julian K. Lucas^8^, Walfred Ma^20^, Juan F. Macias-Velasco^1,6,54^, Kateryna D. Makova^55^, Maximillian G. Marin^36,48^, Christopher Markovic^1^, Tobias Marschall^25,30^, Franco L. Marsico^18^, Fergal J. Martin^3^, Mira Mastoras^8^, Capucine Mayoud^32^, Brandy McNulty^8^, Jack A. Medico^10^, Julian M. Menendez^8^, Karen H. Miga^8^, Anna Minkina^56^, Matthew W. Mitchell^57^, Saswat K. Mohanty^58^, Younes Mokrab^43,59,60^, Jean Monlong^61^, Shabir Moosa^43^, Avelina Moreno-Ochando^62,63^, Shinichi Morishita^64^, Jonathan M. Mudge^3^, Katherine M. Munson^9^, Njagi Mwaniki^65^, Nasna Nassir^4^, Chiara Natali^22^, Shloka Negi^8^, Lingbin Ni^9^, Adam M. Novak^8^, Faith Okamoto^8^, Pilar N. Ossorio^66^, Chie Owa^64^, Sadye Paez^10^, Benedict Paten^8^, Clelia Peano^12,67^, Adam M. Phillippy^7^, Brandon D. Pickett^7^, Laura Pignata^18^, Nadia Pisanti^65^, David Porubsky^9,68^, Pjotr Prins^18^, Anandi Radhakrishnan^8^, T. Rhyker Ranallo-Benavidez^11^, Brian J. Raney^8^, Mikko Rautiainen^69^, Alessandro Raveane^12^, Luyao Ren^9,31^, Arang Rhie^7^, Fedor Ryabov^70,71^, Samuel Sacco^23^, Farnaz Salehi^18^, Michael C. Schatz^50,72^, Laura B. Scheinfeldt^73^, Aarushi Sehgal^34^, William E. Seligmann^23^, Mahsa Shabani^74^, Kishwar Shafin^19^, Shadi Shahatit^32^, Ruhollah Shemirani^13^, Vikram S. Shivakumar^50^, Swati Sinha^3^, Jouni Sirén^8^, Linnéa Smeds^58^, Steven J. Solar^7^, Marco Sollitto^10,22^, Nicole Soranzo^12,28,75^, Andrew B. Stergachis^9,56^, Marie-Marthe Suner^3^, Yoshihiko Suzuki^64^, Arda Söylev^25,30^, Ahmad Abou Tayoun^76,77^, Jack A.S. Tierney^3^, Chad Tomlinson^1^, Francesca Floriana Tricomi^3^, Mohammed Uddin^4,78^, Matteo Tommaso Ungaro^23,79^, Rahul Varki^14^, Flavia Villani^18^, Ivo Violich^8^, Mitchell R. Vollger,^56^, Brian P. Walenz^7^, Charles Wang^80^, Lisa E. Wang^13^, Ting Wang^1,6,54^, Aaron M. Wenger^81^, Conor V. Whelan^10^, Zilan Xin^6^, Zheng Xu^6^, Kai Ye^82^, DongAhn Yoo^9^, Wenjin Zhang^6^, Ying Zhou^36^, Xiaoyu Zhuo^6^, Giulia Zunino^12^

### Affiliations

1. McDonnell Genome Institute, Washington University School of Medicine, St. Louis, MO 63108, USA
2. Department of Human Molecular Genetics and Biochemistry, Faculty of Medical and Health Sciences, Tel Aviv University, Tel Aviv 69978, Israel
3. European Molecular Biology Laboratory, European Bioinformatics Institute (EMBL-EBI), Wellcome Genome Campus, Hinxton, Cambridge CB10 1SD, UK
4. Center for Applied and Translational Genomics (CATG), Mohammed Bin Rashid University of Medicine and Health Sciences, Dubai Health, Dubai, UAE
5. Department of Genetics, Stanford University, Palo Alto, CA 94304 USA
6. Department of Genetics, Washington University School of Medicine, St. Louis, MO 63110, USA
7. Genome Informatics Section, Center for Genomics and Data Science Research, National Human Genome Research Institute, National Institutes of Health, Bethesda, MD 20892, USA
8. UC Santa Cruz Genomics Institute, University of California, Santa Cruz, CA 95060, USA
9. Department of Genome Sciences, University of Washington School of Medicine, Seattle, WA 98195, USA
10. The Vertebrate Genome Laboratory, The Rockefeller University, New York, NY 10065, USA
11. Bioinnovation and Genome Sciences, The Translational Genomics Research Institute (TGen), Phoenix, AZ 85004, USA
12. Human Technopole, Milan, Italy
13. Institute for Genomic Health, Icahn School of Medicine at Mount Sinai, New York, NY 10029, USA
14. Department of Computer and Information Science and Engineering, University of Florida, Gainesville, FL 32611, USA
15. Canadian Center for Computational Genomics, McGill University, Montréal, QC H3A 0G1, Canada
16. Department of Human Genetics, McGill University, Montréal, QC H3A 0G1, Canada
17. Victor Phillip Dahdaleh Institute of Genomic Medicine, Montréal, QC H3A 0G1, Canada
18. Department of Genetics, Genomics and Informatics, University of Tennessee Health Science Center, Memphis, TN 38163, USA
19. Google LLC, Mountain View, CA 94043, USA
20. Quantitative and Computational Biology, University of Southern California, Los Angeles, CA 90089, USA
21. Department of Biomedical Informatics and Data Science, Yale School of Medicine, New Haven, CT 06510, USA
22. Department of Biology, University of Florence, Sesto Fiorentino, FI 50019, Italy
23. Department of Ecology and Evolutionary Biology, University of California, Santa Cruz, CA 95060, USA
24. Arizona State University, Consortium for Science, Policy & Outcomes, Washington, DC 20006, USA
25. Center for Digital Medicine, Heinrich Heine University Düsseldorf, Düsseldorf, NRW, DE
26. Department for Endocrinology and Diabetology at the Medical Faculty and University Hospital Düsseldorf, Heinrich Heine University Düsseldorf, Düsseldorf, NRW, DE
27. Paul-Langerhans-Group Computational Diabetology, German Diabetes Center (DDZ) and Leibniz Institute for Diabetes Research, Düsseldorf, NRW, DE
28. Wellcome Sanger Institute, Genome Campus, Hinxton, CB10 1RQ, UK
29. Department of Genetics, University of Cambridge, Cambridge, CB2 3EH, UK
30. Institute for Medical Biometry and Bioinformatics, Medical Faculty and University Hospital Düsseldorf, Heinrich Heine University, Düsseldorf, NRW, DE
31. Howard Hughes Medical Institute, Chevy Chase, MD 20815, USA
32. ISEM, Univ Montpellier, CNRS, IRD, Montpellier, FR
33. Institut Universitaire de France, Paris, FR
34. Department of Computer Science and Engineering, University of California San Diego, La Jolla, CA 92093, USA
35. Department of Bioethics & Humanities, University of Washington School of Medicine, Seattle, WA 98195, USA
36. Department of Data Science, Dana-Farber Cancer Institute, Boston, MA 02215, USA
37. Department of Anthropology, University of Kansas, Lawrence, KS 66045, USA
38. School of Health Sciences, University of Manchester, Manchester M13 9PL, UK
39. Traditional, ancestral and unceded territory of the Gabrielino/Tongva peoples, Institute for Society & Genetics, University of California, Los Angeles, Los Angeles, CA 90095, USA
40. Traditional, ancestral and unceded territory of the Gabrielino/Tongva peoples, Institute for Precision Health, David Geffen School of Medicine, University of California, Los Angeles, Los Angeles, CA 90095, USA
41. Traditional, ancestral and unceded territory of the Gabrielino/Tongva peoples, Division of General Internal Medicine & Health Services Research, David Geffen School of Medicine, University of California, Los Angeles, Los Angeles, CA 90095, USA
42. Department of Electrical Engineering and Computer Science, University of California, Berkeley, Berkeley, CA 94720, USA
43. Medical and Population Genomics Lab, Sidra Medicine, Doha, Qatar
44. Montreal Heart Institute, Montréal, QC, Canada
45. Department of Pediatrics, University of California San Diego, La Jolla, CA 92093, USA
46. Center for Genomic Health, Yale University School of Medicine, New Haven, CT 06510, USA
47. Department of Genetics, Yale University School of Medicine, New Haven, CT 06510, USA
48. Department of Biomedical Informatics, Harvard Medical School, Boston, MA 02115, USA
49. Department of Evolution and Ecology and the Center for Population Biology, University of California, One Shields, Davis, CA 95616, USA
50. Department of Computer Science, Johns Hopkins University, Baltimore, MD 21218, USA
51. Department of Genetics, Epigenetics Institute, Perelman School of Medicine, University of Pennsylvania, Philadelphia, PA 19104, USA
52. Department of Pediatrics, Division of Genetics, School of Medicine, University of California, Irvine, CA 92697, USA
53. Sun Yat-sen University, Guangzhou, China
54. Edison Family Center for Genome Sciences & Systems Biology, Washington University School of Medicine, St. Louis, MO 63110, USA
55. Department of Biology and Center for Medical Genomics, Penn State University, University Park, PA 16802, USA
56. Division of Medical Genetics, Department of Medicine, University of Washington School of Medicine, Seattle, WA 98195, USA
57. The Jackson Laboratory for Genomic Medicine, Farmington, CT 06032, USA
58. Department of Biology, Penn State University, University Park, PA 16802, USA
59. Department of Biomedical Science, College of Health Sciences, Qatar University, Doha, Qatar
60. Department of Genetic Medicine, Weill Cornell Medicine-Qatar, Doha, Qatar
61. IRSD - Digestive Health Research Institute, University of Toulouse, INSERM, INRAE, ENVT, UPS, Toulouse, FR
62. MATCH biosystems, S.L., Elche, Spain
63. Universidad Miguel Hernández de Elche, Elche, Spain
64. Department of Computational Biology and Medical Sciences, The University of Tokyo, Kashiwa, Chiba 277-8561, Japan
65. Department of Computer Science, University of Pisa, Pisa, Italy
66. Law School, University of Wisconsin-Madison, Madison, WI 53706, USA
67. Institute of Genetics and Biomedical Research, UoS of Milan, National Research Council, Milan, Italy
68. Genome Biology Unit, European Molecular Biology Laboratory (EMBL), Heidelberg, DE
69. Institute for Molecular Medicine Finland, Helsinki Institute of Life Science, University of Helsinki, Helsinki, Finland
70. The Center for Bio- and Medical Technologies, Moscow, RUS
71. Centre for Biomedical Research and Technology, HSE University, Moscow, RUS
72. Department of Biology, Johns Hopkins University, Baltimore, MD 21218, USA
73. Coriell Institute for Medical Research, Camden, NJ 08103, USA
74. University of Amsterdam, Amsterdam, Netherlands
75. School of Clinical Medicine, University of Cambridge, Cambridge, CB2 0SP, UK
76. Center for Genomic Discovery, Mohammed Bin Rashid University, Dubai Health, UAE
77. Dubai Health Genomic Medicine Center, Dubai Health, UAE
78. GenomeArc Inc, Mississauga, ON, Canada
79. Department of Biology and Biotechnologies “Charles Darwin”, University of Rome “La Sapienza”, Rome 00185, IT
80. Center for Genomics, Loma Linda University School of Medicine, Loma Linda, CA 92350, USA
81. PacBio, Menlo Park, CA 94025, USA
82. The first affiliated hospital of Xi’an Jiaotong University, Xi’an Jiaotong University, Xi’an, Shaanxi, 710049, China

## Human Genome Structural Variation Consortium Authors

Hufsah Ashraf^1,2^, Peter A. Audano^3^, Marcelo Ayllon^4^, Andrey Azov^5^, Parithi Balachandran^3^, Anna O. Basile^6^, Christine R. Beck^3,7^, Marc Jan Bonder^8-10^, Lucy Brooks^5^, Marta

Byrska-Bishop^6^, Mark J. P. Chaisson^11^, Zechen Chong^12^, André Corvelo^6^ , Jonathan Crabtree^13^, Scott E. Devine^13^, Peter Ebert^14,2^, Jana Ebler^1,2^, Evan E. Eichler^4,15^, Aine Fairbrother-Browne^5^, Chia-Hsuan Fan^12^, Mallory Freeberg^5^, Shenghan Gao^16^, Mark B. Gerstein^17,18^, Bida Gu^11^, Pille Hallast^3^, Patrick Hasenfeld^19^, Mir Henglin^1,2^, Kendra Hoekzema^4^, Kaili Hu^12^, Sarah Hunt^5^, Matthew Jensen^17,18^, Yunzhe Jiang^17,18^, Kwondo Kim^3^, Miriam K. Konkel^20,21^, Jan O. Korbel^19^, Youngjun Kwon^4^, Peter M. Lansdorp^22^, Charles Lee^3^, Tiffany Leung^22^, Jiaqi Li^17,18^, Chong Li^23,24^, Jiadong Lin^4^, Mark Loftus^3^, Glennis A. Logsdon^16^, Tobias Marschall^1,2^, Gianni V. Martino^20^, Ryan E. Mills^25^, Yulia Mostovoy^26^, Katherine M. Munson^4^, Giuseppe Narzisi^6^, Lingbin Ni^4^, Keisuke K. Oshima^16^, Carolyn Paisie^3^, Samarendra Pani^1,2^, Zishan Peng^12^, David Porubsky^4^, Timofey Prodanov^1,2^, Keon Rabbani^11^, Tobias Rausch^19^, Xinghua Shi^23,24^, Yuwei Song^12^, Kaitlyn Sun^4^, Likhitha Surapaneni^5^, Michael E. Talkowski^27-29^, Vasiliki Tsapalou^19^, Andres Veidenberg^5^, Feyza Yilmaz^3^, DongAhn Yoo^4^, Xuefang Zhao^27-29^, Weichen Zhou^25^, Qihui Zhu^3^ , Michael C. Zody^6^

### Affiliations

1. Institute for Medical Biometry and Bioinformatics, Medical Faculty and University Hospital Düsseldorf, Heinrich Heine University, Düsseldorf, Germany
2. Center for Digital Medicine, Heinrich Heine University, Düsseldorf, Germany
3. The Jackson Laboratory for Genomic Medicine, Farmington, CT, USA
4. University of Washington School of Medicine, Department of Genome Sciences, Seattle, WA, USA
5. European Molecular Biology Laboratory, Wellcome Genome Campus, European Bioinformatics Institute, Cambridge, UK
6. New York Genome Center, New York, NY, USA
7. The University of Connecticut Health Center, Farmington, CT, USA
8. Department of Genetics, University Medical Center Groningen, University of Groningen, Groningen, The Netherlands
9. Oncode Institute, Utrecht, The Netherlands
10. Division of Computational Genomics and Systems Genetics, German Cancer Research Center (DKFZ), Heidelberg, Germany
11. Department of Quantitative and Computational Biology, University of Southern California, Los Angeles, CA, USA
12. Department of Biomedical Informatics and Data Science, Heersink School of Medicine, University of Alabama, Birmingham, AL, USA
13. Institute for Genome Sciences, University of Maryland School of Medicine, Baltimore, MD, USA
14. Core Unit Bioinformatics, Medical Faculty and University Hospital Düsseldorf, Heinrich Heine University, Düsseldorf, Germany
15. Howard Hughes Medical Institute, University of Washington, Seattle, WA, USA
16. Department of Genetics, Epigenetics Institute, Perelman School of Medicine, University of Pennsylvania, Philadelphia, PA, USA
17. Department of Molecular Biophysics and Biochemistry, Yale University, New Haven, CT, USA
18. Program in Computational Biology and Bioinformatics, Yale University, New Haven, CT, USA
19. European Molecular Biology Laboratory (EMBL), Genome Biology Unit, Heidelberg, Germany
20. Clemson University, Department of Genetics & Biochemistry, Clemson, SC, USA
21. Institute for Human Genetics, Clemson University, Greenwood, SC, USA
22. Terry Fox Laboratory, BC Cancer Agency, Vancouver, BC, Canada
23. Department of Computer and Information Sciences, Temple University, Philadelphia, PA, USA
24. Institute for Genomics and Evolutionary Medicine, Temple University, Philadelphia, PA, USA
25. Department of Computational Medicine & Bioinformatics, University of Michigan, MI, USA
26. Cardiovascular Research Institute and Institute for Human Genetics, UCSF School of Medicine, CA, USA
27. Program in Medical and Population Genetics, Broad Institute of MIT and Harvard, Cambridge, MA, USA
28. Center for Genomic Medicine, Massachusetts General Hospital, Boston, MA, USA
29. Department of Neurology, Massachusetts General Hospital and Harvard Medical School, MA, USA

## References

1. Miga KH, Alexandrov IA. Variation and Evolution of Human Centromeres: A Field Guide and Perspective. Annu Rev Genet. 2021;55:583–602. 10.1146/annurev-genet-071719-020519

2. Altemose N, Logsdon GA, Bzikadze AV, Sidhwani P, Langley SA, Caldas GV, et al. Complete genomic and epigenetic maps of human centromeres. Science [Internet]. American Association for the Advancement of Science (AAAS); 2022 [cited 2025 July 9];376. 10.1126/science.abl4178

3. Logsdon GA, Rozanski AN, Ryabov F, Potapova T, Shepelev VA, Catacchio CR, et al. The variation and evolution of complete human centromeres. Nature. Springer Science and Business Media LLC; 2024;629:136–45. 10.1038/s41586-024-07278-3

4. Gao S, Oshima KK, Chuang S-C, Loftus M, Montanari A, Gordon DS, et al. A global view of human centromere variation and evolution [Internet]. Genomics; 2025 [cited 2026 May 6]. 10.64898/2025.12.09.693231

5. Jaggi KE, Hoyt SJ, O’Neill RJ, Sullivan BA. A genomic and epigenomic view of human centromeres. Nat Rev Genet. 2026;27:371–84. 10.1038/s41576-025-00923-1

6. De Rop V, Padeganeh A, Maddox PS. CENP-A: the key player behind centromere identity, propagation, and kinetochore assembly. Chromosoma. 2012;121:527–38. 10.1007/s00412-012-0386-5

7. McAinsh AD, Marston AL. The Four Causes: The Functional Architecture of Centromeres and Kinetochores. Annu Rev Genet. 2022;56:279–314. 10.1146/annurev-genet-072820-034559

8. Miga KH, Koren S, Rhie A, Vollger MR, Gershman A, Bzikadze A, et al. Telomere-to-telomere assembly of a complete human X chromosome. Nature. 2020;585:79–84. 10.1038/s41586-020-2547-7

9. Logsdon GA, Vollger MR, Hsieh P, Mao Y, Liskovykh MA, Koren S, et al. The structure, function and evolution of a complete human chromosome 8. Nature. 2021;593:101–7. 10.1038/s41586-021-03420-7

10. Gershman A, Sauria MEG, Guitart X, Vollger MR, Hook PW, Hoyt SJ, et al. Epigenetic patterns in a complete human genome. Science. 2022;376:eabj5089. 10.1126/science.abj5089

11. Salinas-Luypaert C, Dubocanin D, Lee RJ, Andrade Ruiz L, Gamba R, Grison M, et al. DNA methylation influences human centromere positioning and function. Nat Genet. 2025;57:2509–21. 10.1038/s41588-025-02324-w

12. Sidhwani P, Schwartz JP, Fryer KA, Straight AF. Histone H3 lysine methyltransferase activities control compartmentalization of human centromeres [Internet]. Genomics; 2025 [cited 2026 May 6]. 10.1101/2025.07.01.662447

13. Carty BL, Dubocanin D, Murillo-Pineda M, Dumont M, Volpe E, Mikulski P, et al. Heterochromatin boundaries maintain centromere position, size and number. Nat Struct Mol Biol. 2026;33:220–34. 10.1038/s41594-025-01706-2

14. Logsdon GA, Ebert P, Audano PA, Loftus M, Porubsky D, Ebler J, et al. Complex genetic variation in nearly complete human genomes. Nature. 2025;644:430–41. 10.1038/s41586-025-09140-6

15. Porubsky D, Dashnow H, Sasani TA, Logsdon GA, Hallast P, Noyes MD, et al. Human de novo mutation rates from a four-generation pedigree reference. Nature. 2025;643:427–36. 10.1038/s41586-025-08922-2

16. Xu Y, Loucks H, Menendez J, Ryabov F, Lucas JK, Cechova M, et al. Haplotype-resolved centromeric chromatin organization from a complete diploid human genome [Internet]. Genomics; 2026 [cited 2026 May 6]. 10.64898/2026.03.27.714900

17. Mahlke MA, Lumerman L, Nath P, Chittenden C, Hoyt S, Koeppel J, et al. Evolution and instability of human centromeres are accelerated by heterochromatin boundary loss and CENP-A overexpression [Internet]. Cell Biology; 2025 [cited 2026 May 6]. 10.1101/2025.02.03.636285

18. Amor DJ, Choo KHA. Neocentromeres: Role in Human Disease, Evolution, and Centromere Study. Am J Hum Genet. 2002;71:695–714. 10.1086/342730

19. Marshall OJ, Chueh AC, Wong LH, Choo KHA. Neocentromeres: New Insights into Centromere Structure, Disease Development, and Karyotype Evolution. Am J Hum Genet. 2008;82:261–82. 10.1016/j.ajhg.2007.11.009

20. Fukagawa T, Earnshaw WC. Neocentromeres. Curr Biol. 2014;24:R946–7. 10.1016/j.cub.2014.08.032

21. Hoyt SJ, Hartley GA, Antopia MC, Tullius TW, Taylor DJ, Tillquist NM, et al. Haplotype-Resolved Genomics Reveals Conserved Chromatin Architecture and Epigenetic Constraints of Human Neocentromeres [Internet]. Genomics; 2025 [cited 2026 May 6]. 10.64898/2025.12.23.696241

22. Barra V, Fachinetti D. The dark side of centromeres: types, causes and consequences of structural abnormalities implicating centromeric DNA. Nat Commun [Internet]. Springer Science and Business Media LLC; 2018 [cited 2025 July 9];9. 10.1038/s41467-018-06545-y

23. Paul S, Kaplan MH, Khanna D, McCourt PM, Saha AK, Tsou P-S, et al. Centromere defects, chromosome instability, and cGAS-STING activation in systemic sclerosis. Nat Commun. 2022;13:7074. 10.1038/s41467-022-34775-8

24. Mastrorosa FK, Rozanski AN, Harvey WT, Knuth J, Garcia G, Munson KM, et al. Complete chromosome 21 centromere sequences from a Down syndrome family reveal size asymmetry and differences in kinetochore attachment [Internet]. Cold Spring Harbor Laboratory; 2024 [cited 2025 July 9]. 10.1101/2024.02.25.581464

25. Cosenza MR, Rodriguez-Martin B, Korbel JO. Structural Variation in Cancer: Role, Prevalence, and Mechanisms. Annu Rev Genomics Hum Genet. 2022;23:123–52. 10.1146/annurev-genom-120121-101149

26. Steele CD, Abbasi A, Islam SMA, Bowes AL, Khandekar A, Haase K, et al. Signatures of copy number alterations in human cancer. Nature. 2022;606:984–91. 10.1038/s41586-022-04738-6

27. Martinez-A C, Van Wely KHM. Centromere fission, not telomere erosion, triggers chromosomal instability in human carcinomas. Carcinogenesis. 2011;32:796–803. 10.1093/carcin/bgr069

28. Potapova T, Gorbsky G. The Consequences of Chromosome Segregation Errors in Mitosis and Meiosis. Biology. 2017;6:12. 10.3390/biology6010012

29. Choo Z-N, Behr JM, Deshpande A, Hadi K, Yao X, Tian H, et al. Most large structural variants in cancer genomes can be detected without long reads. Nat Genet. 2023;55:2139–48. 10.1038/s41588-023-01540-6

30. Shih J, Sarmashghi S, Zhakula-Kostadinova N, Zhang S, Georgis Y, Hoyt SH, et al. Cancer aneuploidies are shaped primarily by effects on tumour fitness. Nature. 2023;619:793–800. 10.1038/s41586-023-06266-3

31. Watson EV, Lee JJ-K, Gulhan DC, Melloni GEM, Venev SV, Magesh RY, et al. Chromosome evolution screens recapitulate tissue-specific tumor aneuploidy patterns. Nat Genet. 2024;56:900–12. 10.1038/s41588-024-01665-2

32. Liao W-W, Asri M, Ebler J, Doerr D, Haukness M, Hickey G, et al. A draft human pangenome reference. Nature. Springer Science and Business Media LLC; 2023;617:312–24. 10.1038/s41586-023-05896-x

33. Nurk S, Koren S, Rhie A, Rautiainen M, Bzikadze AV, Mikheenko A, et al. The complete sequence of a human genome. Science. American Association for the Advancement of Science (AAAS); 2022;376:44–53. 10.1126/science.abj6987

34. The Human Pangenome Reference Consortium. The human pangenome second release. to_appear_on_bioRxiv.

35. Shiraishi Y, Ochi Y, Sugawa M, Sakamoto Y, Kimura K, Tsujimura T, et al. Rare k-mers reveal centromere haplogroups underlying human diversity and cancer translocations [Internet]. Genomics; 2025 [cited 2026 May 22]. 10.1101/2025.07.26.666712

36. Ranallo-Benavidez TR, Chen Y-A, Potapova T, Alanko J, Loucks H, Lucas J, et al. KaryoScope: rapid, alignment-free sequence annotation for the pangenome era [Internet]. Bioinformatics; 2026 [cited 2026 May 21]. 10.64898/2026.05.15.725544

37. The 1000 Genomes Project Consortium, Corresponding authors, Auton A, Abecasis GR, Steering committee, Altshuler DM, et al. A global reference for human genetic variation. Nature. 2015;526:68–74. 10.1038/nature15393

38. Byrska-Bishop M, Evani US, Zhao X, Basile AO, Abel HJ, Regier AA, et al. High-coverage whole-genome sequencing of the expanded 1000 Genomes Project cohort including 602 trios. Cell. 2022;185:3426–3440.e19. 10.1016/j.cell.2022.08.004

39. Bergström A, McCarthy SA, Hui R, Almarri MA, Ayub Q, Danecek P, et al. Insights into human genetic variation and population history from 929 diverse genomes. Science. 2020;367:eaay5012. 10.1126/science.aay5012

40. Koenig Z, Yohannes MT, Nkambule LL, Zhao X, Goodrich JK, Kim HA, et al. A harmonized public resource of deeply sequenced diverse human genomes. Genome Res. 2024;34:796–809. 10.1101/gr.278378.123

41. Schloissnig S, Pani S, Ebler J, Hain C, Tsapalou V, Söylev A, et al. Structural variation in 1,019 diverse humans based on long-read sequencing. Nature. 2025;644:442–52. 10.1038/s41586-025-09290-7

42. Frichot E, Mathieu F, Trouillon T, Bouchard G, François O. Fast and Efficient Estimation of Individual Ancestry Coefficients. Genetics. 2014;196:973–83. 10.1534/genetics.113.160572

43. Fortes-Lima C, Bybjerg-Grauholm J, Marin-Padrón LC, Gomez-Cabezas EJ, Bækvad-Hansen M, Hansen CS, et al. Exploring Cuba’s population structure and demographic history using genome-wide data. Sci Rep. 2018;8:11422. 10.1038/s41598-018-29851-3

44. The ICGC/TCGA Pan-Cancer Analysis of Whole Genomes Consortium, Aaltonen LA, Abascal F, Abeshouse A, Aburatani H, Adams DJ, et al. Pan-cancer analysis of whole genomes. Nature. 2020;578:82–93. 10.1038/s41586-020-1969-6

45. Logsdon GA, Rozanski AN, Ryabov F, Potapova T, Shepelev VA, Catacchio CR, et al. The variation and evolution of complete human centromeres. Nature. Springer Science and Business Media LLC; 2024;629:136–45. 10.1038/s41586-024-07278-3

46. Volpe E, Colantoni A, Corda L, Di Tommaso E, Pelliccia F, Ottalevi R, et al. The reference genome of the human diploid cell line RPE-1. Nat Commun. 2025;16:7751. 10.1038/s41467-025-62428-z

47. Cheng H, Qu H, McKenzie S, Lawrence KR, Windsor R, Vella M, et al. Efficient near-telomere-to-telomere assembly of nanopore simplex reads. Nature [Internet]. 2026 [cited 2026 May 10]; 10.1038/s41586-026-10105-6

48. Gross C, Potabattula R, Cheng F, Leuchtenberg S, Hartung HS, Kristmann B, et al. Single-Platform Nanopore Sequencing Enables Diploid Telomere-to-Telomere Genome Assembly and Haplotype-Resolved 3D Chromatin Maps [Internet]. Genomics; 2026 [cited 2026 May 10]. 10.64898/2026.03.19.712851

49. Taliun D, Harris DN, Kessler MD, Carlson J, Szpiech ZA, Torres R, et al. Sequencing of 53,831 diverse genomes from the NHLBI TOPMed Program. Nature. 2021;590:290–9. 10.1038/s41586-021-03205-y

50. The UK Biobank Whole-Genome Sequencing Consortium, Manuscript Writing Group, Carss K, Halldorsson BV, Hou L, Liu J, et al. Whole-genome sequencing of 490,640 UK Biobank participants. Nature. 2025;645:692–701. 10.1038/s41586-025-09272-9

51. Dutka T, Faust EJ, Gallagher CS, Hyams T, Kozlowski E, Landis E, et al. All of Us Research Program year in review: 2024. Am J Hum Genet. 2025;112:1983–7. 10.1016/j.ajhg.2025.08.003

52. Zheng Y, Ahmad K, Henikoff S. Total whole-arm chromosome losses predict malignancy in human cancer. Proc Natl Acad Sci. 2025;122:e2505385122. 10.1073/pnas.2505385122

53. Irvine DV, Amor DJ, Perry J, Sirvent N, Pedeutour F, Choo KHA, et al. Chromosome size and origin as determinants of the level of CENP-A incorporation into human centromeres. Chromosome Res. 2004;12:805–15. 10.1007/s10577-005-5377-4

54. Iwata-Otsubo A, Dawicki-McKenna JM, Akera T, Falk SJ, Chmátal L, Yang K, et al. Expanded Satellite Repeats Amplify a Discrete CENP-A Nucleosome Assembly Site on Chromosomes that Drive in Female Meiosis. Curr Biol. 2017;27:2365–2373.e8. 10.1016/j.cub.2017.06.069

55. McNulty SM, Sullivan LL, Sullivan BA. Human Centromeres Produce Chromosome-Specific and Array-Specific Alpha Satellite Transcripts that Are Complexed with CENP-A and CENP-C. Dev Cell. 2017;42:226–240.e6. 10.1016/j.devcel.2017.07.001

56. Sohn M-H, Dubocanin D, Vollger MR, Kwon Y, Minkina A, Munson KM, et al. A telomere-to-telomere map of somatic mutation burden and functional impact in cancer [Internet]. Genomics; 2025 [cited 2026 May 7]. 10.1101/2025.10.10.681725

57. Dong S, Xing X, Cechova M, Loucks H, Vijayalingam S, Neilson A, et al. Fully T2T pedigree assemblies reveal genetic stability and epigenetic plasticity of human centromeres across inheritance and cell-fate transitions [Internet]. Genomics; 2026 [cited 2026 June 19]. 10.64898/2026.02.14.705860

58. Wagner J, Keskus AG, Oshima KK, Ranallo-Benavidez TR, McDaniel J, Sikic M, et al. A complete human pancreatic cancer genome [Internet]. Genomics; 2026 [cited 2026 June 9]. 10.64898/2026.05.01.722316

59. Showman S, Talbert PB, Xu Y, Adeyemi RO, Henikoff S. Expansion of human centromeric arrays in cells undergoing break-induced replication. Cell Rep. 2024;43:113851. 10.1016/j.celrep.2024.113851

60. Quinlan AR, Hall IM. BEDTools: a flexible suite of utilities for comparing genomic features. Bioinformatics. 2010;26:841–2. 10.1093/bioinformatics/btq033

61. Li H. Minimap2: pairwise alignment for nucleotide sequences. Birol I, editor. Bioinformatics. 2018;34:3094–100. 10.1093/bioinformatics/bty191

62. Rausch T, Fritz MH-Y, Untergasser A, Benes V. Tracy: basecalling, alignment, assembly and deconvolution of sanger chromatogram trace files. BMC Genomics. 2020;21:230. 10.1186/s12864-020-6635-8

63. Li H. Aligning sequence reads, clone sequences and assembly contigs with BWA-MEM [Internet]. arXiv; 2013 [cited 2026 May 7]. 10.48550/ARXIV.1303.3997

64. Song L, Florea L, Langmead B. Lighter: fast and memory-efficient sequencing error correction without counting. Genome Biol. 2014;15:509. 10.1186/s13059-014-0509-9

65. Chikhi R, Limasset A, Medvedev P. Compacting de Bruijn graphs from sequencing data quickly and in low memory. Bioinformatics. 2016;32:i201–8. 10.1093/bioinformatics/btw279

66. Marçais G, Kingsford C. A fast, lock-free approach for efficient parallel counting of occurrences of *k* -mers. Bioinformatics. 2011;27:764–70. 10.1093/bioinformatics/btr011

67. Marçais G, Delcher AL, Phillippy AM, Coston R, Salzberg SL, Zimin A. MUMmer4: A fast and versatile genome alignment system. Darling AE, editor. PLOS Comput Biol. 2018;14:e1005944. 10.1371/journal.pcbi.1005944

68. De Coster W, Rademakers R. NanoPack2: population-scale evaluation of long-read sequencing data. Alkan C, editor. Bioinformatics. 2023;39:btad311. 10.1093/bioinformatics/btad311

