## Supplementary figures for "HOROSCOPE: Decoding human centromere architecture from short reads using *k*-mer signatures"

A

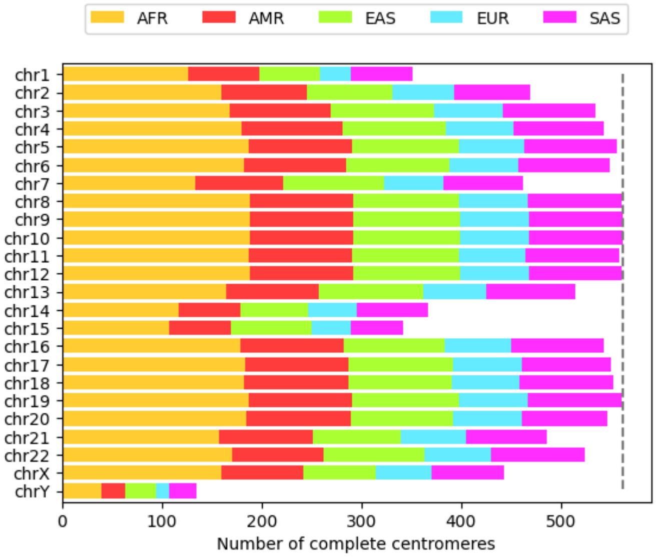


B

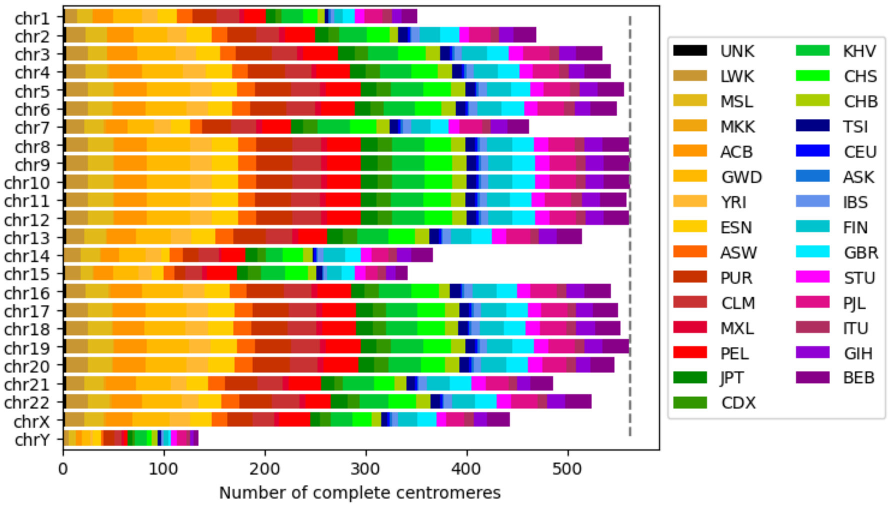


**Figure S1: Number of complete centromeres extracted per chromosome from (near) T2T assemblies, colored by superpopulation (A) or population (B).**
The dashed line at 562 marks the maximum possible number of haplotypes, based on 280 diploid and 2 haploid samples used in this analysis.

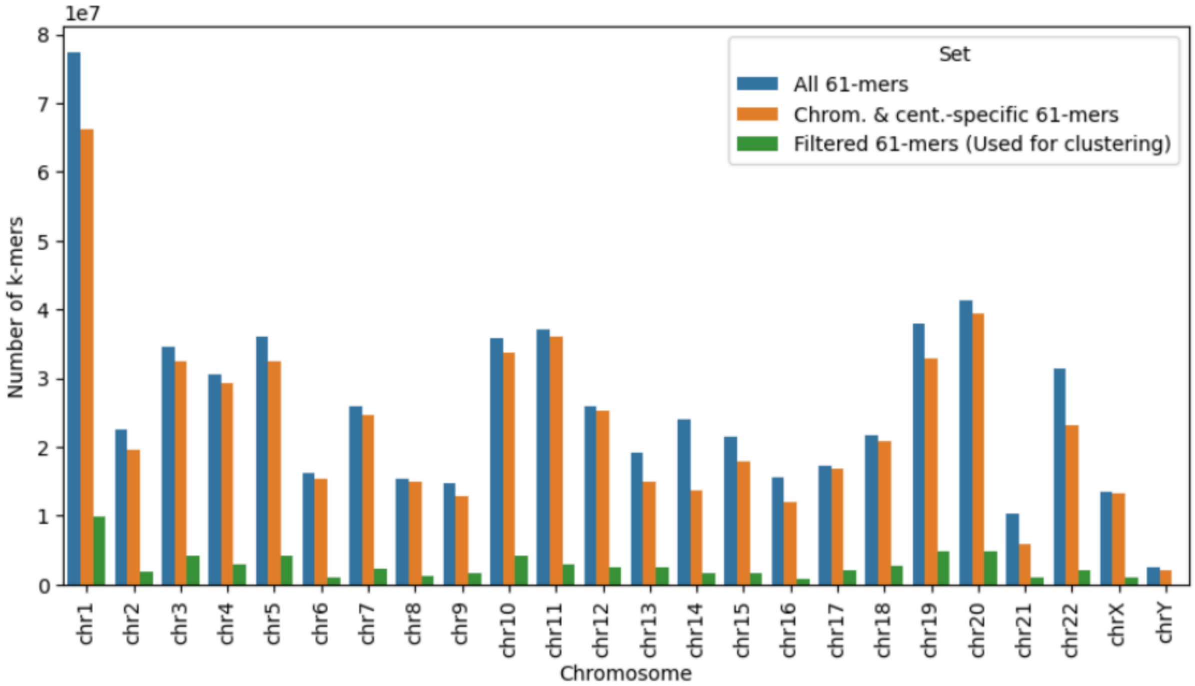


**Figure S2: Number of unique 61-mers extracted from population-scale centromere assemblies.**

Blue bars indicate the total number of 61-mers derived from all complete centromere assemblies for each chromosome. For the detection of architecture-specific *k*-mers, all 61-mers present in the centromeres of other chromosomes or in the non-centromeric regions of the CHM13 genome were removed (orange). For clustering, all rare and highly shared *k*-mers were excluded, which substantially reduced the number of 61-mers per chromosome (green).

**Figure S3: 61-mer based clustering of human centromeres.**
For each human autosome (chr1-22) k-mer based clustering was performed (**A-V**). For each chromosome the top panel shows the phylogenetic tree, the middle panel displays a matrix of pairwise similarity between haplotypes, and the lower panel indicates the superpopulation of the individual haplotype. The HOR structure of each haplotype is shown to the right, with the full contig shown in light gray and HORs colored according to their repeat unit length. A full legend with all color-coded HOR repeat counts is given at the end of this figure in panel **W**. Superpopulation colors follow the 1000 Genomes Project scheme: African (yellow), American (red), East Asian (green), European (blue), and South Asian (purple).

**A** chr1

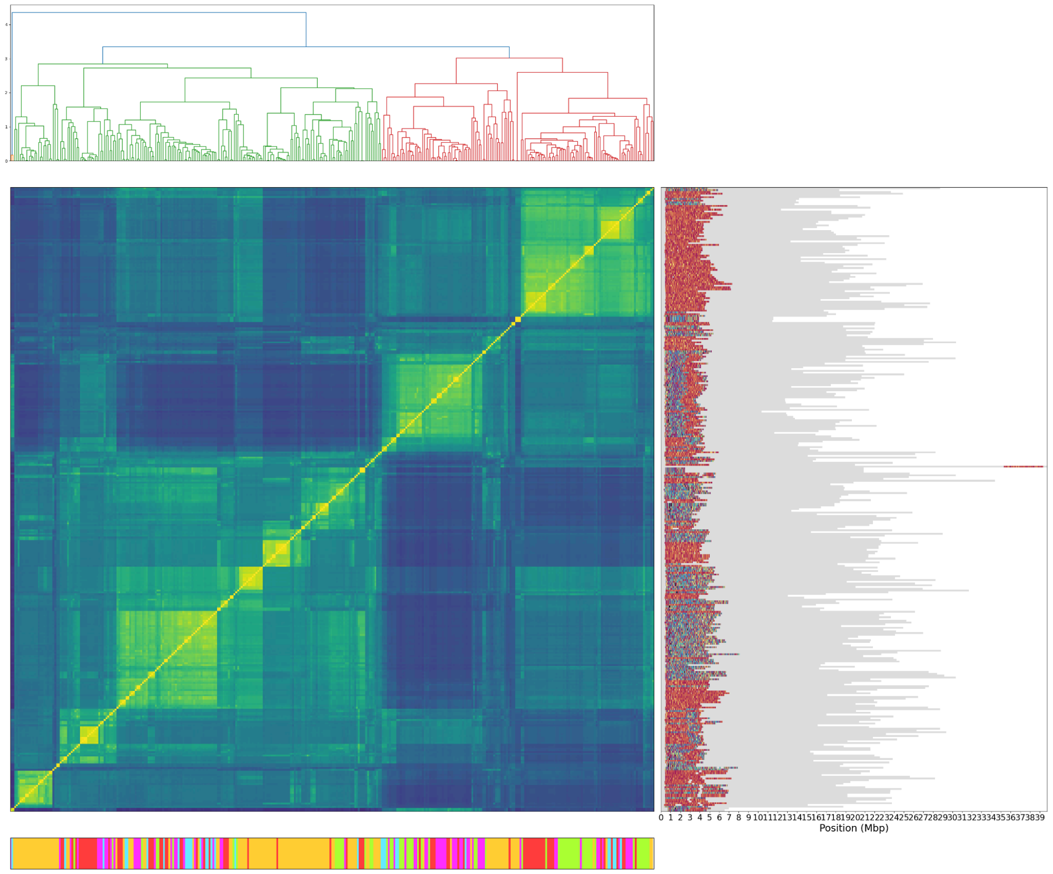

**B** chr2

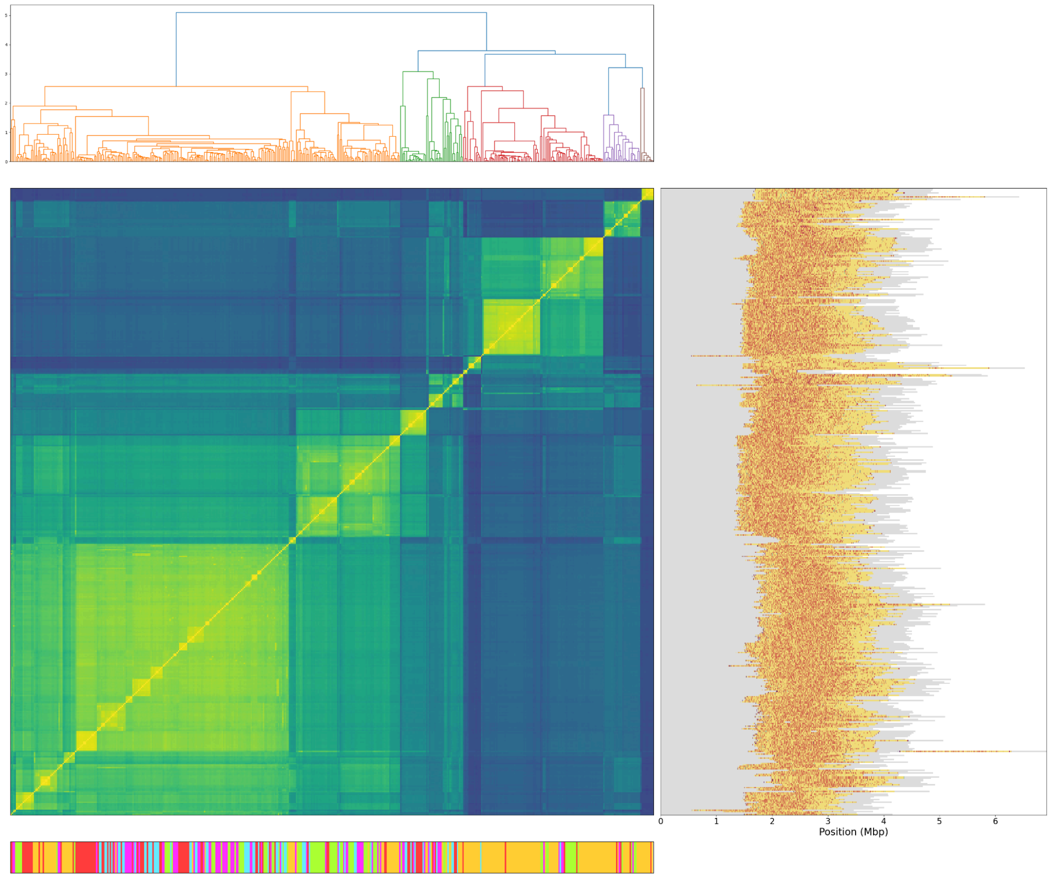


**C** chr3

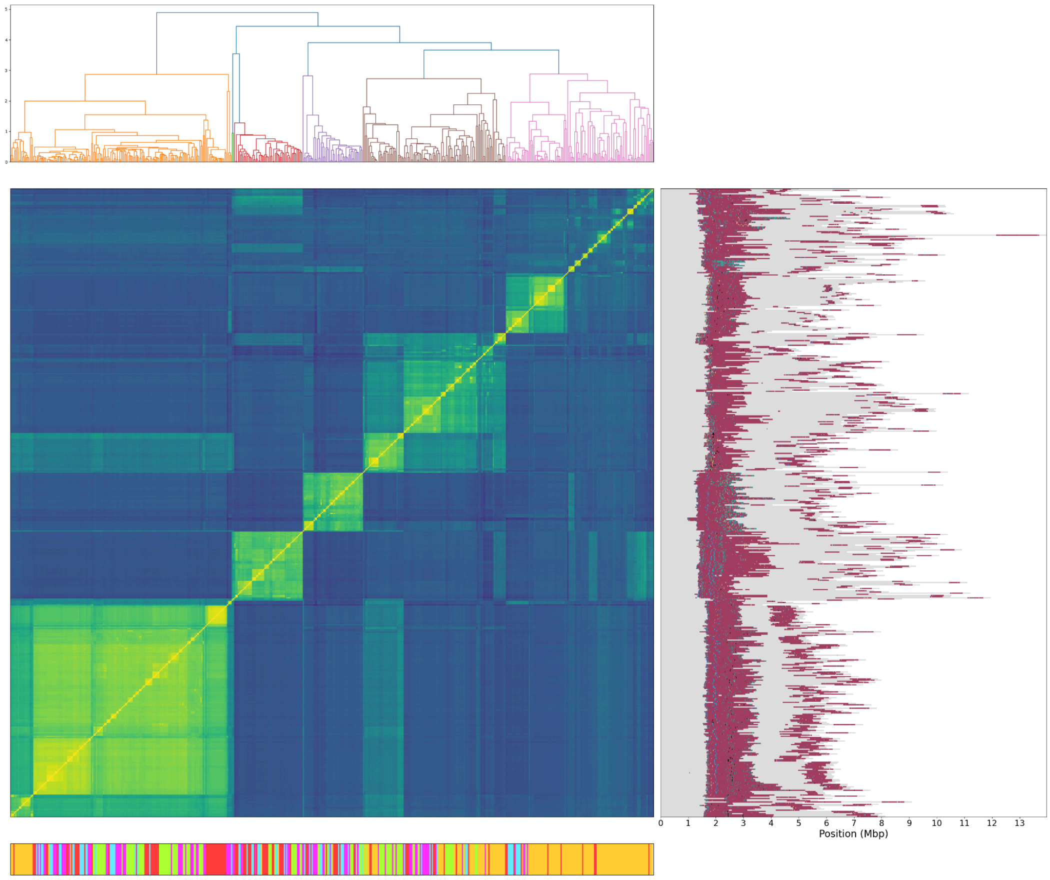

**D** chr4

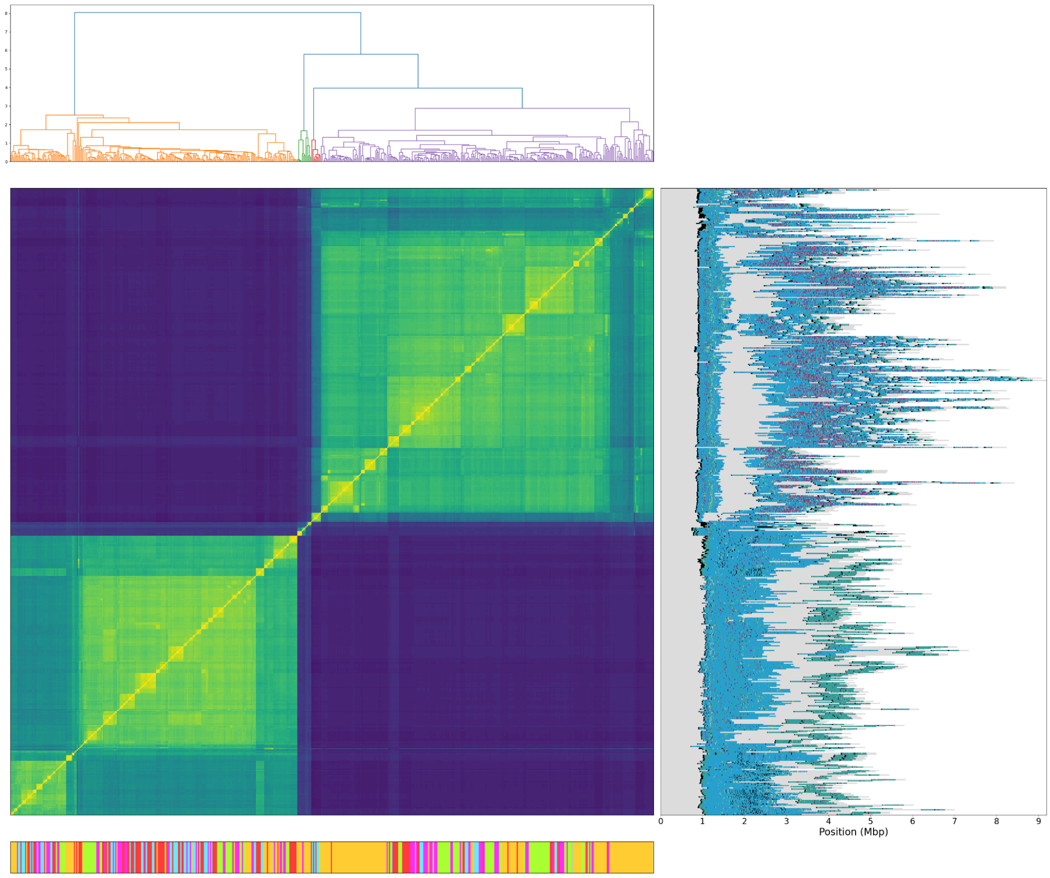


**E** chr5

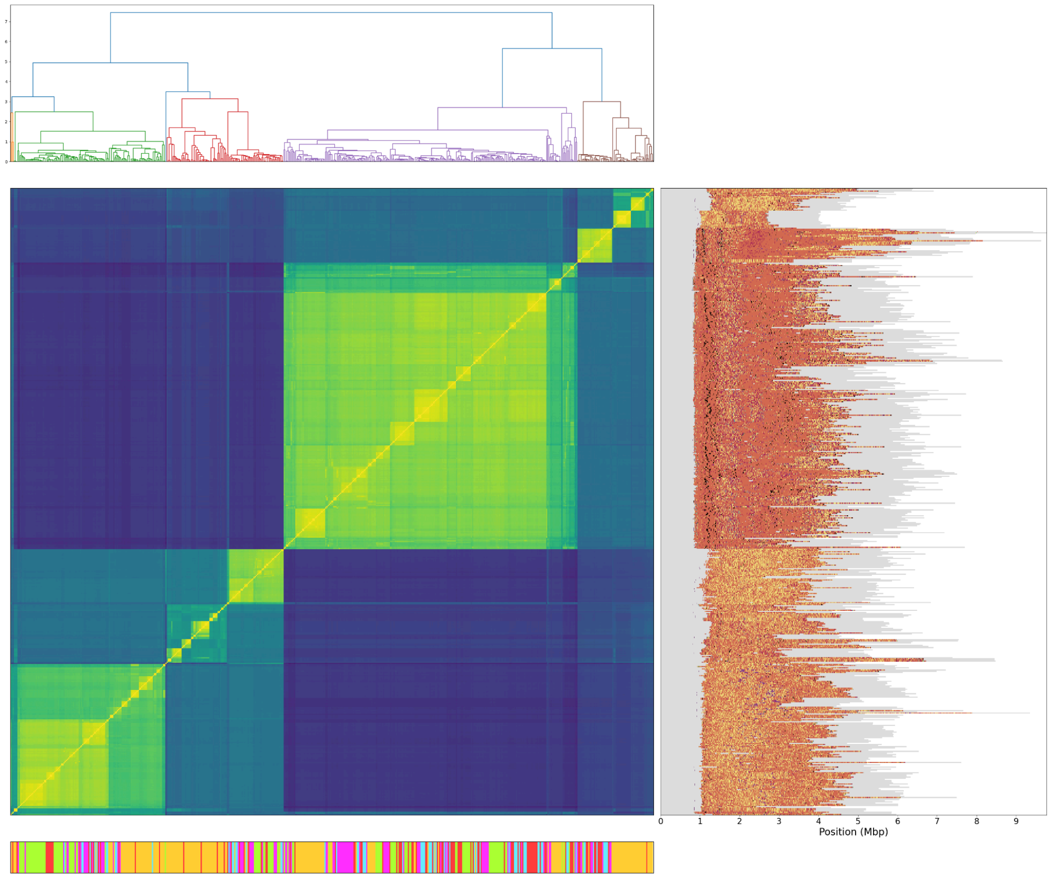

**F** chr6

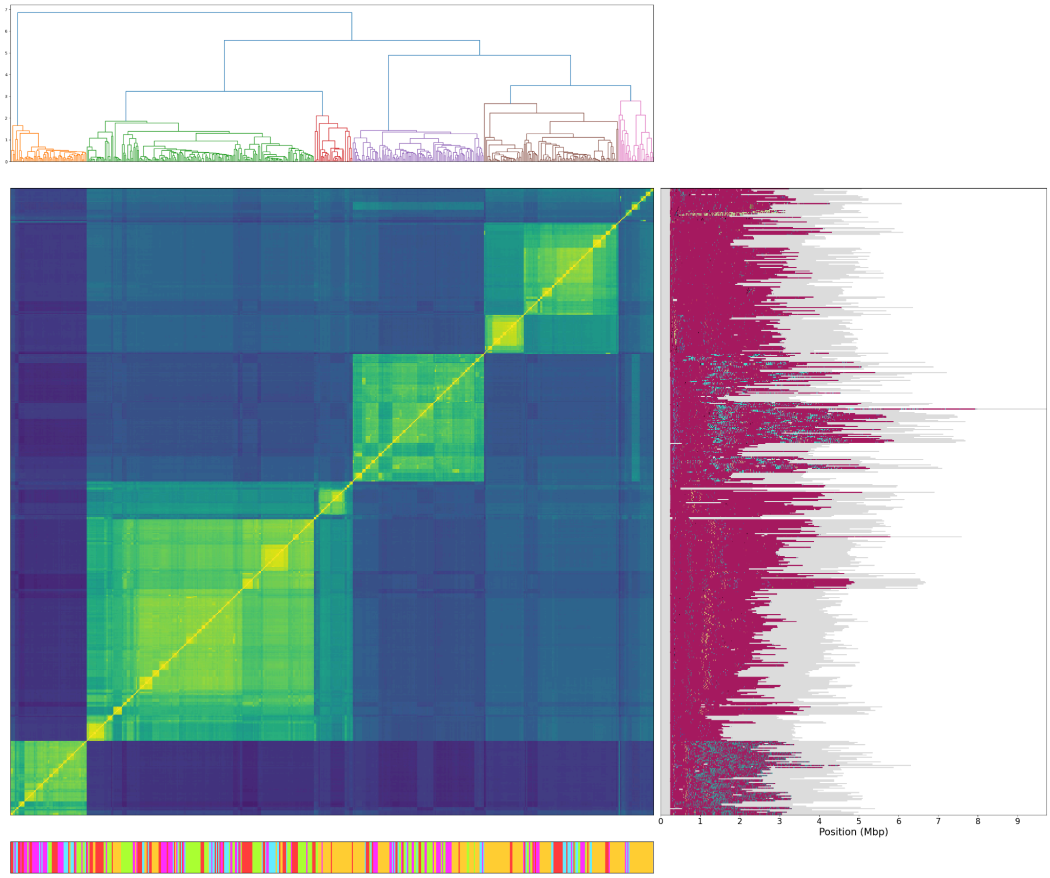


**G** chr7

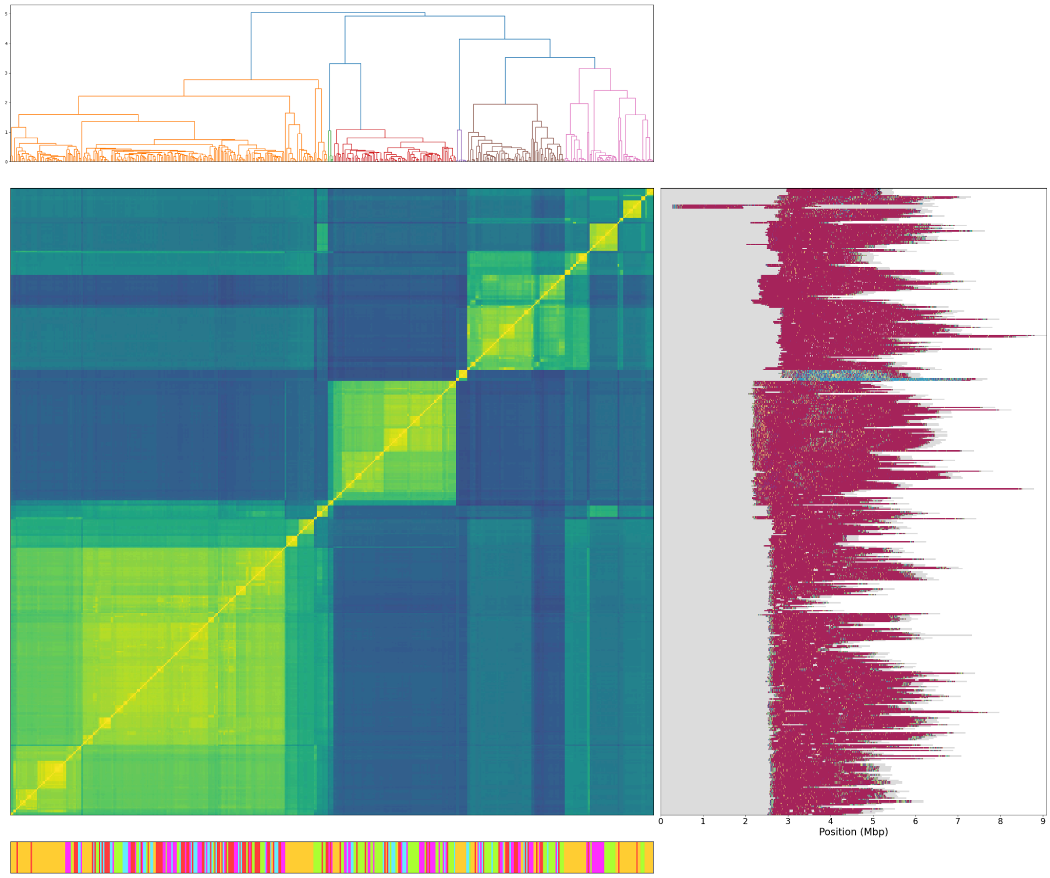

**H** chr8

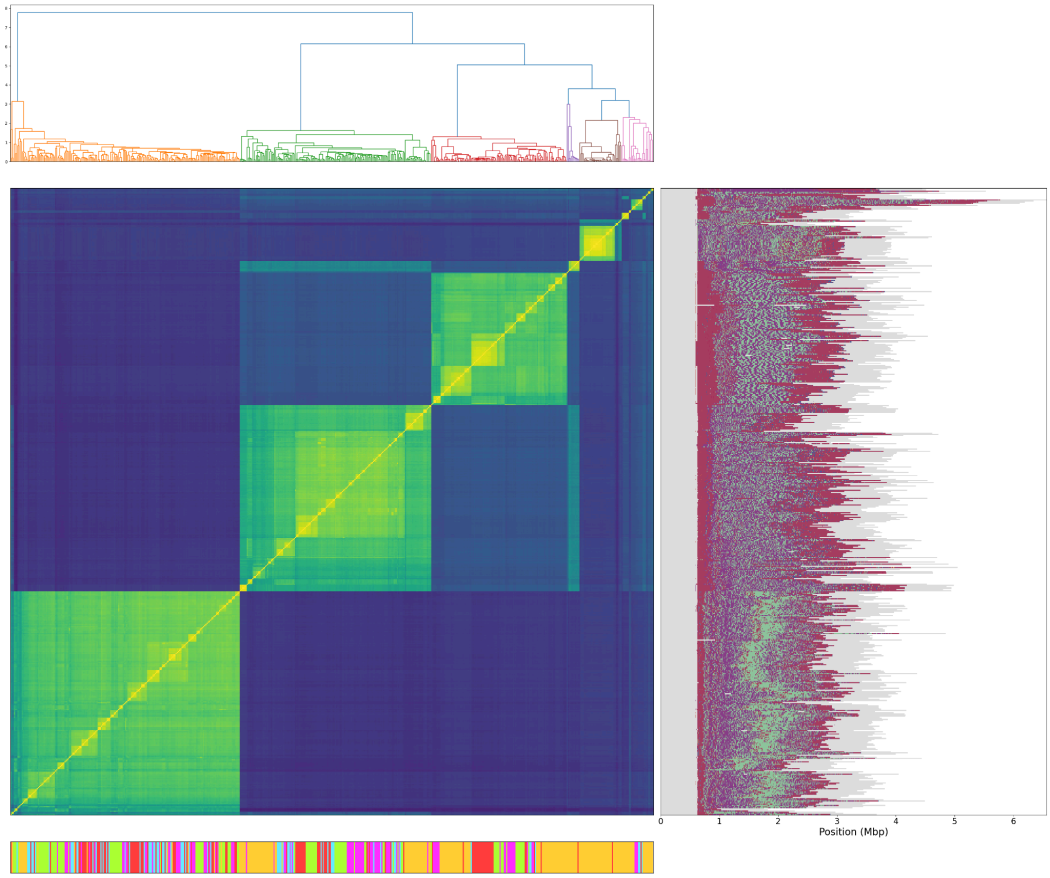


**I** chr9

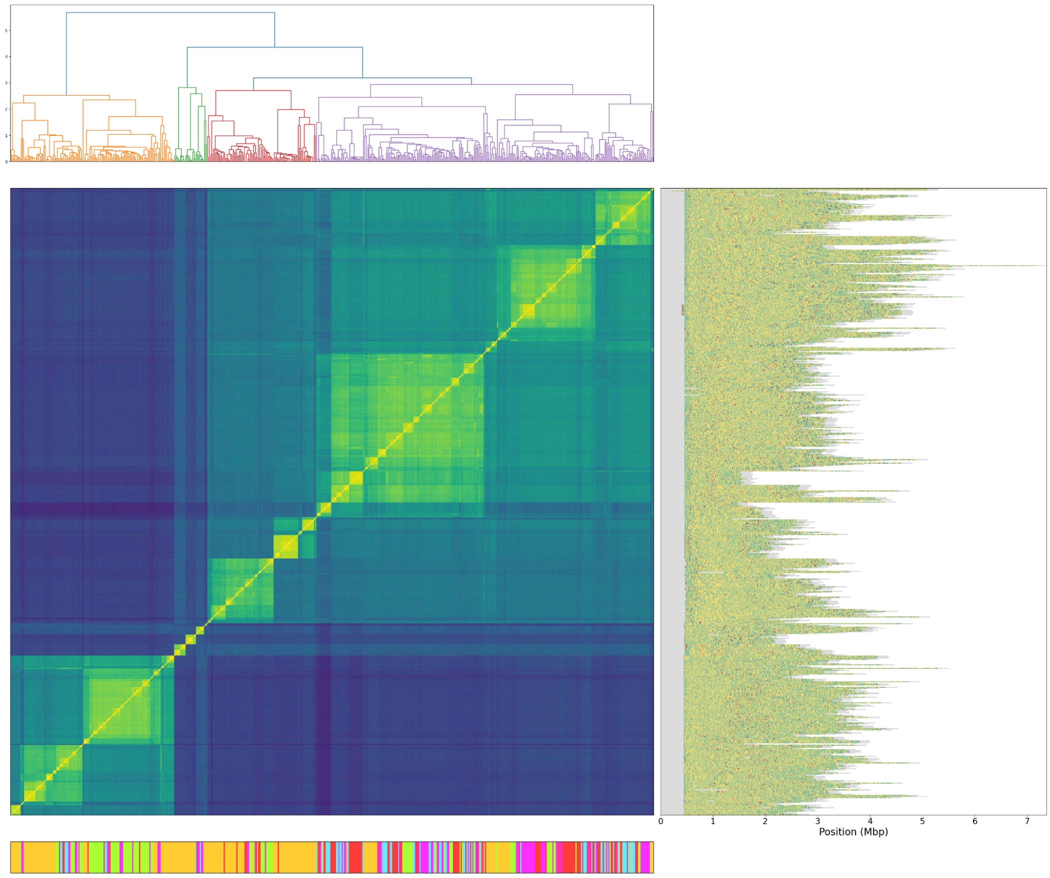

**J** chr10

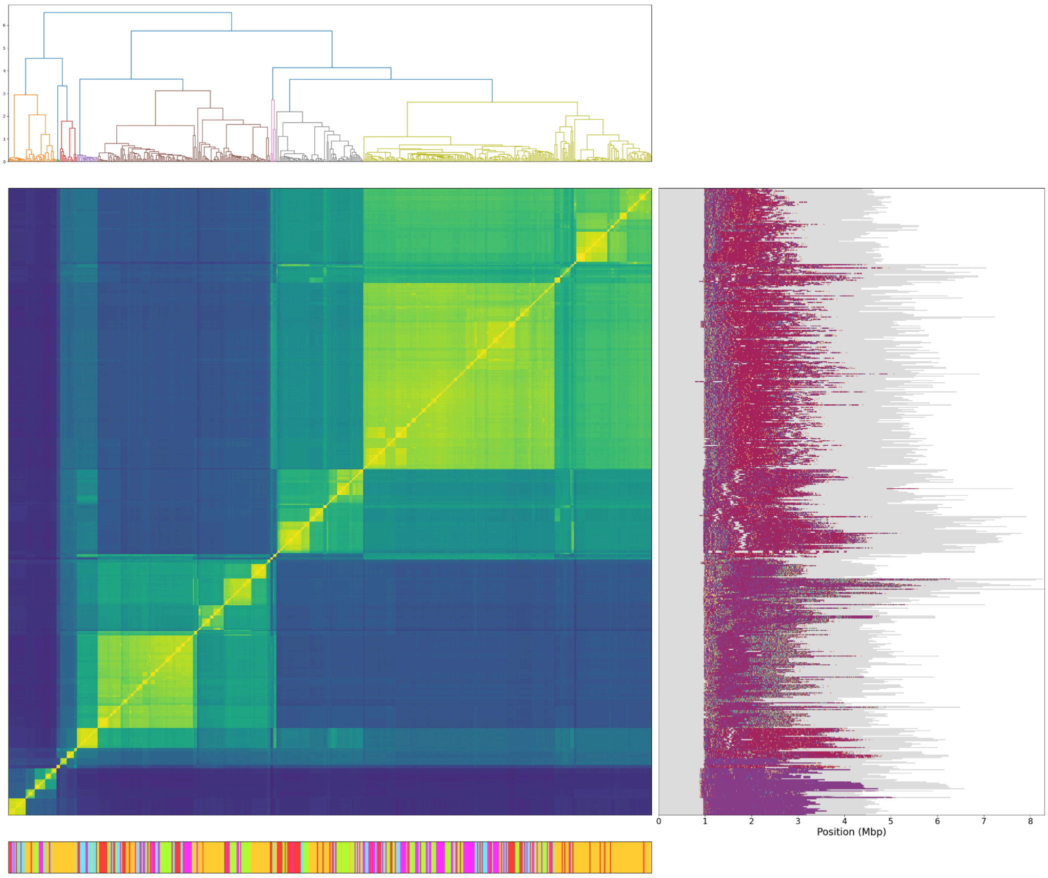


**K** chr11

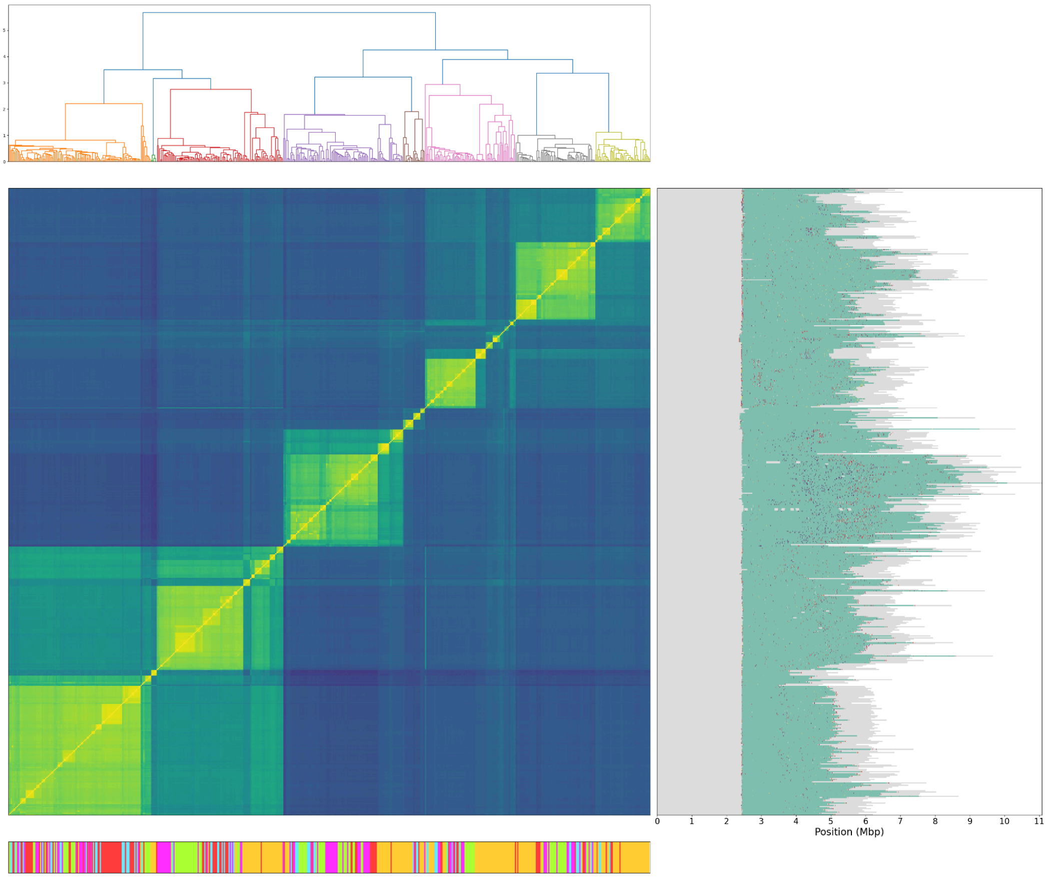

**L** chr12

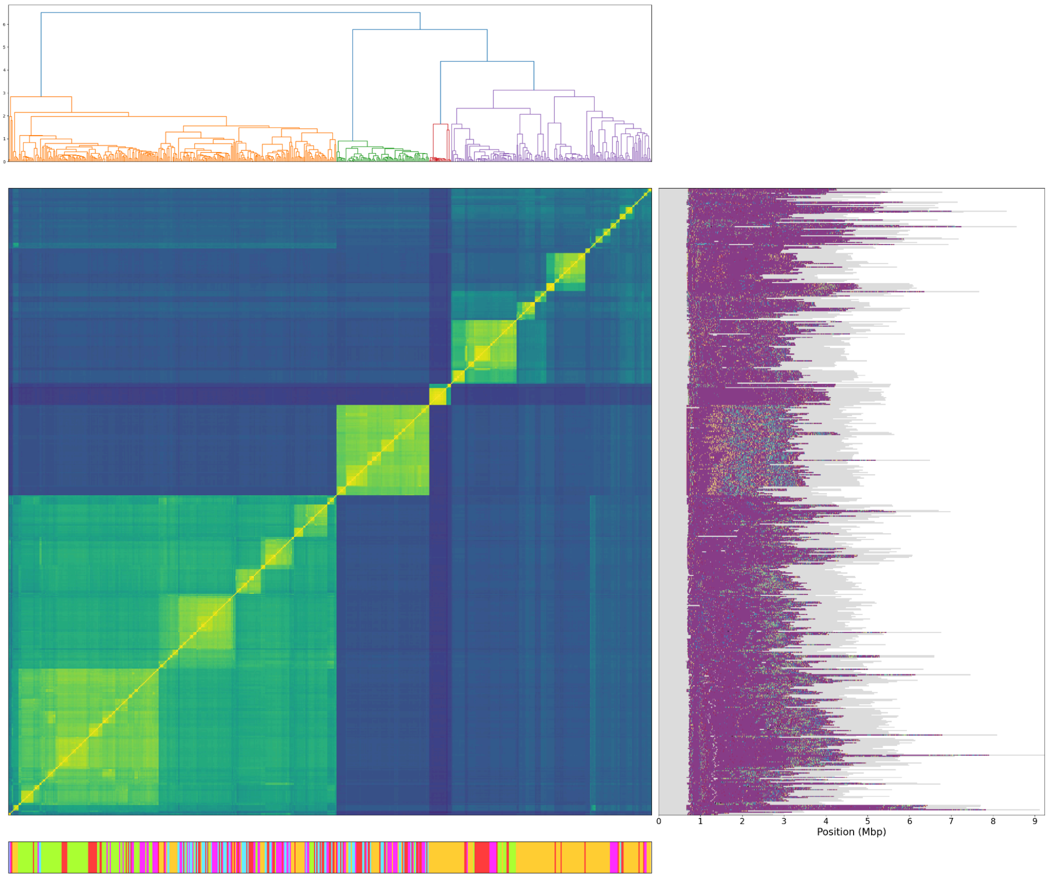


**M** chr13

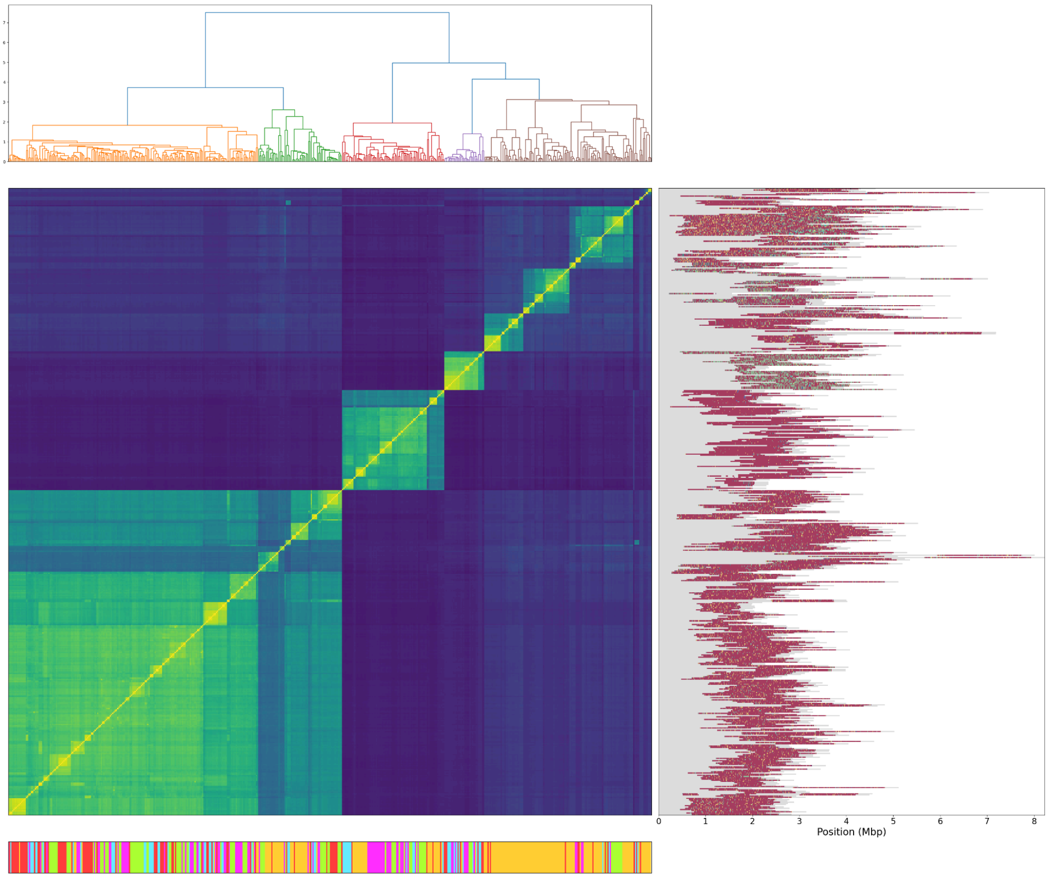

**N** chr14

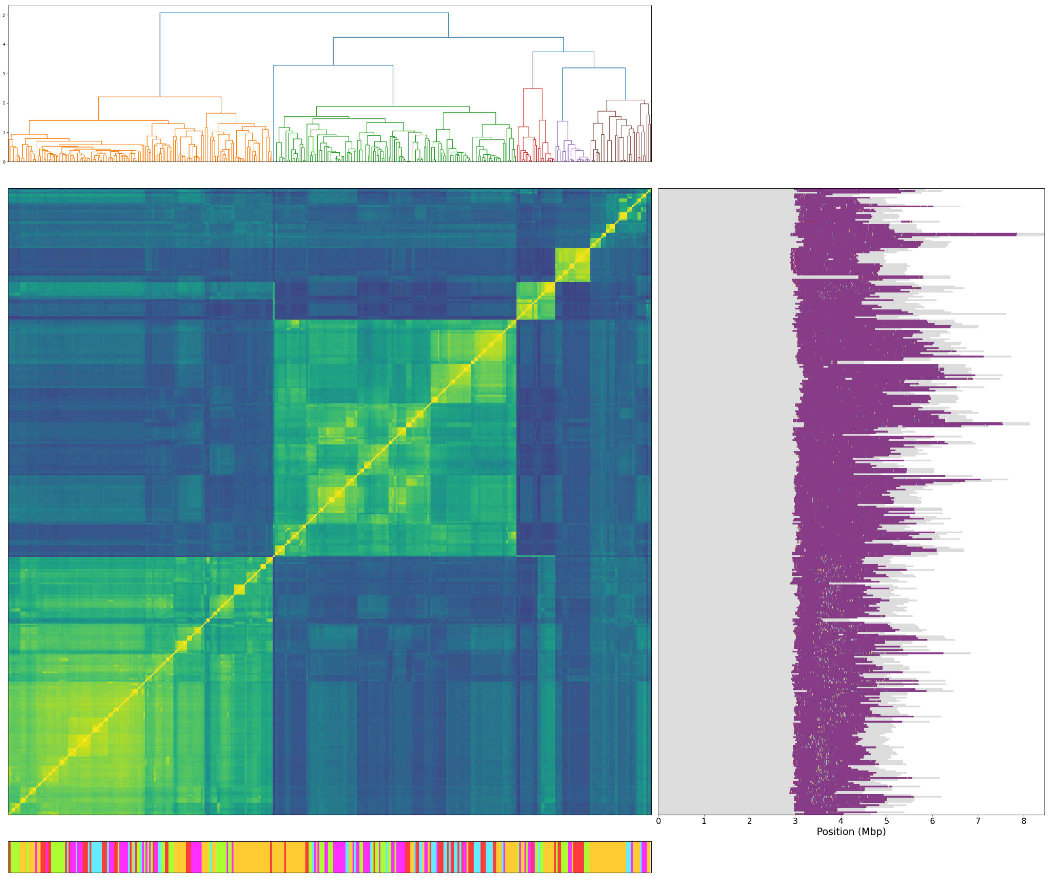


**O** chr15

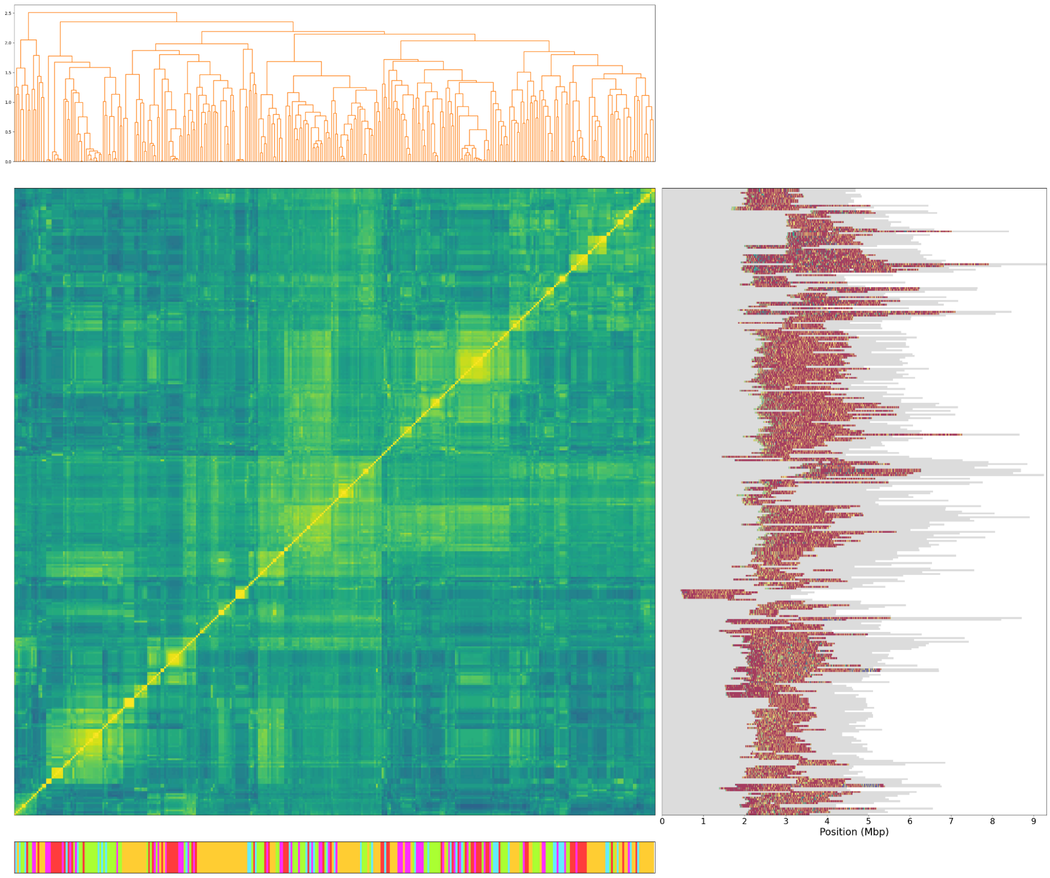

**P** chr16

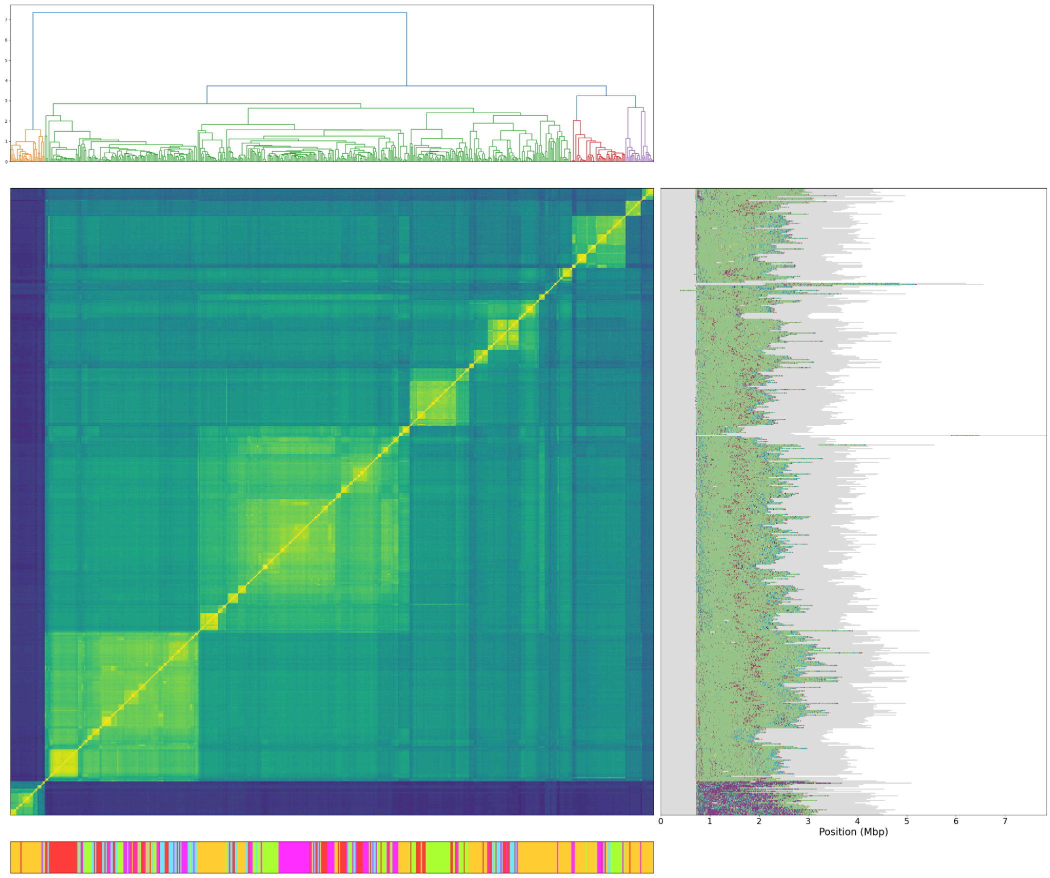


**Q** chr17

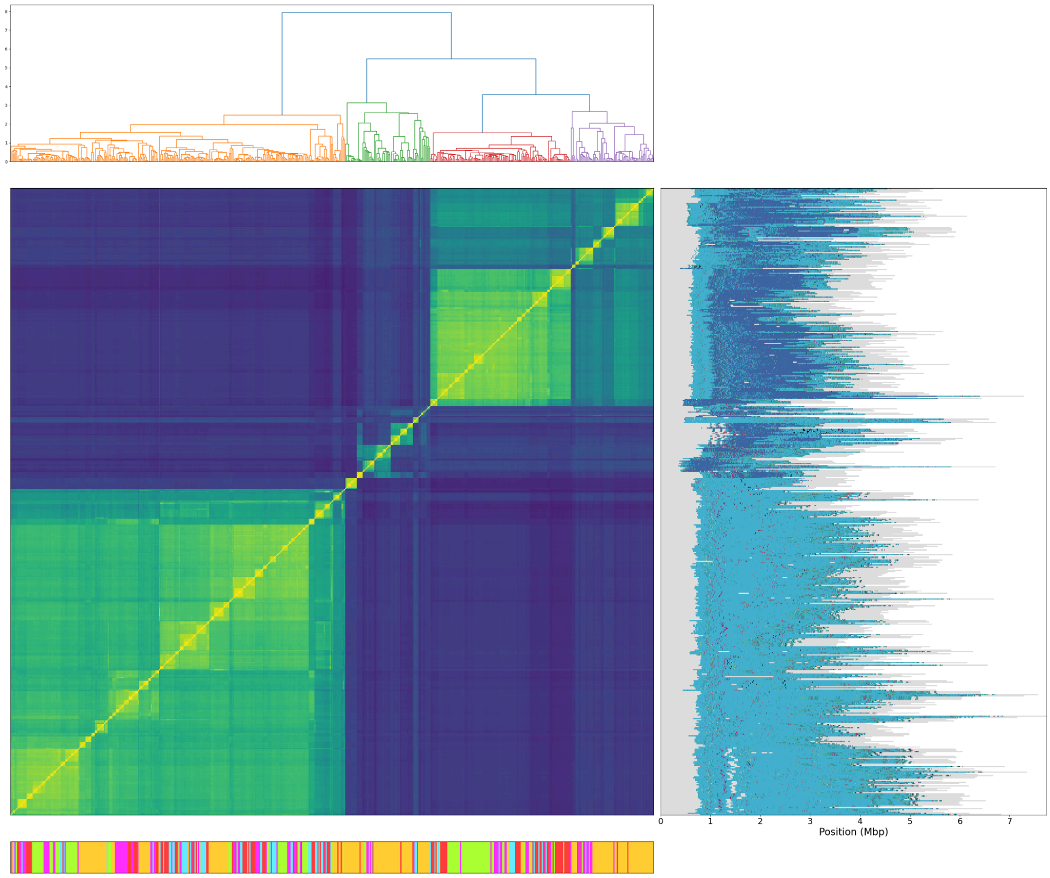

**R** chr18

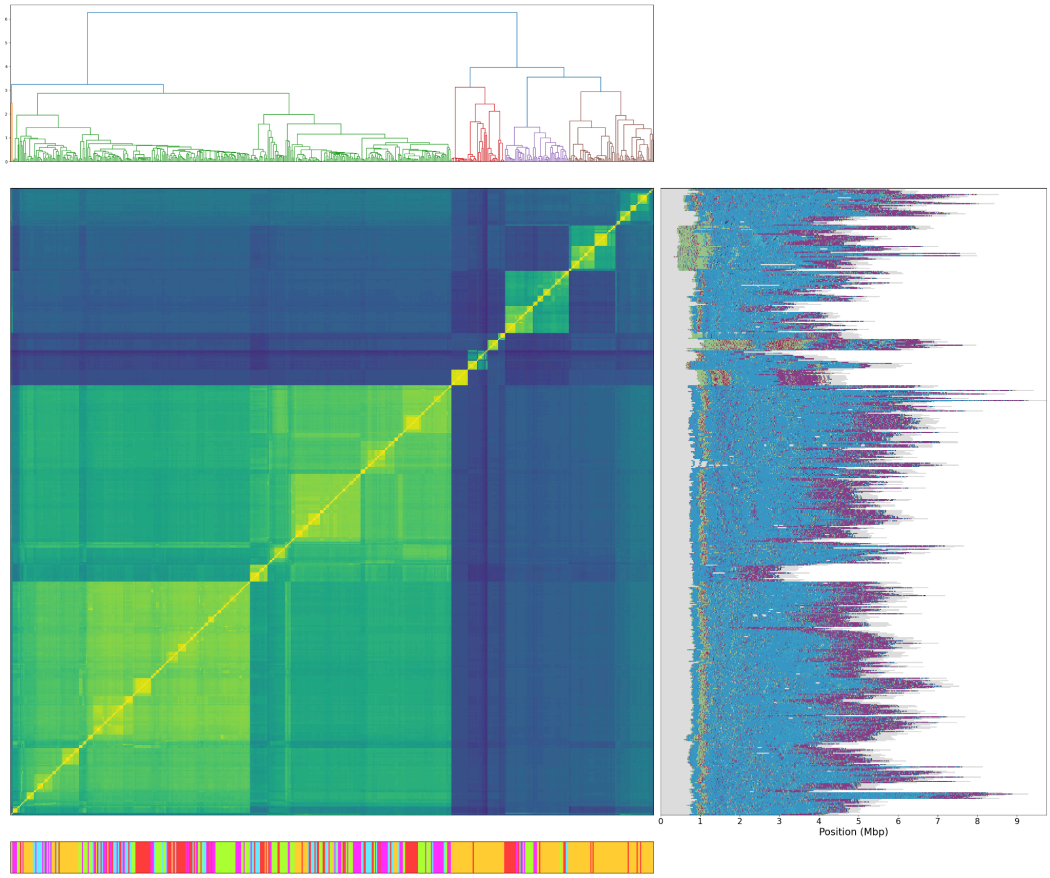


**S** chr19

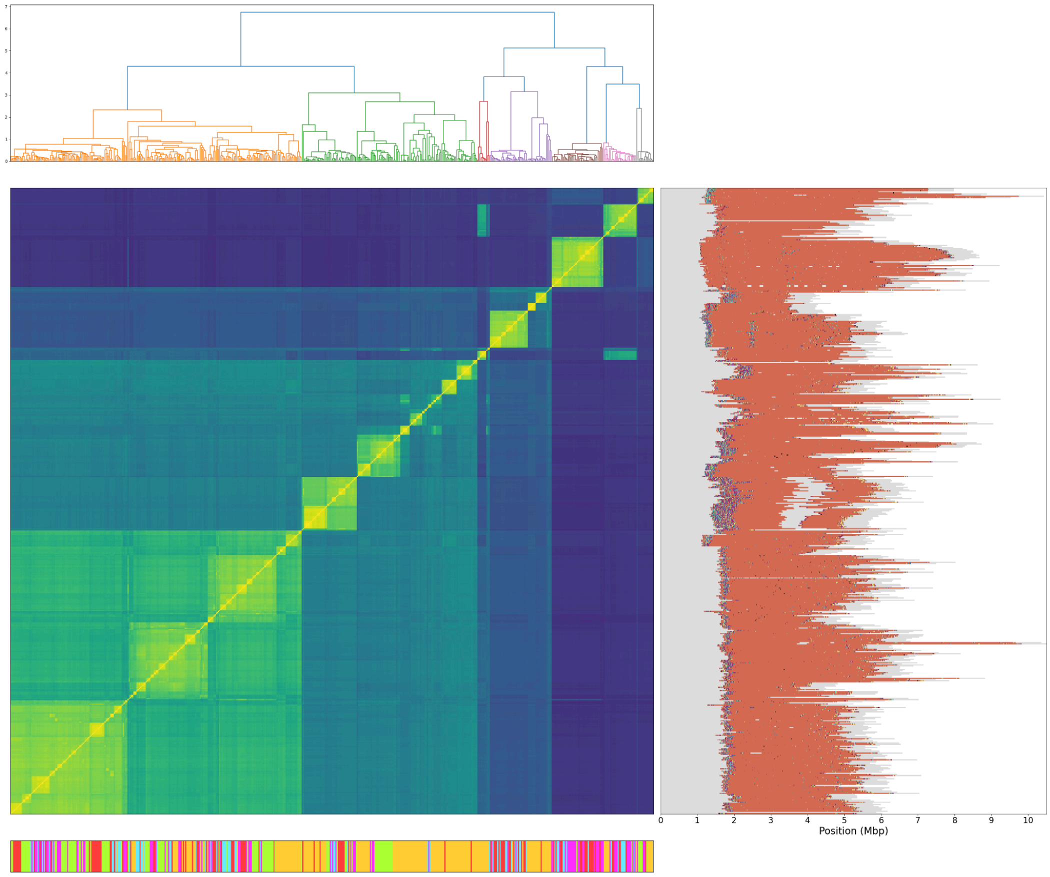

**T** chr20

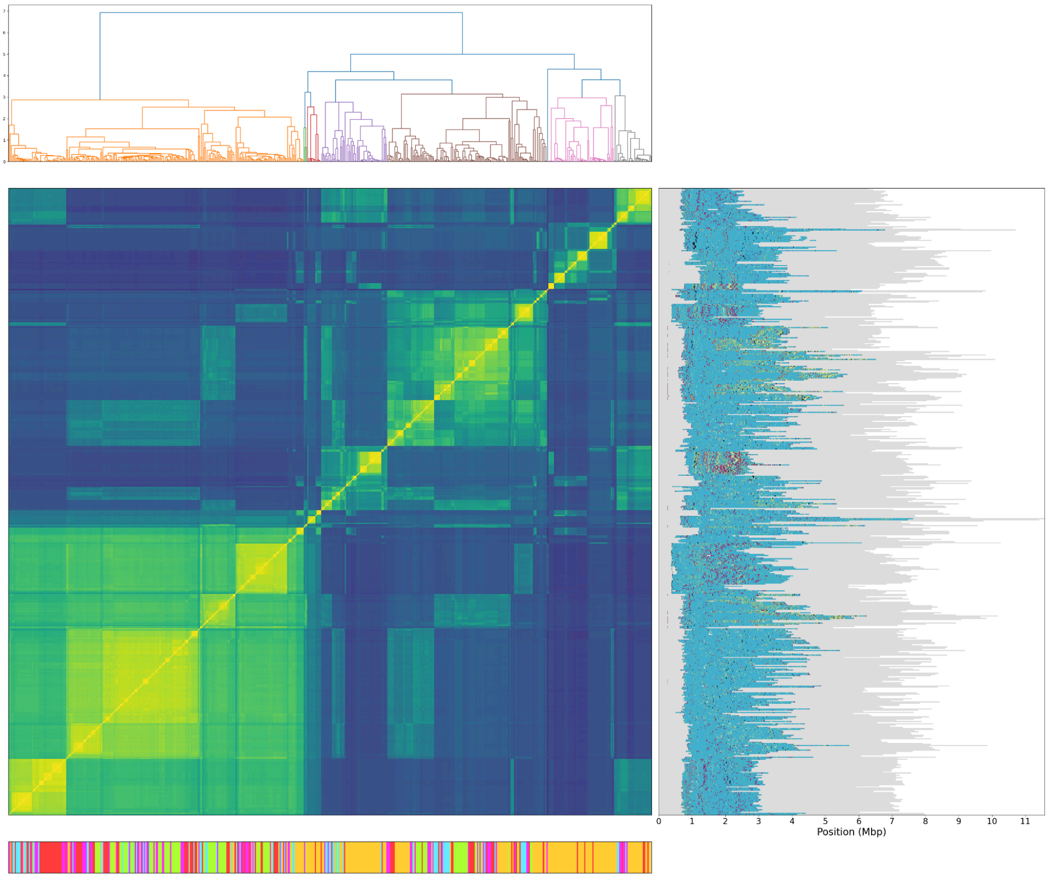


**U** chr21

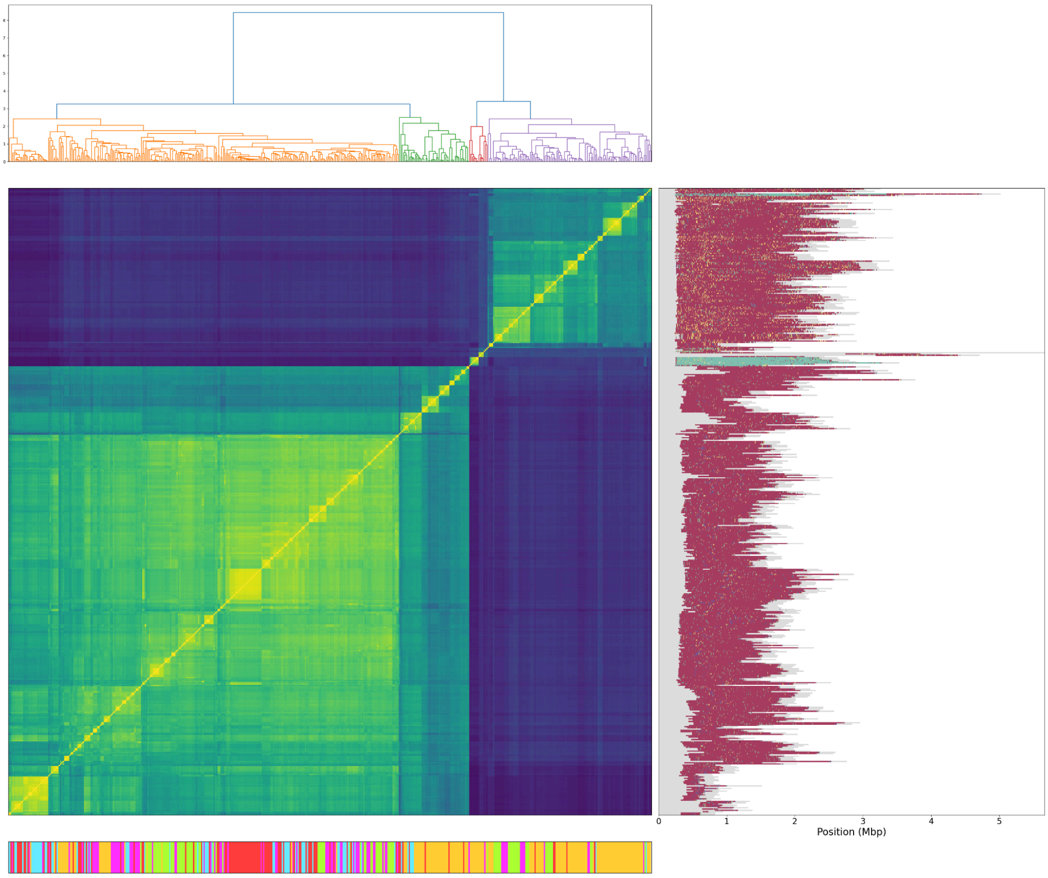

**V** chr22

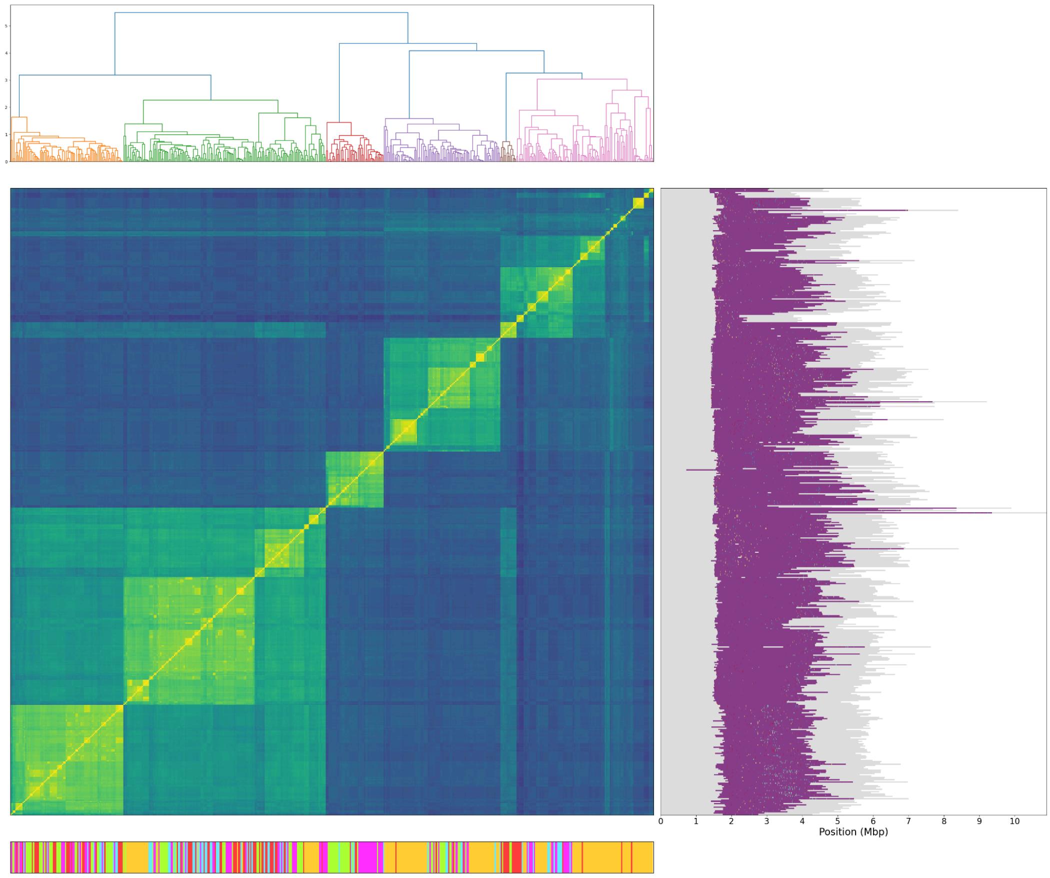


**W** HOR legend

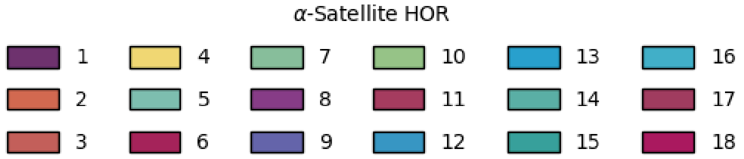

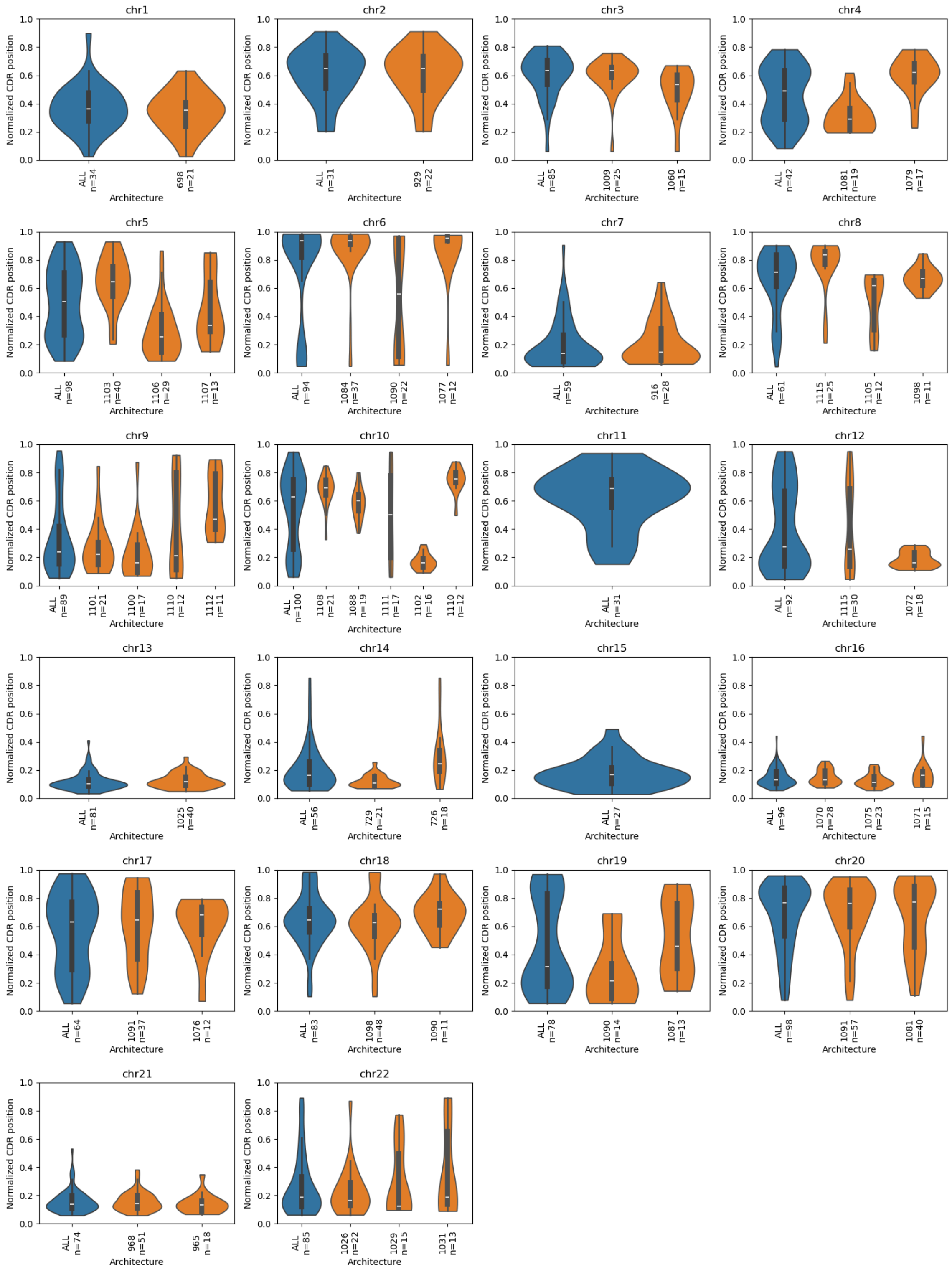


**Figure S4: Distribution of CDR position for different chromosomes stratified by architecture.**
In complete centromere assemblies with published Nanopore sequencing data the position of the demethylated CDR region was determined. Shown here is the relative position of the CDR inside the HORs extracted from T2T assemblies. The data is shown for all assemblies of one chromosome (blue) and also stratified into the identified centromere architectures (orange). The sample number is indicated and only architectures with 10 or more samples are shown.

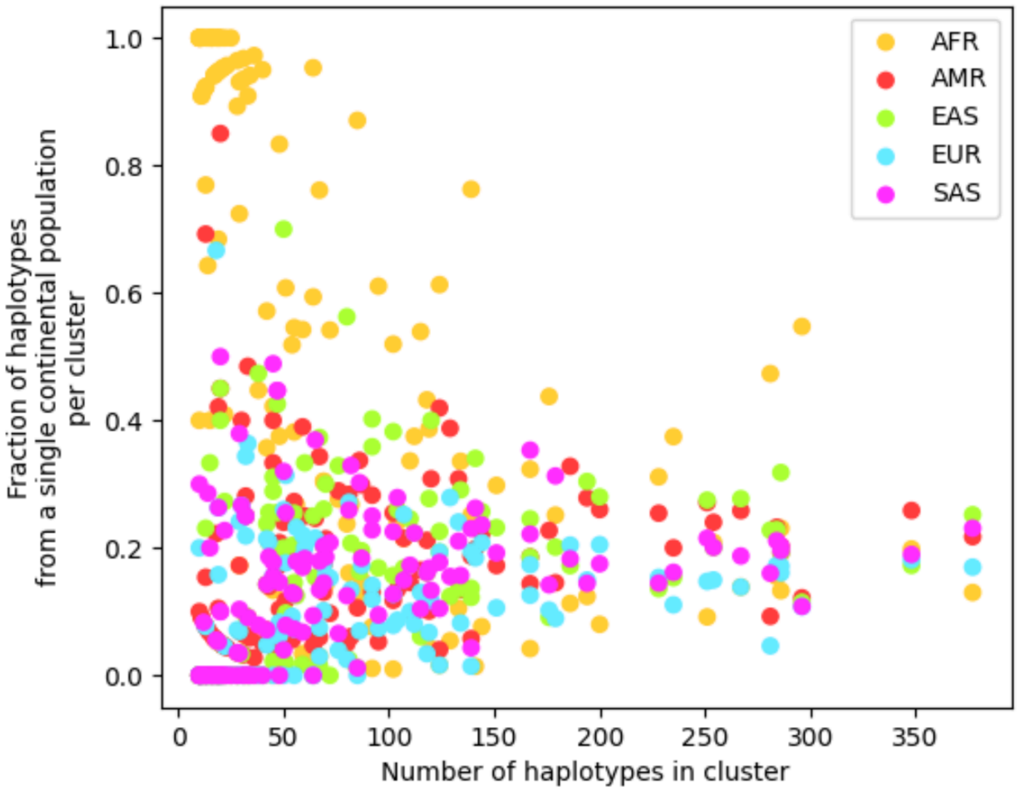


**Figure S5: Centromere clustering reveals several small to medium-sized clusters enriched for haplotypes of African descent - an effect not observed for haplotypes from other continental populations.**

For each initial cluster derived from the phylogenetic tree, the total number of haplotypes and the contribution of each continental population were quantified and plotted. Non-African haplotypes typically comprise 10-30% of the haplotypes within a given cluster. In contrast, African haplotypes can be found in small to medium-sized clusters where they are highly enriched and contribute more than half of all haplotypes. This enrichment diminishes in larger clusters, where the representation of all continental populations becomes more uniform.

**Figure S6: Nodes in phylogenetic trees of centromeres with architecture-specific k-mers.**

Based on the phylogenetic tree of all centromeres from a single chromosome (**A–V**), all nodes with branch lengths between 1.2 and 10 were queried for architecture-specific *k*-mers. Nodes where architecture-specific *k*-mers were found are marked with a circle. The size of each circle corresponds to the number of *k*-mers identified, while the circle's color indicates the maximum mean copy number of any architecture-specific *k*-mer at that node (see legend in panel **W**). Selected architectures used for downstream prediction are marked with a diamond. These nodes represent a compromise between high resolution of different architectures and possible taggability of a wide range of haplotypes (see **Methods**).

**A** chr1

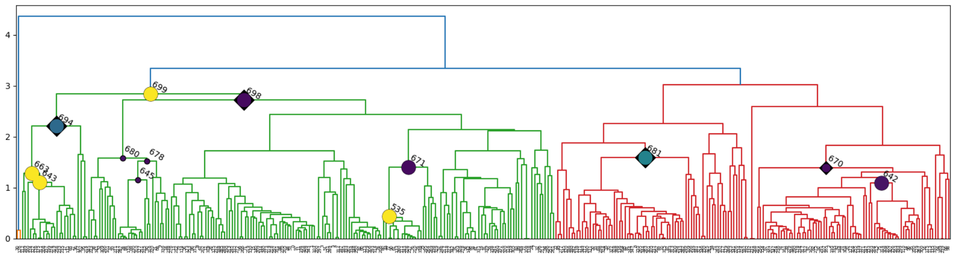

**B** chr2

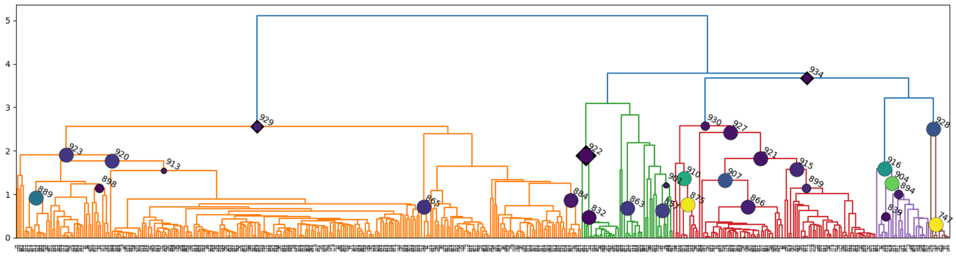

**C** chr3

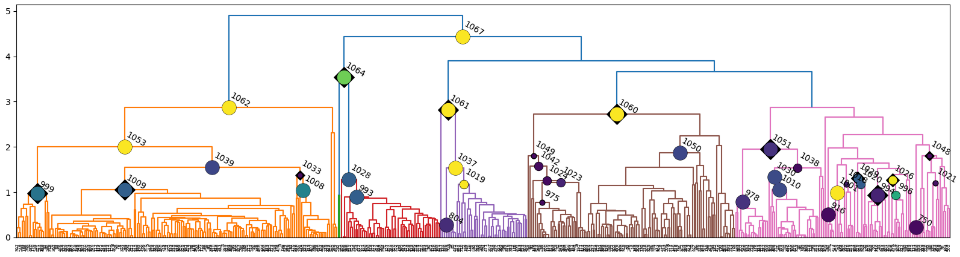

**D** chr4

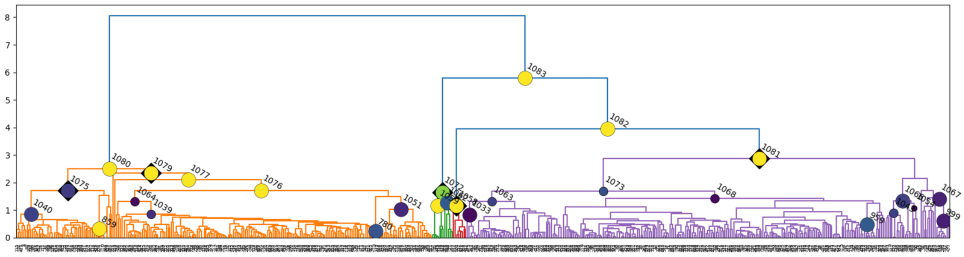

**E** chr5

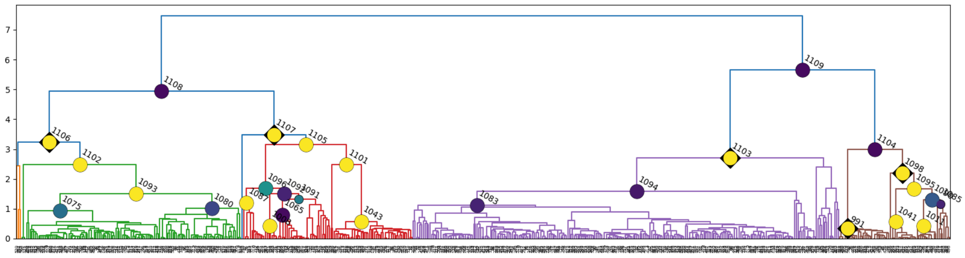

**F** chr6

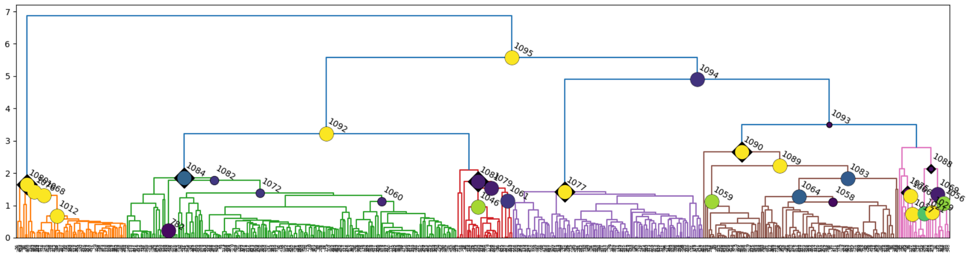

**G** chr7

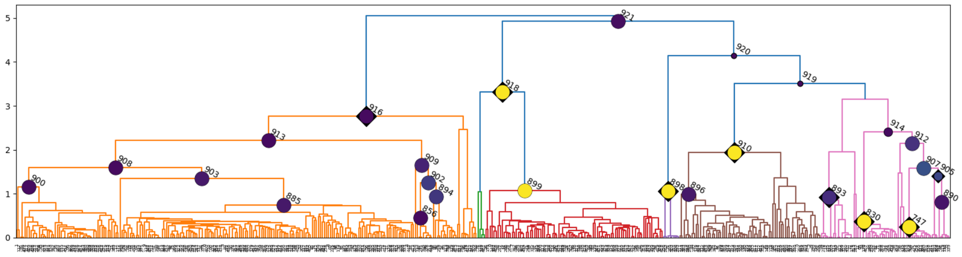

**H** chr8

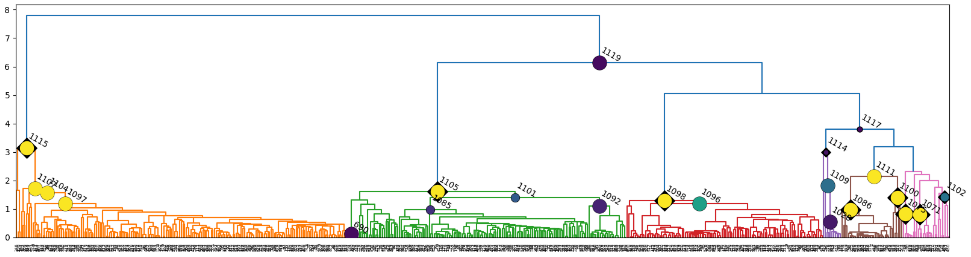

**I** chr9

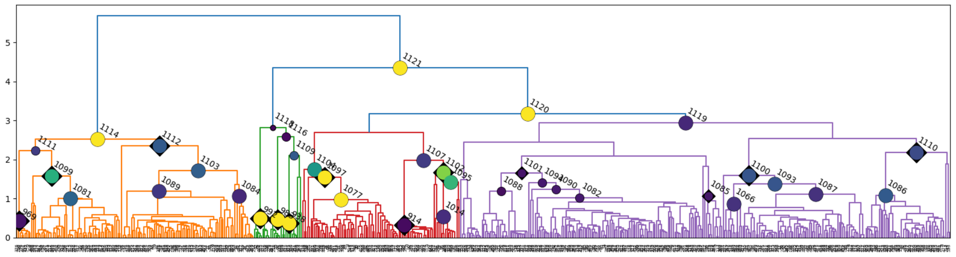

**J** chr10

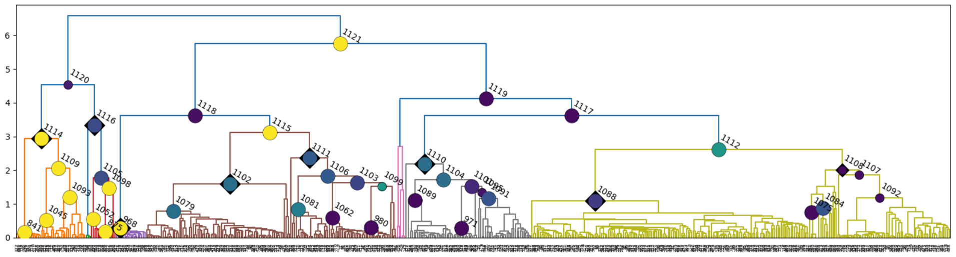

**K** chr11

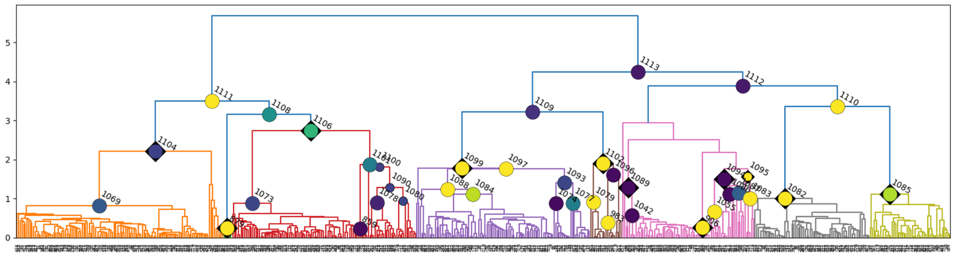

**L** chr12

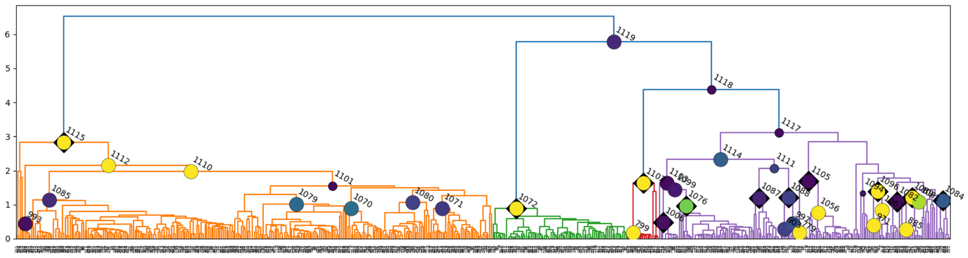

**M** chr13

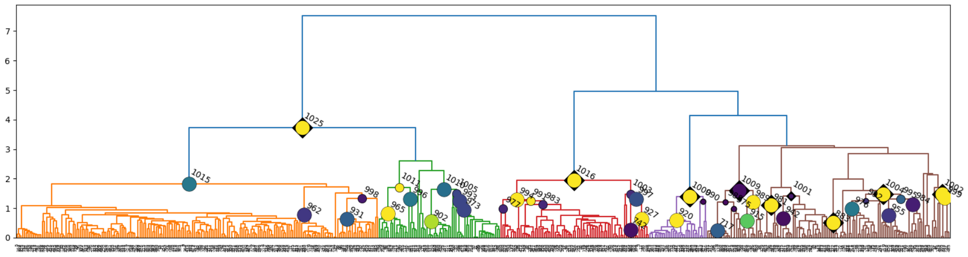

**N** chr14

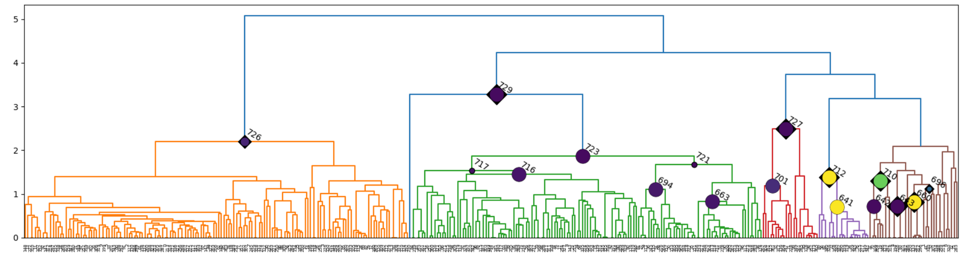

**O** chr15

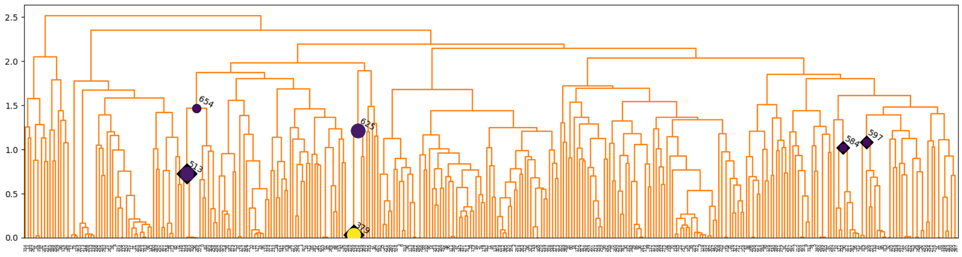

**P** chr16

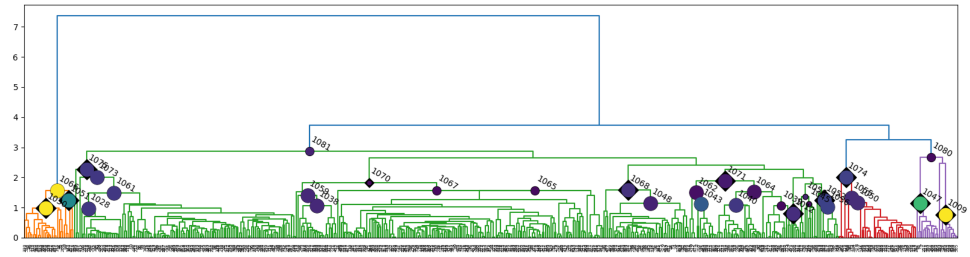

**Q** chr17

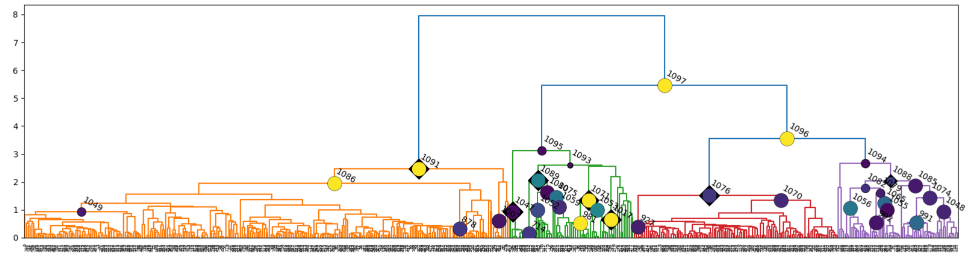

**R** chr18

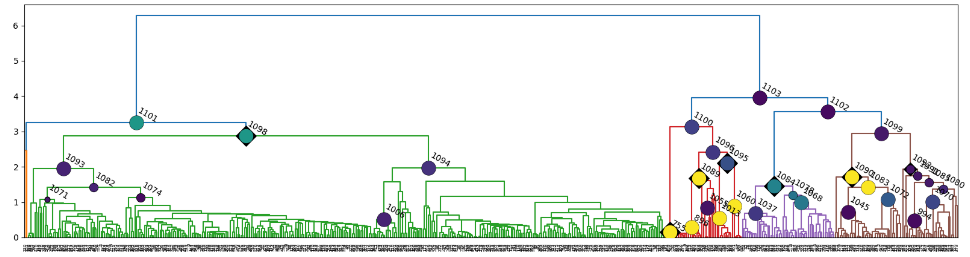

**S** chr19

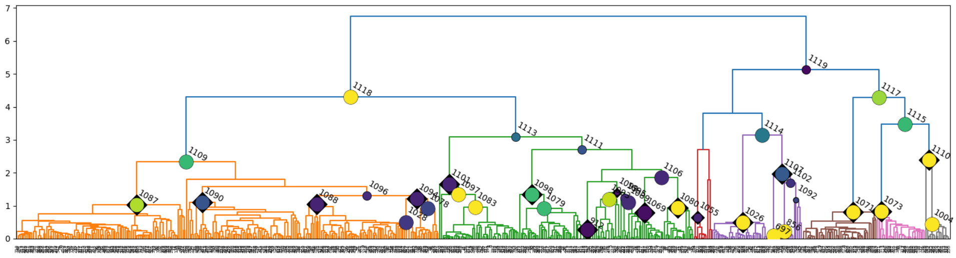

**T** chr20

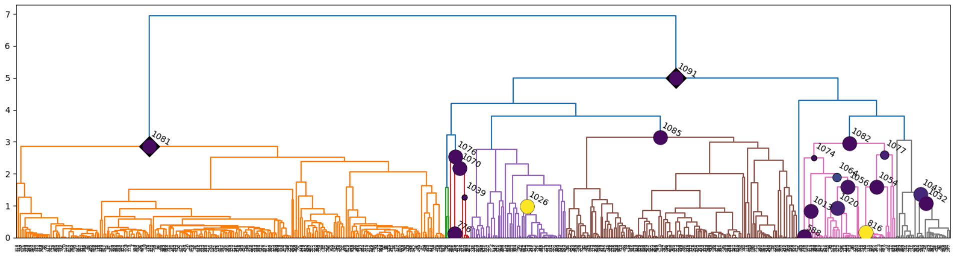

**U** chr21

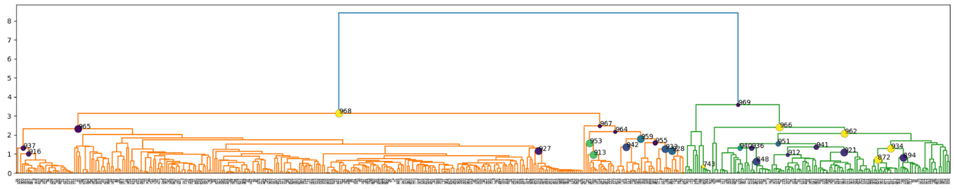

**V** chr22

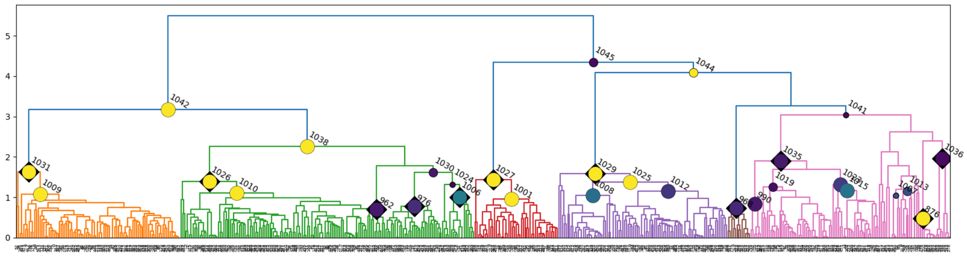

**W** Figure legend

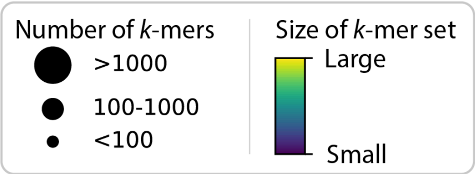

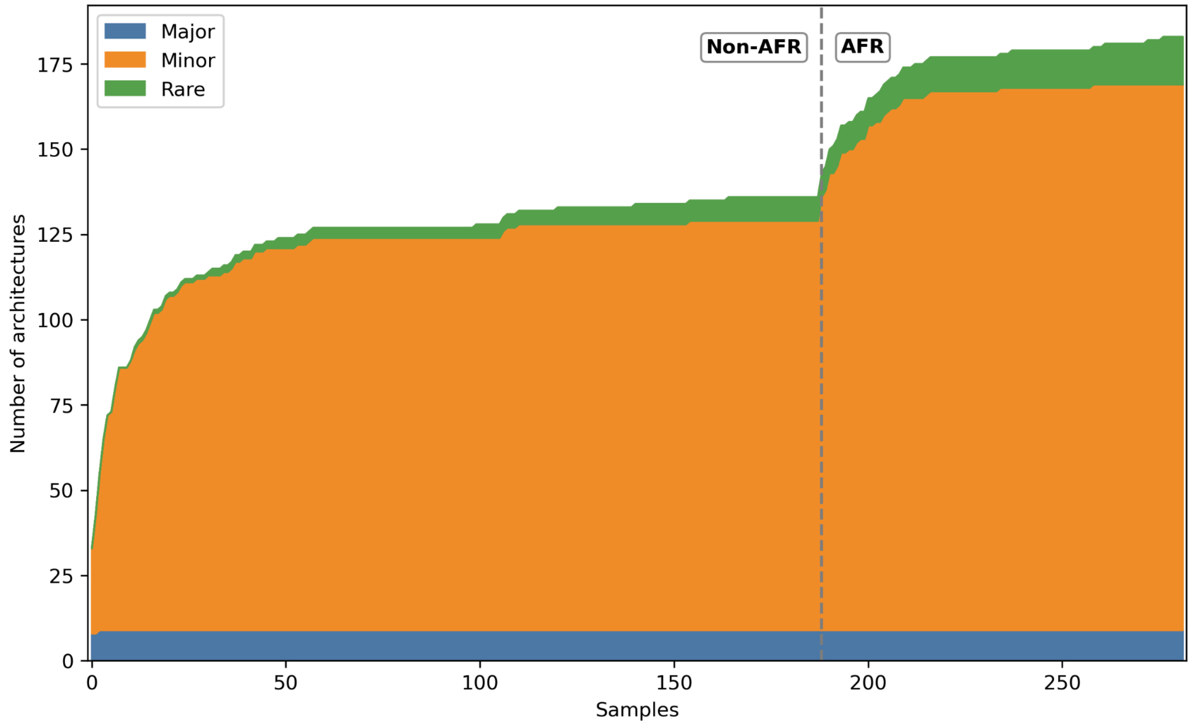


**Figure S7: Cumulative centromere architecture counts across assembled centromere haplotypes.**

Cumulative number of identified centromere architectures detected after sequentially adding samples with near T2T assemblies. Samples are ordered by continental ancestry, with non-African samples shown first and African samples added to the right of the dashed vertical line. Architectures are classified as major (blue), minor (orange), or rare/untagged (green). Major and minor architectures are supported by at least five haplotypes and are included in the HOROSCOPE inference framework. Major architectures have allele frequency ≥0.5, whereas minor architectures have allele frequency <0.5. Rare/untagged architectures represent centromere haplotypes with at least one similar haplotype, defined by a *k*-mer similarity >0.95, but with insufficient support for architecture-specific *k*-mer selection (<5 supporting haplotypes), and are therefore not queried and used by HOROSCOPE. Singleton centromere haplotypes without a similar assembly (again defined by a *k*-mer similarity <0.95), were excluded from this analysis (n=186 centromeres, 1.7% of all analyzed centromere haplotypes). Chromosomes 15, X, and Y were excluded as chromosome 15 did not produce extensive architecture clusters, while chromosomes X and Y are in general not included in the HOROSCOPE framework.

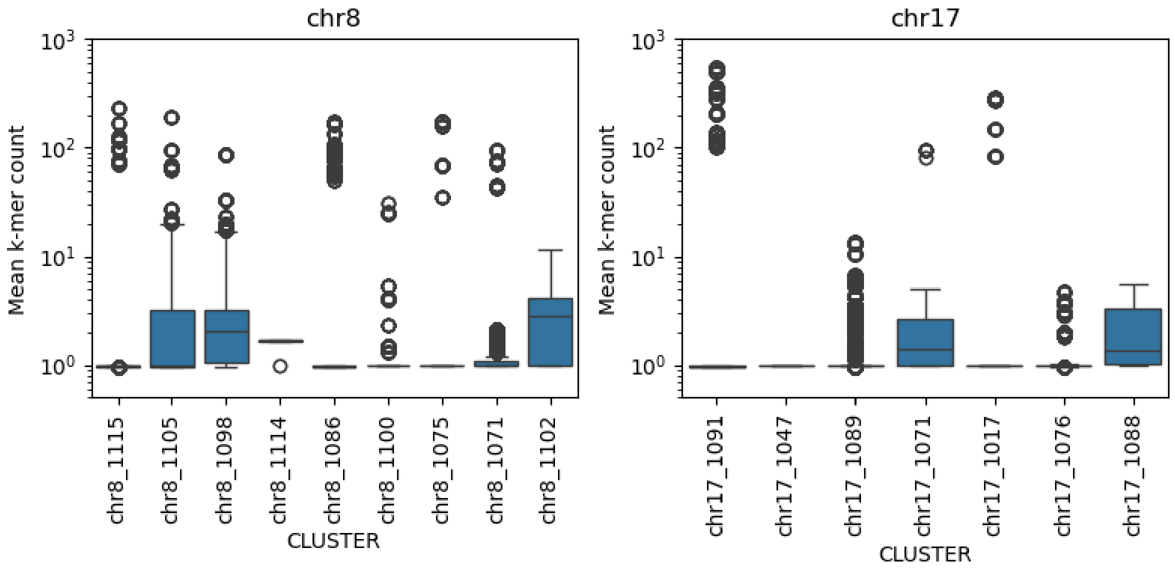


**Figure S8: Mean *k*-mer counts of architecture-specific *k*-mers identified in assemblies of carrier haplotypes.**

For each architecture of chr8 (left) and chr17 (right), the distribution of mean *k*-mer counts for the corresponding architecture-specific *k*-mers is shown.

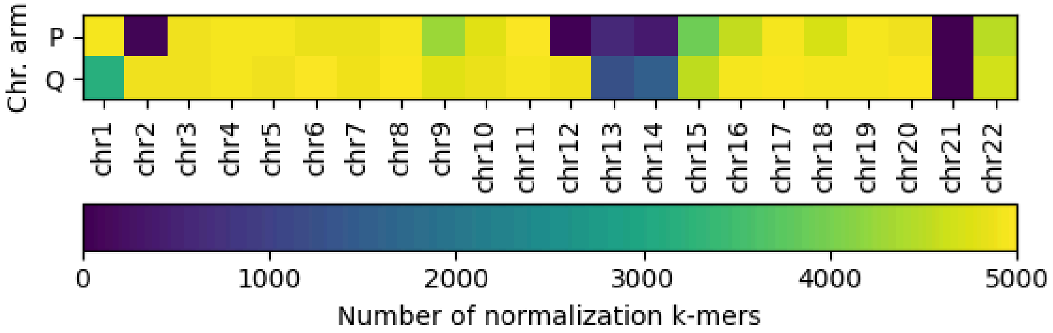


**Figure S9: Number of normalization *k*-mers for the p- and q-arm flank of the centromeres of each chromosome.**

Initially 5000 potential *k*-mers per flank with a copy-number of 1 in each centromere assembly of one chromosome were randomly chosen. The normalization kmer sets were further filtered to remove *k*-mers present in other centromeres or in other locations of the CHM13 genome. No normalization *k*-mers were found for chr21 and only 75 and 29 normalization *k*-mers were identified for chr2p and chr12p respectively. For chr13 and chr14 at least 383 and 1286 normalization *k*-mers were found for the p- and q-arm.

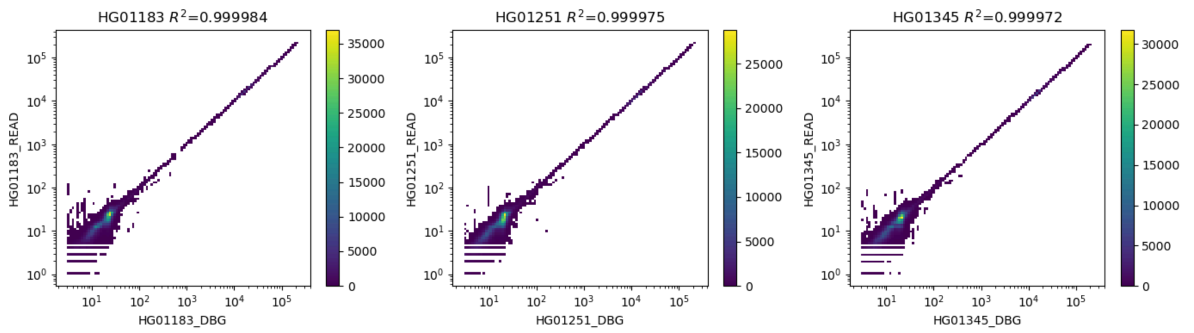


**Figure S10: Correlation of *k*-mer counts derived from raw sequencing reads and from the de Bruijn graph (DBG) assembly for centromere-related *k*-mers (including architecture-specific, length-informative, and normalization *k*-mers) across three samples.**

Each panel represents one sample; the x-axis shows *k*-mer counts from the DBG, and the y-axis shows *k*-mer counts from raw reads.

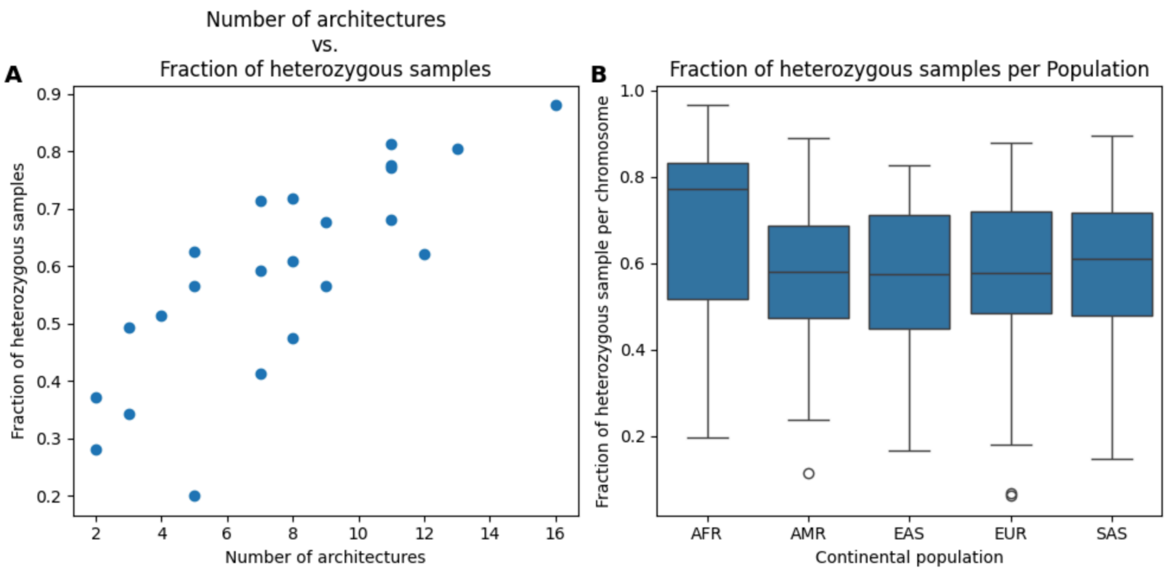


**Figure S11: Fraction of heterozygous samples across chromosomes and continental populations.**

(**A**) Scatter plot showing the relationship between the number of distinct centromere architectures and the fraction of heterozygous samples per chromosome. A sample is considered heterozygous if its two centromere alleles derive from different architectures. (**B**) Fraction of heterozygous samples across continental subpopulations.

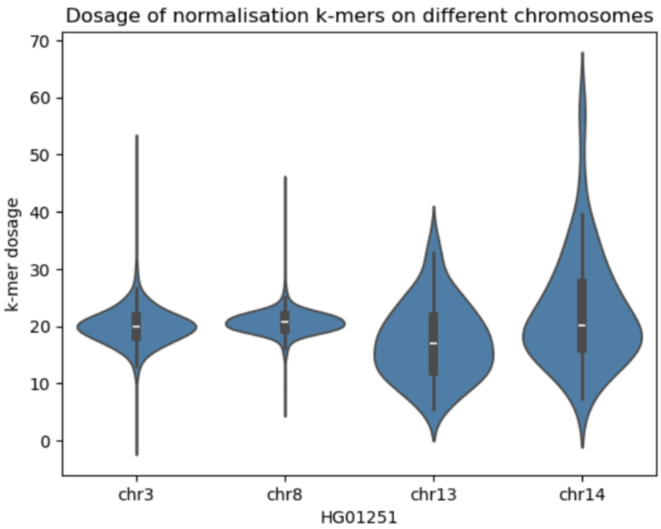


**Figure S12: Dosage distribution of normalization k-mers across selected chromosomes in a representative sample from the 1000 Genomes Project.**
The dosages of all normalization k-mers of chromosomes 3, 8, 13, and 14 were extracted from the De Bruijn graph of sample HG01251 and displayed as violin plots. Normalization k-mers showed a narrow dosage distribution on chromosomes 3 and 8, but a broader distribution on chromosomes 13 and 14.

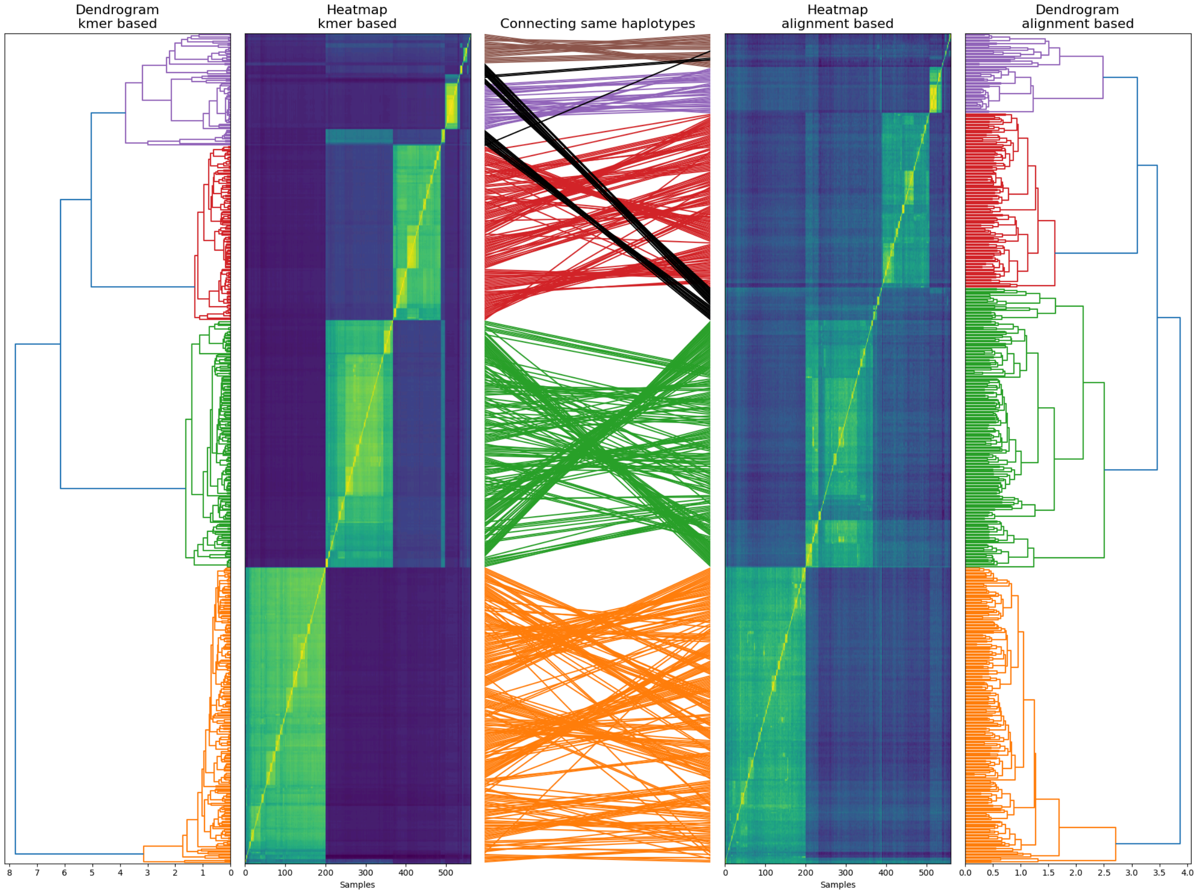


**Figure S13: Comparison of centromere haplotype clustering on chromosome 8 using *k*-mer-based and alignment-based similarity measures.**

This figure compares the clustering of 561 chromosome 8 centromere haplotypes using either *k*-mer-based (left) or alignment-based (right) pairwise similarity. From outer to inner panels: Outer panels: Dendrograms constructed from the respective pairwise similarity matrices. Distinct clusters are color-coded, and clusters with largely overlapping haplotype compositions between methods are assigned the same color. Inner panels: Heatmaps of pairwise similarity values between haplotypes (haplotypes are ordered identically along the x and y axes). Center panel: Correspondence plot connecting the positions of individual haplotypes in the two clustering results. Colored lines indicate that the haplotype is assigned to the same cluster (based on matched colors) in both methods, while black lines indicate different cluster assignments. This layout enables visual assessment of both the concordance and divergence between the two clustering approaches at the cluster and haplotype level.

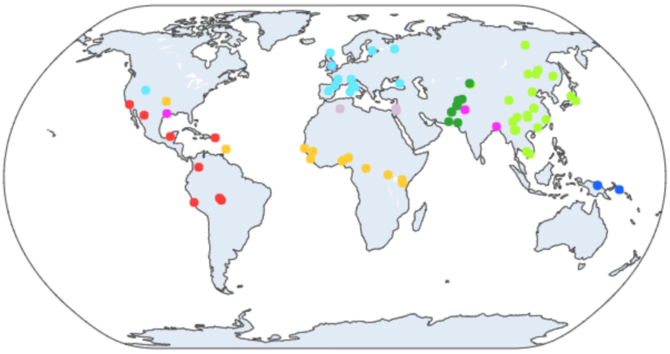

**Figure S14: Geographic distribution of populations from the 1000 Genomes Project and the Human Genome Diversity Project.**
Points indicate the reported sampling locations or population-associated geographic coordinates for each population and are plotted on a world map. Populations are colored according to their continental population.

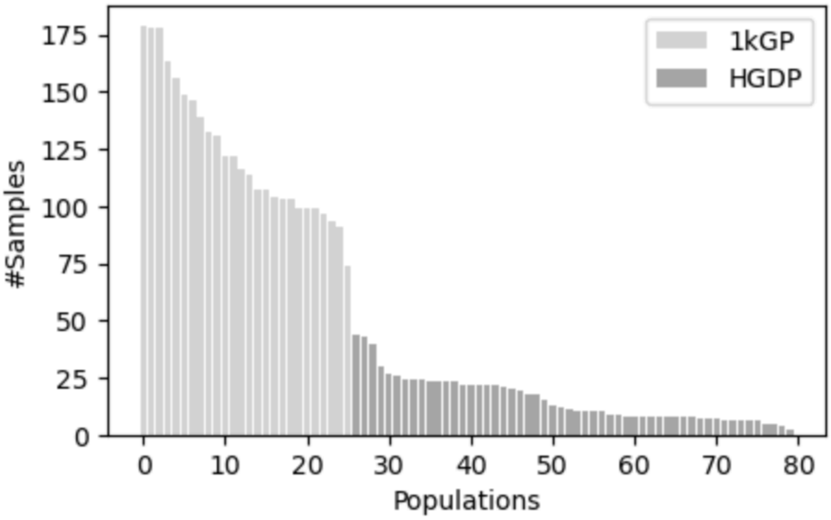


**Figure S15: Number of sequenced samples for the different populations of the 1000 Genomes (1kGP) and Human Genome Diversity Project (HGDP).**

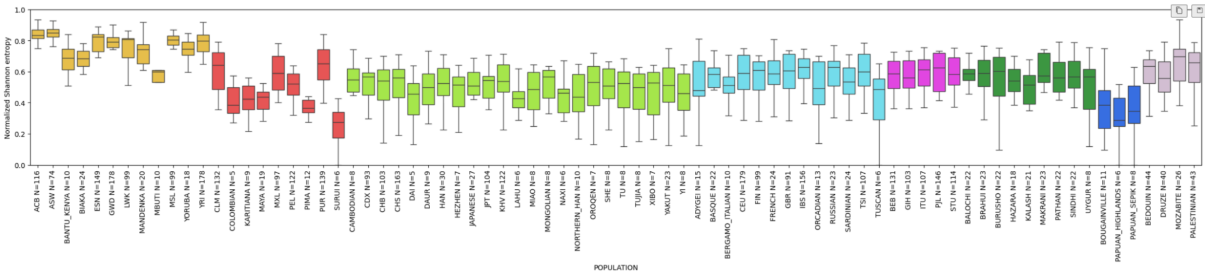

**Figure S16: Population-level diversity of centromere architectures.**

Distribution of centromere architecture diversity across populations, measured using normalized Shannon entropy. For each population, the boxplot shows chromosome-wise diversity values, with one value per chromosome calculated from centromere architecture frequencies within that superpopulation. Populations are grouped and colored according to their continental population: African (AFR), American (AMR), East Asian (EAS), European (EUR), South Asian (SAS), Central/South Asian (CSA), Oceanian (OCE), and Middle Eastern (MEE).

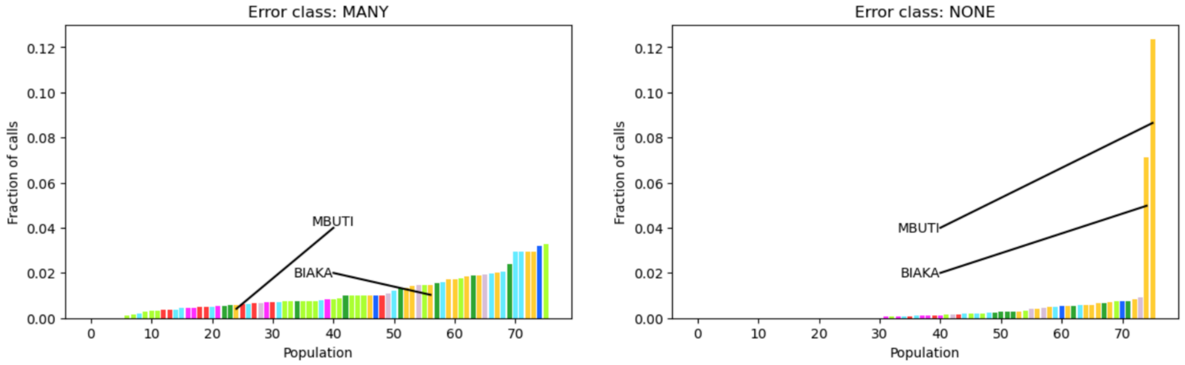


**Figure S17: Fraction of centromeres with three (left) or no (right) called centromere architectures stratified by population.**

The two outlier populations MBUTI and BIAKA with a high fraction of centromere without any call are labeled. Each population is colored according to the associated continental population.

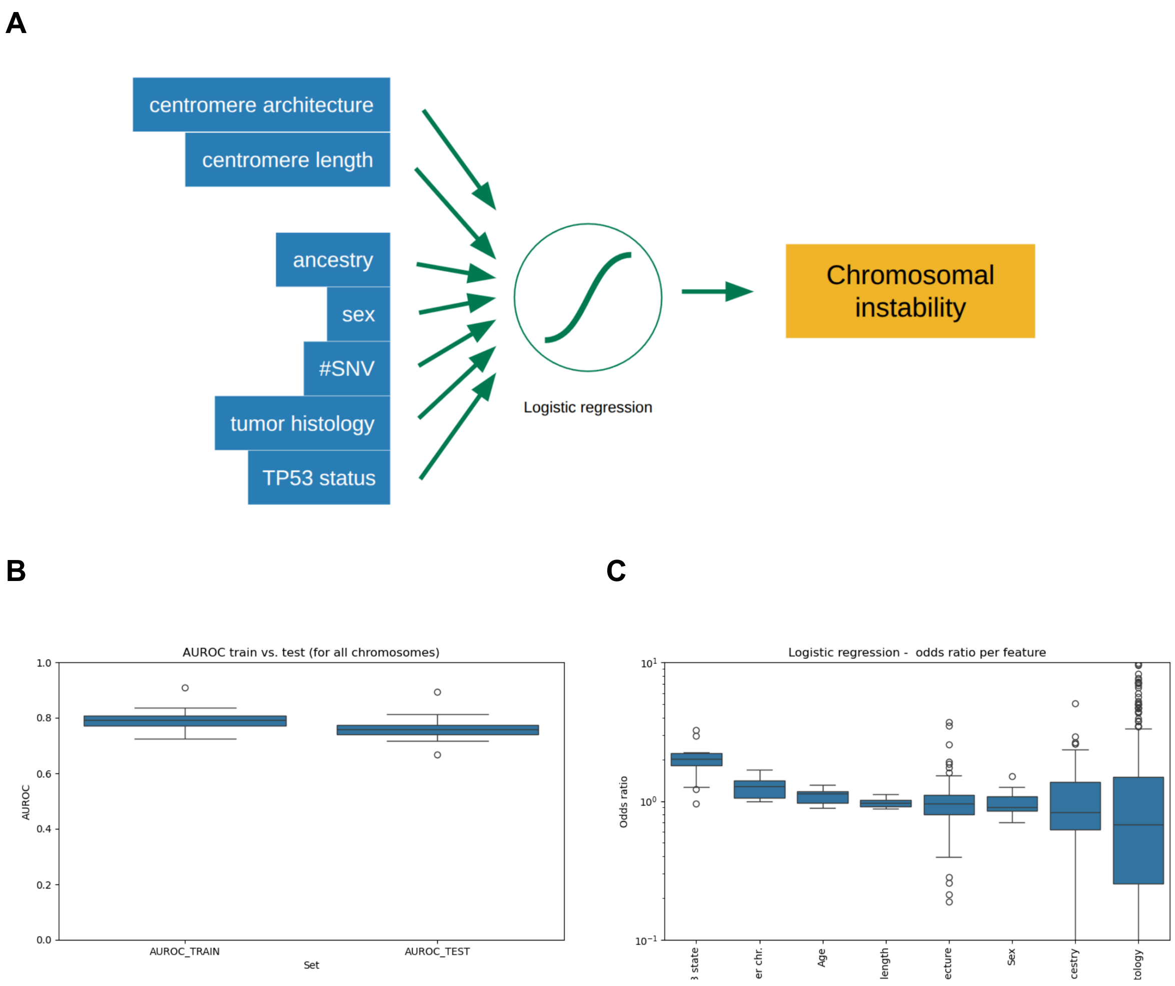


**Figure S18: Prediction of chromosomal instability from cancer genomics and metadata using logistic regression modeling.**

**(A)** Chromosomal instability (CIN) was modeled using logistic regression based on TP53 status, the number of SNVs per chromosome, tumor histology, sex, genetic ancestry, and inferred centromere characteristics, including architecture and length. **(B)** AUROC scores across cross-validation folds. **(C)** Distribution of feature odds ratios. Odds ratios were averaged across cross-validation folds; values greater than 1 indicate increased odds of CIN, values below 1 indicate decreased odds of CIN.

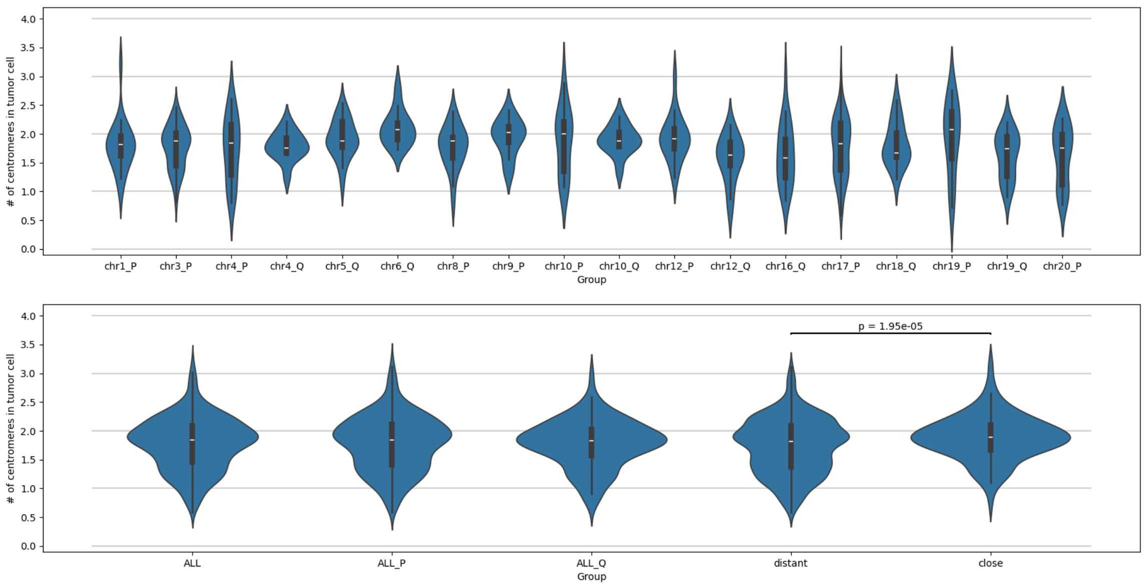


**Figure S19: Number of centromeres per tumor cell in chromosome arm-level loss events, stratified across multiple groups.**

Events are shown for all arm-level losses on individual chromosomal arms (top, e.g. chr1_P for p-arm loss in chr1) as well as for all (ALL), all p-arm (ALL_P), and all q-arm (ALL_Q) losses combined (bottom). Chromosomal arm losses are further grouped according to the position of the CDR relative to the lost arm, classified as either close ("close") or distant ("distant") and these groups were compared using a two-sample Kolmogorov–Smirnov test. The close group contains configurations where the CDR is on the side of the lost arm: p arm losses of chr1, chr9, chr12 and chr19 and q arm losses of chr4, chr5, chr6, chr10 and chr18.

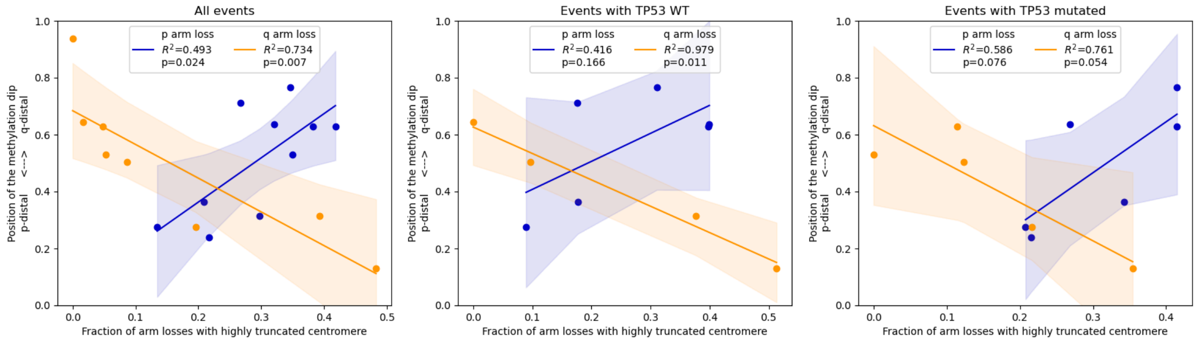


**Figure S20: Correlation between the fraction of strongly truncated centromeres in arm-loss events and the relative CDR position, stratified by TP53 mutation state.**

The fraction of strongly truncated centromeres in arm-loss events was determined using Gaussian mixture modeling based on inferred centromere numbers per tumor cell and plotted against the mean CDR position inside the HOR array. Analyses are shown for all samples combined (left), samples with wild-type TP53 only (middle), and samples with TP53 mutations only (right). Regression fits with confidence intervals, R² values, and p-values are shown.
